# Chlamydomonas mutant *hpm91* lacking PGR5 is a scalable and valuable strain for algal hydrogen (H_2_) production

**DOI:** 10.1101/2022.02.23.481610

**Authors:** Peng Liu, De-Min Ye, Mei Chen, Jin Zhang, Xia-He Huang, Li-Li Shen, Ke-Ke Xia, Xiao-Jing Xu, Yong-Chao Xu, Ya-Long Guo, Ying-Chun Wang, Fang Huang

## Abstract

Clean and sustainable H_2_ production is essential toward a carbon-neutral world. H_2_ generation by *Chlamydomonas reinhardtii* is an attractive approach for solar-H_2_ from H_2_O. However, it is currently not scalable because of lacking ideal strains. Here, we explore *hpm91*, a previously reported PGR5-deletion mutant with remarkable H_2_ production, that possesses numerous valuable attributes towards large-scale application and in-depth study issues. We show that *hpm91* is at least 100-fold scalable (upto 10 liter) with H_2_ collection sustained for averagely 26 days and 7287 ml H_2_/10L-HPBR. Also, *hpm91* is robust and active over the period of sulfur-deprived H_2_ production, most likely due to decreased intracellular ROS relative to wild type. Moreover, quantitative proteomic analysis revealed its features in photosynthetic antenna, primary metabolic pathways and anti-ROS responses. Together with success of new high-H_2_-production strains derived from *hpm91*, we highlight that *hpm91* is a potent strain toward basic and applied research of algal-H_2_ photoproduction.

## Introduction

Hydrogen (H_2_) derived from H_2_O is a clean and versatile energy carrier that could be obtained sustainably through sunlight-driven chemical and biological means such as photocatalysis and algal photoproduction (Bayro-Kaiser and Nelson, 2017; Nishiyama et al., 2021). Under anaerobic condition, H_2_ photoproduction occurs in microalgae such as *Chlamydomonas reinhardtii* (henceforth referred to as *Chlamydomonas*) virtually via water-oxidation in visible light (Gfeller and Gibbs, 1984; Greenbaum, 1988). H_2_ formation is through the catalytic activity of (Fe-Fe) hydrogenases (Forestier et al., 2003) induced under anoxia, using mainly photosynthetic electrons on the acceptor side of photosystem I (PSI) in reduction of protons into H_2_ (Forestier et al., 2003; Ghirardi, 2015). As a result, renewable H_2_ production is achieved in the organisms using energy input from solar irradiation and electrons derived from photosynthetic water-splitting reaction. In wild-type Chlamydomonas strains, however, H_2_ photoproduction is only transient in anoxia or sustains for several days under sulfur-deprived anaerobic condition (Ghirardi, 2015; Kruse et al., 2005; Melis et al., 2000). This is a major obstacle to make this bio-system economically viable for solar-H_2_ production.

Using various genetic strategies, dozens of Chlamydomonas mutants with increased H_2_ production have been isolated over the last decades. Several of them are reasonably well documented such as the state transition mutant *(stm6)*, truncated light-harvesting antenna mutant *(tla1)*, D1 mutant (Kosourov et al., 2011; Kruse et al., 2005; Scoma et al., 2012; Volgusheva et al., 2013), and the *pgr* mutants named as *pgrl1* (proton gradient regulation like 1), *pgr5* (proton gradient regulation 5) and *hpm91* (Chen et al., 2016; 2019; Steinbeck et al., 2015; Tolleter et al., 2011). Of those, the *pgr* mutants are of considerable interest because their target gene products are intimately involved in the PGR5-dependent branch of photosynthetic cyclic electron flow (CEF), which is the major route of CEF (Chen et al., 2016; Schwenkert et al., 2022; Steinbeck et al., 2015; Takahashi et al., 2013; Tolleter et al., 2011). While a negative correlation between the CEF and H_2_ photoproduction was initially indicated in the *stm6* mutant deficient in *MOC1* gene which encodes an assembly factor of the mitochondrial respiratory chain (Schonfeld et al., 2004), more direct experimental evidence was obtained via analysis of *pgr (pgrl1, pgr5, hpm91)* as well as *fnr* (ferredoxin-NADPH reductase) mutants (Sun et al., 2013; Yacoby et al., 2011). PGRL1 and FNR have been identified in the CEF supercomlex (Iwai et al., 2010) and PGR5 are known being a regulatory protein involved in the CEF branch (Munekage et al., 2002; Suorsa et al., 2012), for which the mechanism of action is currently not yet clear (Schwenkert et al., 2022).

Most remarkably, the *hpm91* mutant sustains H_2_ photoproduction for 25 days (Chen et al., 2016). Although an inverse relationship was revealed between PGR5 levels and H_2_ production, and the prolonged H_2_ evolution was largely attributed to the enhanced anti-ROS capability protecting the photosynthetic electron transport chain from photooxidative damage (Chen et al., 2016), questions arise as i) whether the phenotype of *hpm91* is stable in large scale setting-ups? ii) how intracellular ROS content of *hpm91* is altered during sulfur-deprived H_2_ photoproduction? iii) what are the proteomic characteristics of *hpm91* under such conditions. We report here the performance of *hpm91* in a large scale (upto 10 liter) H_2_-photobioreactor (HPBR) systems. We also describe new findings of *hpm91* at the level of systems biology associated with the intial analytical scale HPBR (100 ml). We highlight the *hpm91* as a valuble algal strain not only for basic research of understanding the molecular mechanisms of H_2_ photoproduction but also for development of economically viable solar-powered H_2_ production systems.

## Results and Discussion

### H_2_ production of *hpm91* was large-scalable

Previous experimental results and literature survey revealed top ranking of *hpm91* among Chlamydomonas mutants with enhanced H_2_ photoproduction under sulfur-deprived condition (Chen et al., 2016; 2019). To determine its potential towards application, we carried out scaling-up and gas-handling experiments in the laboratory using *hpm91* (Fig. 1; Movie S1). Initially, the experiments were performed with H_2_-photobioreactors (HPBR) of scaling-up 10-, 20-, and 30-times (from 100 ml) using chlorophyll concentration of 25 µg/ml as previously (Chen et al., 2016; 2019). We found the highest yield of H_2_ from the 3L-HPBR as well as a positive correlation between the yield and size of HPBR (Fig. 1A, B). These results encouraged us to extend the scaling-up directly to 10L-HPBR (headspace 1700 ml, light path 22 cm, illuminated on 3 sides) and to investigate its H_2_ production profiles. To reduce potential shading effects (Hemschemeier et al., 2009; Kosourov et al., 2002) and low-light stress on algal cells, we reasonably reduced initial chlorophyll concentration (ICC) to 20 µg/ml of cell suspension and cultured under increased light irradiance (130 µE m^-2^ s^-1^) for H_2_ production.

**Fig. 1.**
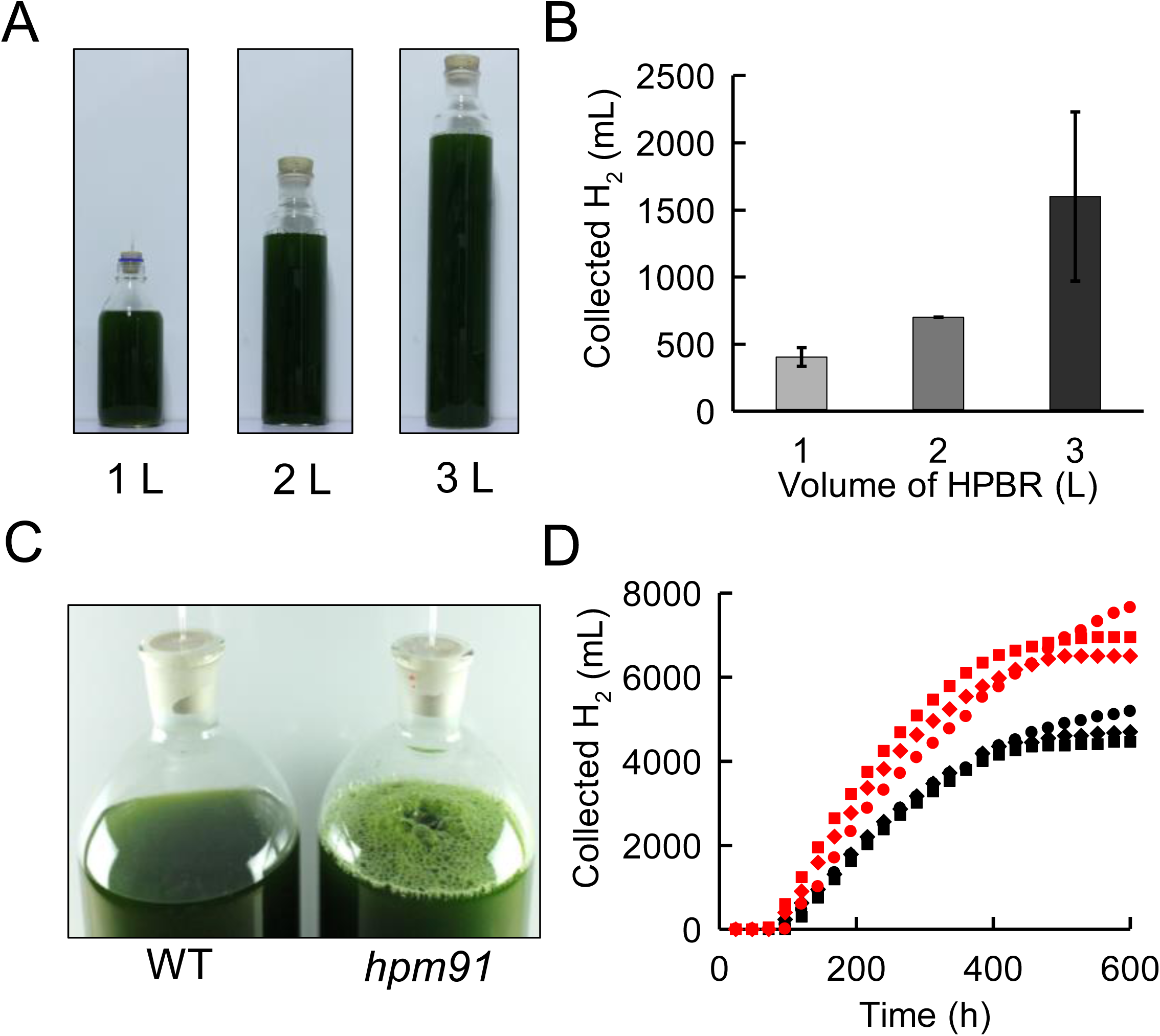
H_2_ production of *hpm91* in 1 to 10 L-HPBR under sulfur deprivation. *(A)* Photograph of 1 to 3 L-HPBR (optical path 10 cm, 100 µE m^-2^ s^-1^.) taken at 10 day of sulfur deprivation. *(B)* Correlation between H_2_ output and size of HPBR. Standard deviations were estimated from three biological replicates. *(C)* Photograph of wild type and *hpm91* in 10L-HPBR taken at 4 day of sulfur deprivation. *(D)* Comparison of H_2_ output profiles of *hpm91* in 10L-HPBR under light intensity of 130 (in black) and 230 µE m^-2^ s^-1^ (in red). The data are from three independent experiments.

Fig. 1C compares H_2_ photoproduction of *hpm91* and wild type in 10L-HPBR under identical cultural condition. In contrast to wild type, which generated few gas bubbles with collectable H_2_ quantity negligible, we observed a bulk of H_2_ bubbles in *hpm91* after 3-4 days of sulfur deprivation with the gas collectable for up to 33 days (Fig. 1C). To confirm these results, experiments were repeated twice using different batch of algal cells. The data shows that continuous H_2_ collection from *hpm91* could be achieved at an average of 28 days (*SI Appendix*, Table S1), which was the longest duration of H_2_ production under sulfur deprivation reported thus far for Chlamydomonas. These results promoted us to explore more possibilities to enhance H_2_ production of *hpm91* in 10L-HPBR.

Indeed, further increase of light intensity to 230 µE m^-2^ s^-1^ apparently enhanced H_2_-producing capability of *hpm91*. Compared to that under lower light (130 µE m^-2^ s^- 1^), output of H_2_ increased up to 57.9% (*SI Appendix*, Table S1). Because H_2_ output stopped in most cases around 25 day of sulfur deprivation, we then compared its H_2_ evolution profiles within the period. As shown in Fig. 1D, the gereral kinetic pattern was largely similar but higher rate and extended linear increasing period (480 h *vs* 420 h) of H_2_ production was observed for those cultured under increased light. An average H_2_ output reached to 7287 ml/10L-HPBR, which was 50.7% higher than that under lower light (*SI Appendix*, Table S1). Further experiments verified its essentially pure H_2_ output which enabled us to make a H_2_ fuel cell-powered toycar drive using ambient air directly (Movie S1). These results allowed us to address the value of *hpm91* in not only basic research but also in advancing bio-H_2_ photoproduction technology.

### Decreased intracellular ROS in *hpm91*

Based on visual comparison of cell suspension, cell growth of *hpm91* was always better than wild type in both small (100 ml) and large HPBR (1 to 10L) especially during prolonged period of H_2_ production. To understand the basis of cell biology, we investigated changes of cell morphology and viability in *hpm91* committed to H_2_ production. Because the small HPBR system was experimentally more efficient, the system was of choice for comparative studies between wild type and *hpm91* during 120 h of H_2_ production (Fig. 2).

**Fig. 2.**
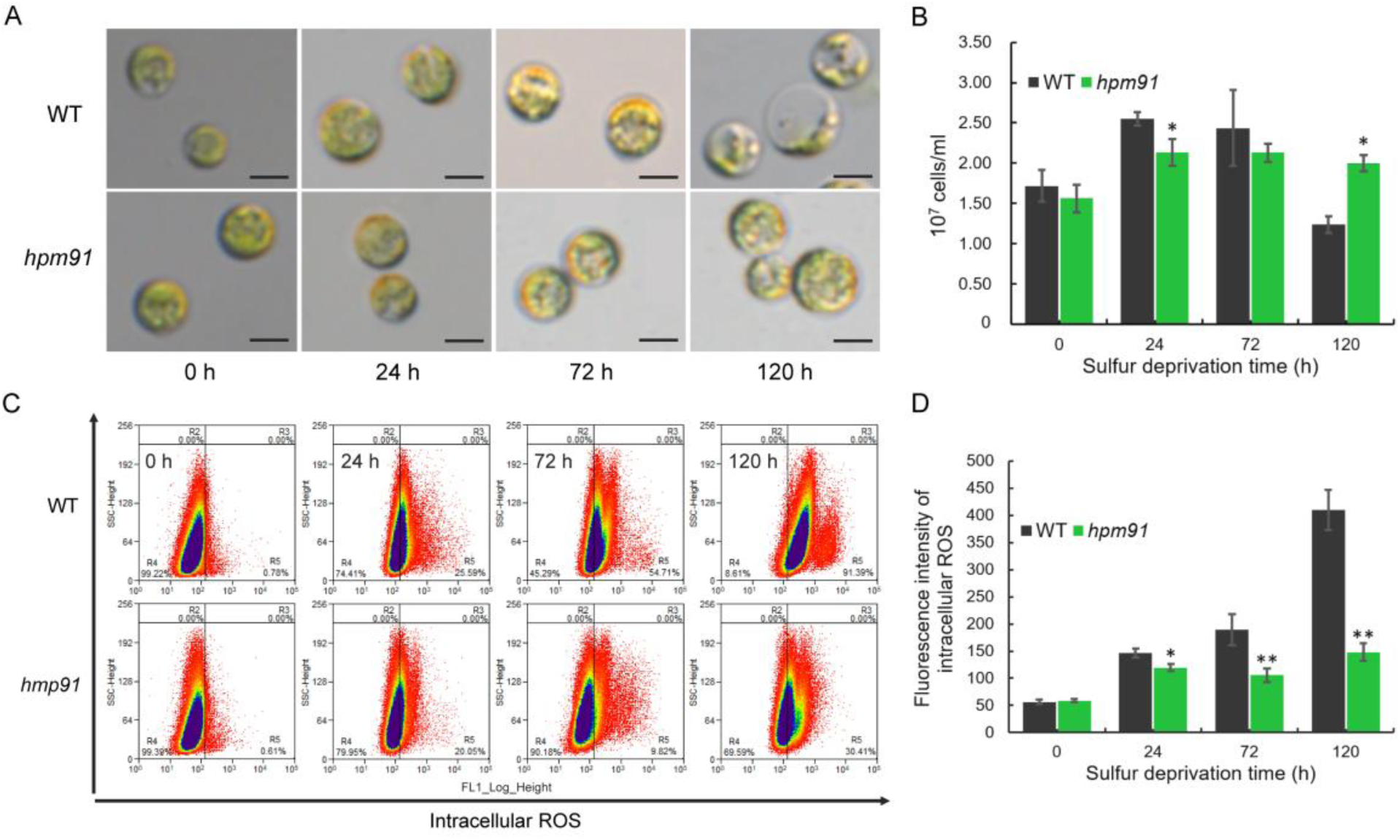
Comparison of intracellular ROS contents of *hpm91* and wild type during 120 h of sulfur-deprived H_2_ production process. *(A)* and *(B)* Cell morphology and proliferation of the two strains. *(C)* and *(D)* Scatter diagram and quantification of intracellular ROS contents in the two strains. Sample preparation was performed under anoxia and ROS levels were determined using a MoFlo XDP high-speed flow cytometer (Beckman-Coulter, Inc. USA) by following the manufacturer’s instructions. Average fluorescence intensity of DCF from 3×10^5^ cells was recorded and data acquisition and analysis were carried out using Summit 5.2 software. Experiments were repeated three times with similar results. * and ** refer to p-values <0.05 and <0.01 in Student’s t-test, respectively.

Fig. 2A shows that cells of wild type became largely translucent whereas no such changes were observed in *hpm91*. This remarkable morphological alteration of wild-type cells was highly similar to earlier observations which was mainly attributed to the substantial loss of endogenous starch and a declined cell viability under such condition (Zhang et al., 2002). Also, culture density of *hpm91* was more stable after 24 h and remained significantly higher than wild type at the end of the measurements (120 h) (Fig. 2B). Moreover, we found decreased portion of dead cells in the culture of *hpm91* relative to wild type (*SI Appendix*, Fig. S1). These results are clear indications of robustness of *hpm91* cells towards sulfur-deprived H_2_ production.

To further elucidate this, we measured intracellular ROS contents of *hpm91* and wild type under such conditions (Fig. 2C, D). ROS levels were determined using a MoFlo XDP high-speed flow cytometer (Beckman-Coulter, Inc. USA) by following the manufacturer’s instructions. Sample preparation was performed under anoxia with addition of glucose oxidase, catalase and glucose just before flow cytometric measurements (Chen et al., 2019). Average fluorescence intensity of DCF from 3×10^5^ cells was recorded and data acquisition and analysis were carried out using Summit 5.2 software (Beckman-Coulter, Inc. USA) (Fig. 2C). Our data shows that intracellular level of ROS in *hpm91* remained significantly lower than wild type during H_2_ prodution. At 120 h, the amount of ROS in *hpm91* was only 1/3 of that in wild type (Fig. 2D). These *in vivo* experimental data are in line with our previous results, showing increased activity of ROS-scvenging enzymes in *hpm91* relative to wild type (Chen et al., 2016). Thus, the better cell viability of *hpm91* could be largely attributed to its decreased level of ROS. A question raises how this is elicited in the organism under such conditions.

### Overview of proteome changes in wild type and *hpm91* under sulfur deprivation

Earlier work revealed preliminary metabolic interplay in wild type and *pgr* mutants of Chlamydomonas during sulfur-deprived H_2_ photoproduction (Chen et al., 2010; Melis et al., 2000; Steinbeck et al., 2015; Tolleter et al., 2011; Zhang et al., 2002). To understand molecular mechanism of H_2_ metabolism regulated by PGR5, we then carried out iTRAQ quantitative proteomics of *hpm91* and wild-type cells during 120 h of H_2_ production. As presented in Fig. 3A, samples were taken at four different time points (0, 24, 72 and 120 h) and total proteins were extracted for protein identification and quantification. A total of 3798 proteins with quantitative data were confidently identified (FDR<1%) as listed in (Dataset S1). Identification was mostly based on minimum of two peptide-hits per protein but considering several proteins such as iron hydrogenase Hyd3 were probably involved H_2_ metabolism and numerous functionally important small-sized and/or membrane proteins may possess only one identifiable tryptic peptide in MS analysis, those identified with one-peptide hit were also reasonably included (Dataset S1).

**Fig. 3.**
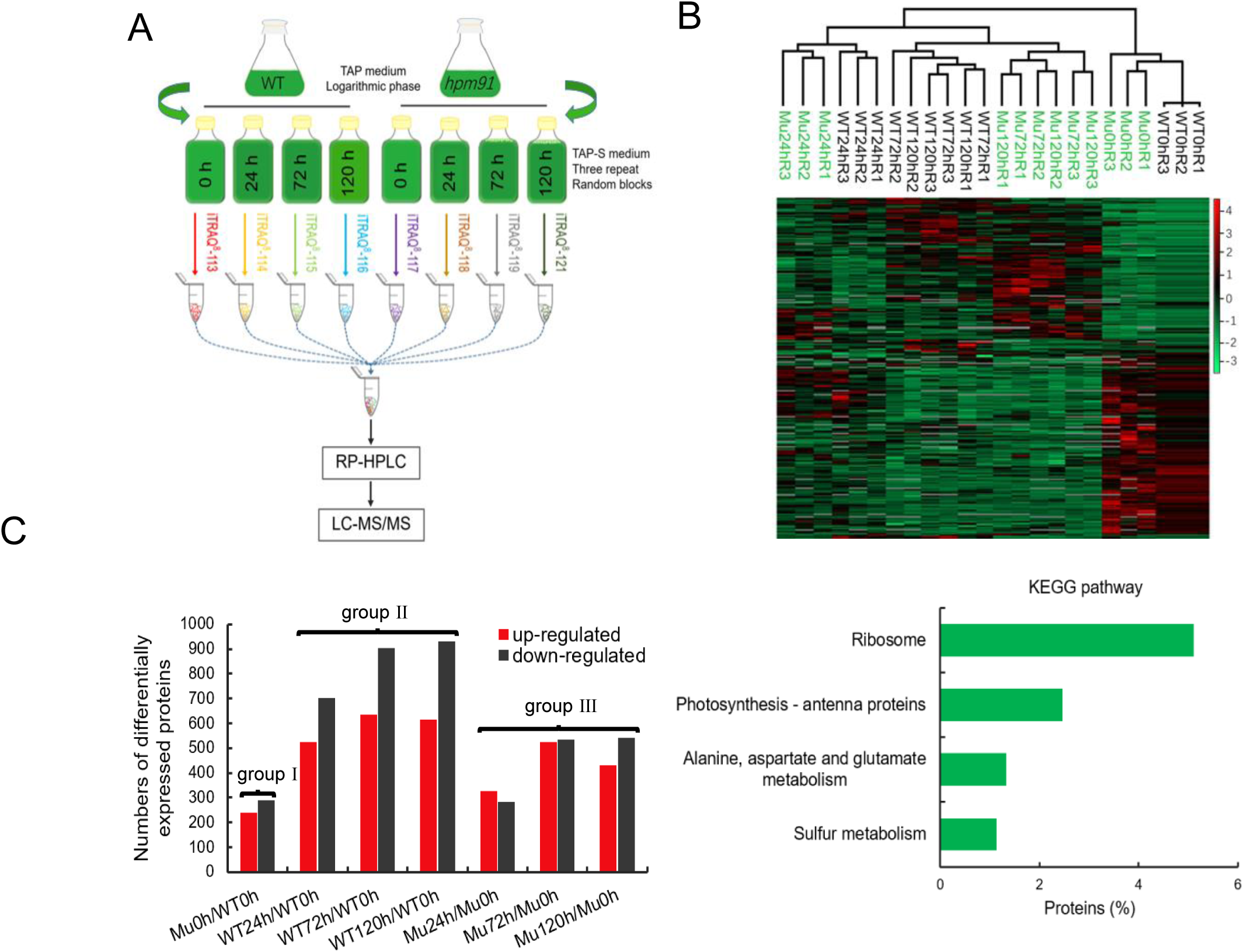
Overview of iTRAQ proteomics of *hpm91* and wild type during 120 h of sulfur-deprived H_2_ production. *(A)* Schematic presentation of iTRAQ experimental design. *(B)* Hierarchy clustering analysis showing high reproducibility of protein quantitation. *(C)* Number of differential expressed proteins in *hpm91* and wild type using a cutoff of 1.2-fold change with significance (p<0.05). *(D)* KEGG analysis of group I proteins in *(C)* showing major proteome changes caused by PGR5 deletion in Chlamydomonas. DAVID Bioinformatic Resources 6.8 (https://david.ncifcrf.gov/summary.jsp) was used.

Reproducibility of protein quantitation was verified by hierarchical clustering analysis (Perseus _1.6.0.7) of these proteins obtained with three biological replicates (Fig. 3B). To confirm functional impairment of *hpm91* in cyclic electron transfer (CEF), we compared the CEF rates in both strains. The data showed that CEF of *hpm91* was significantly decreased relative to wild type (*SI Appendix*, Fig. S2A). To be more certain with the genetic background of the strains, we performed high-throughput genomic sequencing for wild type (CC400, 137c) and the *pgr5* mutants (*hpm91, pgr5*). The data was deposited at the CNGBdb database (Chen et al., 2020; Guo et al., 2020) (https://db.cngb.org/search; accession No. CNP0002674). Further analysis of reads coverage validates previous mutation mapping (Chen et al., 2016; Johnson et al., 2014) and showed large deletions in PGR5-containing region of *hpm91* and *pgr5* mutants *(SI Appendix*, Fig. S2B). Together with the immuno-blot results showing no detectable PGR5 but presence of PGRL1 and FNR in *hpm91* (Chen et al., 2016), we clarify that the impaired CEF of *hpm91* is arributed to loss of PGR5. Differentially expressed proteins were determined using a cutoff of 1.2-fold change with significance (p<0.05), leading to three groups consisting of 529 (group I), 2229 (group II) and 1350 (group III) proteins (Fig. 3C) listed in (Datasets S2-S4). These correspond to three comparisons, *i*.*e. hpm91* at 0 h *vs* wild-type at 0 h, any time of- wild type *vs* wild-type at 0 h, and any time of-*hpm91 vs hpm91* at 0 h, representing differentially expressed proteins caused by deletion of PGR5 and by sulfur-deprived anoxia in wild type and *hpm91*, respectively.

To understand functional significance of the differentially expressed proteins in each group, KEGG and gene ontology (GO) analysis was performed using DAVID Bioinformatic Resources 6.8 (https://david.ncifcrf.gov/summary.jsp), yielding 4 enriched KEGG pathways for group I (Fig. 3D), 30 and 29 enriched biological process (GOPB) for the later two groups, respectively (Fig. 4A, Datasets S5 and S6). Because numerous proteins in the latter two enrichments were multiply or/and with error assignments, we reasonably delineated them into 8 and 9 major groups as (Wang et al., 2012) and shown in (*SI Appendix*, Table S2 and S3), respectively.

**Fig. 4.**
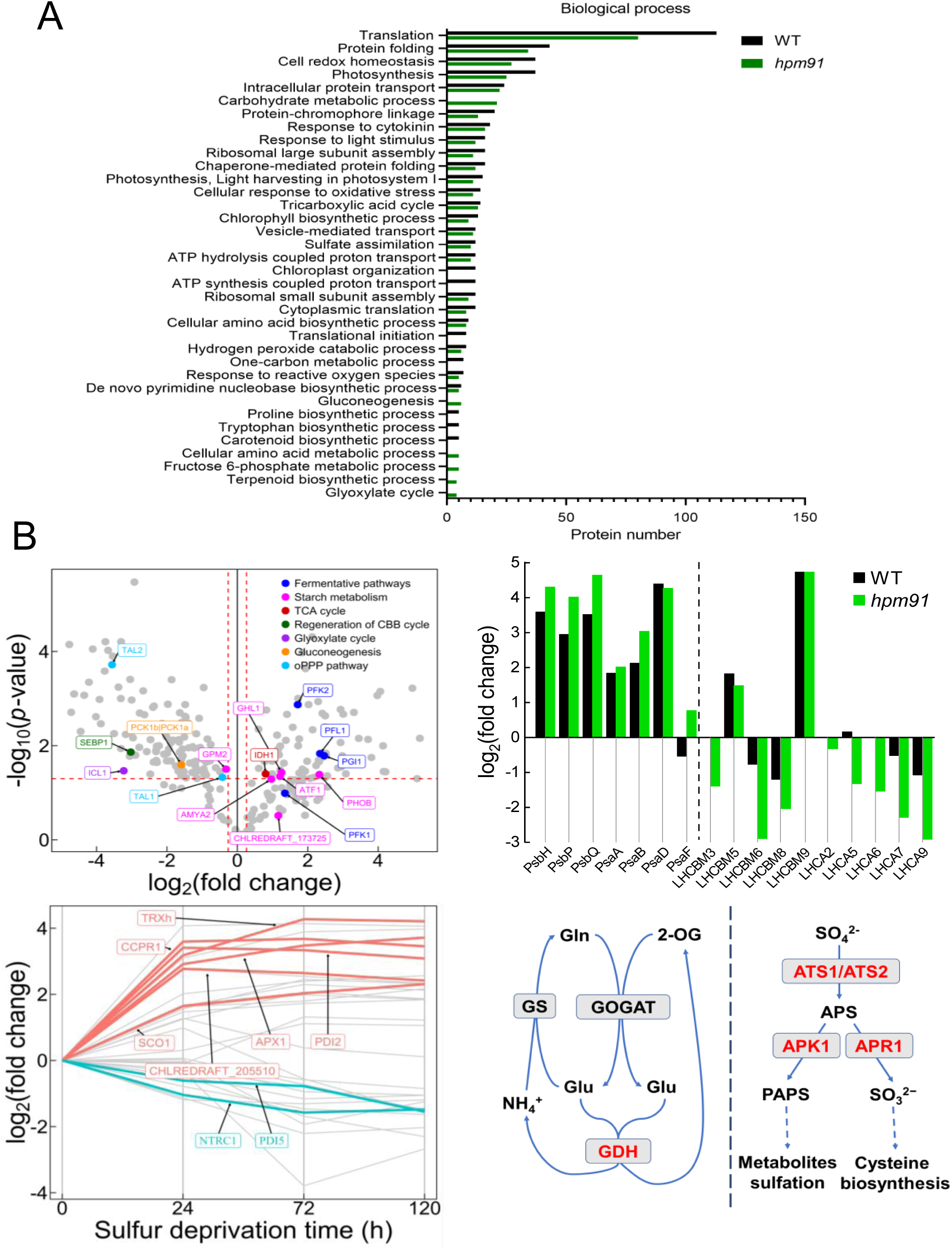
Proteomic characteristics of *hpm91* during sustained H_2_ production. *(A)* Comparison of GOBP enrichmemts in *hpm91* and wild type during 120 h of sulfur seprivation. *(B)* Major proteome feature of *hpm91* under sulfur-deprived condition. Volcano plot shows major changes in carbon metabolism of *hpm91* during 120 h of sulfur-deprivation *(upleft panel)*. Comparison of average fold-change of photosynthetic proteins in *hpm91* and wild type at 120 h sulfur-deprivation *(upright panel)*. Dynamic changes of redox proteins in *hpm91* during 120 of sulfur deprivation *(downleft panel)*. Schematic illustration of N- and S-metabolic features in *hpm91* under sulfur-deprived condition *(downright panel)*.

It can be seen in Table 1, deletion of PGR5 caused significant changes in four pathways in Chlamydomonas. Compared to wild type, all the ribosomal proteins and most of those related to nitrogen metablism were higher-expressed in *hpm91*. The latter may implicate enhanced nitrogen metabolism in *hpm91*. To test this possibility, we compared phenotype of the two strains under N-starved stress condition. The experimental data showed that both cell growth and photosynthetic capability of *hpm91* was indeed better than wild type (*SI Appendix*, Fig. S2C), revealing another impact of PGR5 on chloroplast biology. More interestingly, we found all the photosynthetic antenna proteins except for LHCA3 was lower-expressed in *hpm91*. Because those account for more than 50% of LHCI and LHCII proteins in Chlamydomonas (Shen et al., 2019; Su et al., 2019; Suga et al., 2019), our finding of their reduced levels could be an indication of a smaller photosynthetic antenna in *hpm91* than wild type. Notably, among the proteins involved in sulfur metabolism only APS reductase APR1 encoded by *APR1*/*MET16* (GutierrezMarcos et al., 1996; Setya et al., 1996), known to be involved in sulfur-starvation response (Ravina et al., 2002; Zhang et al., 2004) was higher-expressed in *hpm91* (Table 1).

**Table 1.**
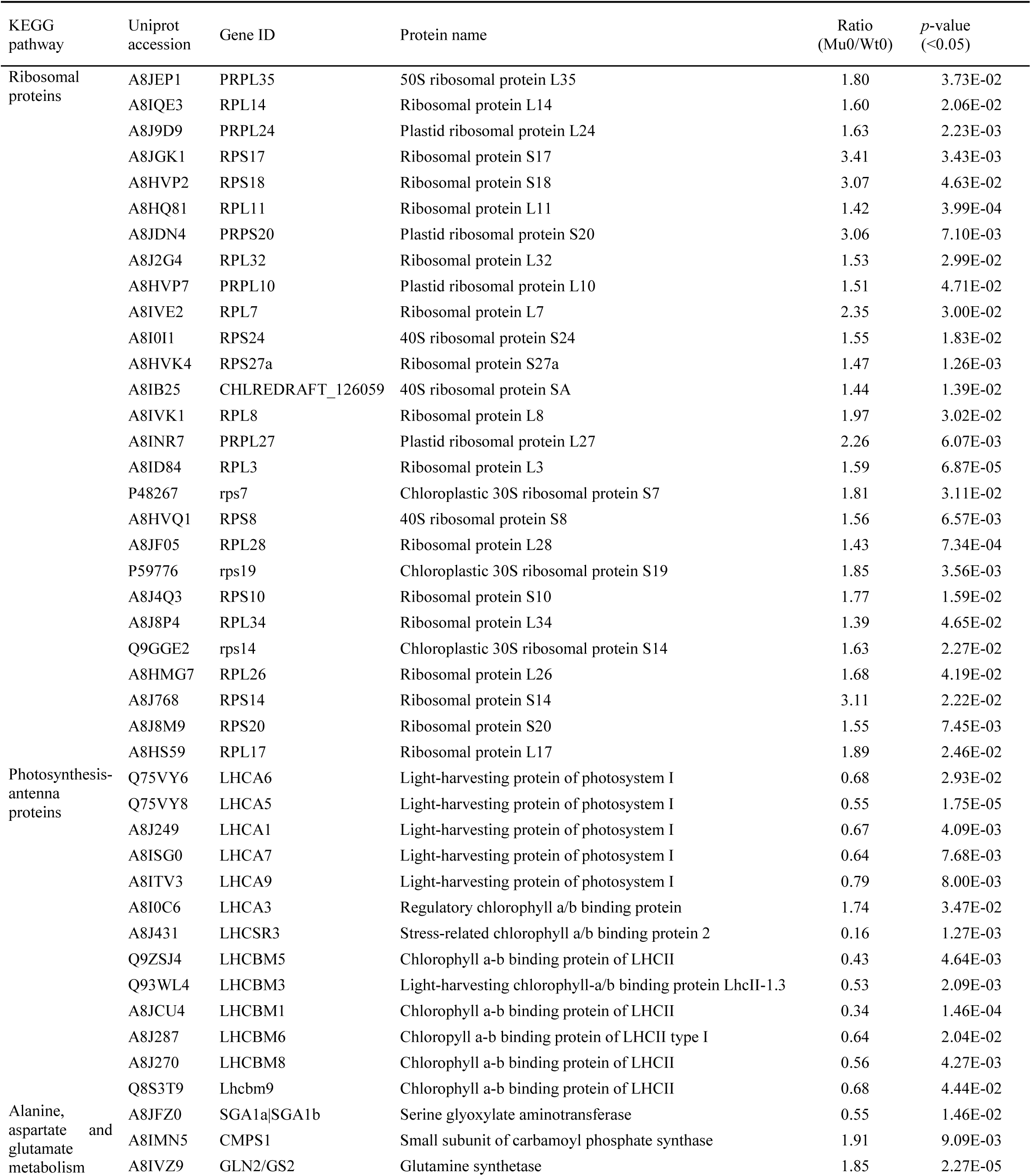
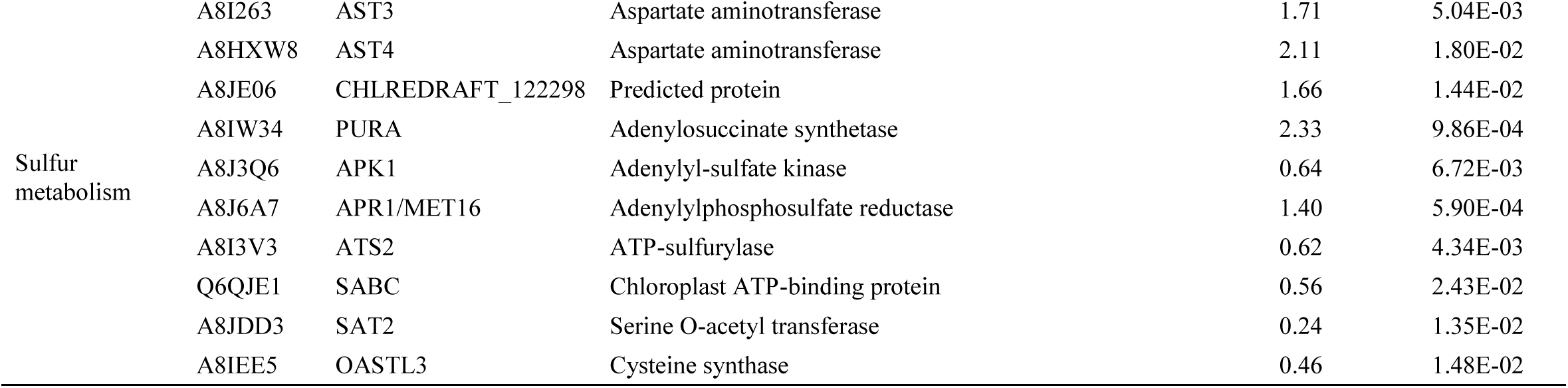
List of proteins corresponding to Fig. 3D.

Comparison of the results in Table S2 and S3 revealed similarities in 5 functional groups, *i*.*e*. translation, protein folding, intracelluar protein trafficking, response to cytokinin, and ATP hydrolysis/production, reflected by similar change patterns of numerous overlapping proteins in the two strains. Differences were also revealed in 4 of those corresponding to carbon metabolism, photosynthetic antenna, cell redox homeostasis/anti-oxidative systems, as well as nitrogen- and sulfur metabolisms. These are the major proteomic characteristics of *hpm91* towards sustained H_2_ production as described/discussed below.

### Proteomic characteristics of *hpm91* during sustained H_2_ production

Apparently, loss of PGR5 in *hpm91* causes compromised primary carbon metabolism during sustained H_2_ production. As can be seen in Fig. 4A, 6 biological processes were only enriched in *hpm91* during H_2_ production. These were ‘carbohydrate metabolism’, ‘gluconeogenesis’, ‘fructose 6-phosphate metabolic process’, ‘glyoxylate cycle’, ‘cellular amino acid metabolic process’ and ‘terpenoid biosynthetic process’ with the first one appeared within top-10 rankings, implicating that loss of PGR5 caused more profound alterations of primary carbon metabolism in *hpm91* than wild type during sulfur-deprived H_2_ production. Considering metabolic relevance in Chlamydomonas under anoxia (Yang et al., 2015), the group ‘tricarboxylic acid cycle’ was thus combined with the first 4 of them (*SI Appendix*, Table S3). While many of them overlapped with wild type, 16 proteins were exclusively revealed in *hpm91* during sustained H_2_ production. These include the key proteins in the regeneration pathway of Calvin-Benson-Bassham (CBB) cycle (SEBP1), glyoxylate cycle (ICL1), gluconeogenesis (PCK1b |PCK1a), oxidative PPP pathway (oPPP) (TAL1, TAL2), fermentative pathways (PFK1, PFK2, PGI1, PFL1), starch metabolism (AMYA2, AMY-like protein, GHL1, ATF1, PHOB, GPM2), and tricarboxylic acid cycle (IDH1), showing increased trends for the most proteins in the first three and decreased trends in the latter four pathways, respectively (Fig. 4B, upleft pannel). It has been earlier reported by Philipps et al. (Philipps et al., 2011) that loss of PFL1 decreased H_2_ photoproduction. Our finding of increased amount of PFL1 in *hpm91* was in line with this and may suggest a partial contribution of PFL1 to its enhanced H_2_ production. Also, we found alpha-amylases (encoded by AMYA2, CHLREDRAFT_173725) and glucosamine-fructose-6-phosphate aminotransferase (ATF1) as well as phosphorylase PHOB increased at an average of 1.86 to 5.02-fold in *hpm91* during H_2_ production (*SI Appendix*, Table S3). These enzymes are known to be essential for starch metabolism (Weigelt et al., 2009). While accumulation of PHOB could be correlated to the marked increase of starch contents in *hpm91* (Chen et al., 2016), elevated level of the amylases was somehow intriguing because starch breakdown in *hpm91* was significantly less than wild type (*SI Appendix*, Table S3; Chen et al., 2016), excluding the major contribution of ‘ indirect pathway’ on the prolonged H_2_ production phenotype of *hpm91*. Moreover, our data revealed down-regulated key enzymes (PRK1, FBP1, SEBP1) in or related to CBB cycle in *hpm91*, suggesting that the main route of carbon fixation was largely repressed towards H_2_ production. It is possible that the increased levels of amylases as well as GHL1 and ATF1 in *hpm91* is to yield various intermediates satisfying increased carbon demand under anoxic conditions (Weigelt et al., 2009). Interestingly, we observed a large variation of the key enzymes in TCA cycle of *hpm91*, such as malate dehydrogenase MDH4 and subunits of succinate dehydrogenase (SDH1, SDH2, SDH4) (*SI Appendix*, Table S3). Because rate of TCA flux is in general set by concentrations of substrates and intermediates that provide optimal levels of ATP and NADH rather than only by the abundance of the enzymes, the observed variations could be an indication of dynamic energetic status in *hpm91* during sulfur-deprived H_2_ production.

Also, *hpm91* is characteristic of more pronounced accumulation of photosynthetic core proteins with more reduced PSI antenna during sulfur-deprived H_2_ production. As shown in Table S3, nearly 50% of the PSII and PSI proteins were accumulated in *hpm91* under sulfur deprivation. These include PsbH, PsbP, PsbQ, PsaA, PsaB, PsaD and PsaF with average values of fold-increase within 3.1 to 25.2. Because H_2_ evolution profile of wild type and *hpm91* was mostly distinct at 120 h of sulfur deprivation (Chen et al., 2019), the change-fold of this time point was compared between the two strains (Fig. 4B, upright pannel). Compared to wild type, fold-increase values of the three PSII proteins was about 2-times larger in *hpm91*. Considering that PsbP and PsbQ are essential to the maintenance of water-splitting reaction (Shen, 2015) and PsbH is crucial for stable assembly and optimal function of PSII (Trosch et al., 2018; Umena et al., 2011), their greater increase in *hpm91* may partially explain its significantly higher residual PSII activity during prolonged H_2_ production (Chen et al., 2019). Regarding to LHCII proteins, we noted that decrease of LHCBM3, LHCBM6, LHCBM8 were more pronounced in *hpm91* under such conditions (Fig. 4B, upright panel).

In PSI, we found that while the fold-increase values of PSI core subunits of *hpm91* were either higher (PsaB and PsaF) or comparable (PsaA, PsaD) to wild type, all the LHCA proteins displayed declined trends with greater values of decrease-fold in *hpm91* than wild type during H_2_ production process (Fig. 4B, upright panel). The results strongly suggest that *PGR5*-deficient *hpm91* mutant is an algal strain characteristic of small-sized PSI antenna under not only normal condition (Table 1) but also during sustained sulfur-deprived H_2_ production. It is generally known that, in wild type, transcriptional regulation of LHC genes plays a central role in antanna size adjustment. To test if this is true for *hpm91* mutant, we then carried out qRT-PCR analysis of the genes encoding the LHCA proteins. Our data showed that their mRNA levels were indeed down-regulated during H_2_ production (*SI Appendix*, Fig. S3). Taken together, we suggest that mutation of *PGR5* caused not only the impaired CEF (*SI Appendix*, Fig. S2A) and higher residual photosynthetic activity (Chen et al., 2016; 2019) but also significantly reduced PSI antenna (Fig. 4B), leading to its higher enficiency of light utilization than wild type towards H_2_ photoproduction (Kosourov et al., 2011).

Yet, deletion of PGR5 reinforced cell redox homeostasis and anti-oxidative systems in *hpm91* under sulfur deprivation. Based on comparison of Table S2 and S3, higher percentage of up-regulated proteins involved in cell redox homeostasis and/or anti-oxidative stress responses were found in *hpm91* than wild type during H_2_ production process. Although most of them overlapped with wild type, accumulation of several proteins in TRX superfamily, mitochondrial proteins SCO1 and cytochrome c peroxidase CCPR1 as well as those related to cellular response to oxidative stress was more pronounced in *hpm91* than wild type. Strikingly, the amount of TRXh, PDI2, APX1 and CCPR1 increased at least 10-fold in *hpm91* during prolonged H_2_ production (Fig. 4B, downleft panel). Together with the previous finding of higher activity of ROS-scavenging enzymes in *hpm91* (Chen et al., 2016), we conclude that the lower amount of ROS observed in *hpm91* (Fig. 2C and D) could be attributed to the marked increase of both protein abundance of those and activity of the ROS-scavenging enzymes as well as putatively reduced PSI antenna mentioned above (Lu et al., 2021). A question rises how this is elicited in the PGR5-lacking mutants. We have found earlier that ROS tolerance was increased in *PGR5*-deficient mutants (*hpm91* and *pgr5*) and proposed a putative link between PGR5 and ROS metabolism in Chlamydomonas (Chen et al., 2016). In line with this, recent studies in *Arabidopsis* indicate that PGR5 was involved in shaping ROS metabolism during changing in light conditions (Haber et al., 2021). Thus, these experiments suggested, apart from the role of PGR5 as a core modulator of CEF (Schwenkert et al., 2022), an additional role of PGR5 in regulating ROS metabolism. In the present investigation, we also found a unique protein in *hpm91*, i.e. the EF-hand domain-containing thioredoxin, with an average fold-increase value of 6.13 during sustained H_2_ photoproduction (*SI Appendix*, Table S3). Because EF-hand-containing proteins are suggested to be the key transducers mediating calcium (Ca^2+^) signal transduction as a secondary messenger (Day et al., 2002; Kawasaki and Kretsinger, 2017), the remarkable accumulation of the EF-hand domain-containing thioredoxin (encoded by CHLREDRAFT_205510) observed in *hpm91* may allow us to propose another putative link between PGR5 and redox regulation involving calcium-signaling.

Moreover, N- and S-metabolisms of *hpm91* were enhanced during sustained H_2_ photoproduction. Apart from compromised carbon metabolism in *hpm91* mentioned above, we observed marked proteomic alterations in its N- and S-metabolic pathways during H_2_ production process. Considering that glutamine and glutamate are the major intracellular amino group donors for the synthesis of several other amino acids and nitrogen-containing compounds including purine and pyrimidine nucleobases (Zhang et al., 2018), we combined the functional groups of ‘amino acid pathways’, ‘terpenoid- and *de novo* biosynthesis of pyrimidine nucleobase’ (Fig. 4A) and referred as nitrogen metabolism in Table S3. In contrast to wild type, 3 proteins involved in nitrogen metabolism were only accumulated in *hpm91* under sulfur deprivation. These were aspartate aminotransferase AST3, glutamate dehydrogenase GDH and GDH2, showing upto 2.20-, 13.42- and 24.56-fold increase relative to their intial levels (*SI Appendix*, Table S3). The former (AST3) is known to be one of the major enzymes catalyzing conversion of glutamate and oxaloacetic acid (OAA) into asparate and 2-oxoglutarate (2-OG, also called alpha-KG), an intermediate of the TCA cycle (Ohashi et al., 2011) that serves as the metabolic basis for coupling N-and C-metabolisms in the organisms. Because AST3 was already higher-expressed in *hpm91* under normal condition (Table 1), the continued increase may strongly indicate its dominant role in asparate biosynthesis and/or maintenance of C/N balance in the *PGR5*-deficient *hpm91* mutant during prolonged H_2_ production.

More strikingly, we found marked accumulation of the latter two glutamate dehydrogenases (GDH and GDH2) in *hpm91* during sustained H_2_ production. These proteins are supposed to play an anaplerotic role in ammonium assimilation via conversion of Glu into 2-oxoglutarate (2-OG) and ammonium in Chlamydomonas (Moyano et al., 1995). Their remarkable increase exclusively observed in *hpm91* may implicate activation of this minor pathway of ammonium assimilation in the mutant under such conditions. Together with upregulation of NADH-dependent glutamate synthase GSN1 (*SI Appendix*, Table S3), the key enzyme of the major route GS-GOGAT cycle, we propose that, due to loss of PGR5, both the major and minor route of ammonium assimilation was operational in *hpm91* toward sulfur-deprived H_2_ photoproduction. It is generally known that enhanced ammonium assimilation requires higher demand of the carbon skeleton 2-oxoglutarate (Ohashi et al., 2011) for coupling between nitrogen and carbon metabolism (Zhang et al., 2018). By activating their anaplerotic role of the glutamate dehydrogenases toward producing more 2-oxoglutarate, carbon and nitrogen metabolism could be better coupled in the mutant cells than wild type under such conditions.

Regarding to S-metabolism, our results revealed remarkable accumulations for 5 key proteins involved in sulfur assimilation in *hpm91* during prolonged H_2_ production (*SI Appendix*, Table S3). These include not only adenylylphosphosulfate reductase APR1 showing continued increase in abundance (Table 1 and *SI Appendix*, Table S3) but also ATP-sulfurylases (ATS1, ATS2) as well as cysteine synthase OASTL3 towards incorporation SO_4_^2+^ into cysteine (Gonzales-Ballester and Grossman, 2009). Because the latter two proteins were found lower-expressed in *hpm91* under normal condition (Table 1), their marked accumulation as well as ATS1 during prolonged H_2_ production could be strongly indicating activation of the pathway towards cysteine biosynthesis (Fig. 4B, downright pannel). On the other hand, we observed upto 4.5-fold accumulation of APK1 in *hpm91* (*SI Appendix*, Table S3), implicating the sulfur-assimilation pathway toward sulfation of metabolites (sulfur-containing lipids, proteins and polysaccharides (Gonzales-Ballester and Grossman, 2009) was also activated in *hpm91* under such condition (Fig. 4B, downright panel). Consequently, overall sulfur assmilation was more enhanced in *hpm91* relative to wild type in the period of H_2_ photoproduction.

### Creating mutants with H_2_ production excess to *hpm91*

As the picture of *hpm91* emerges as better suited for sulfur-deprived anaerobic condition photosynthetically, metabolically and redox poisingly, we then tested a possibility of creating new strains with H_2_ production excess to *hpm91*. Fig. 5A shows a schematic diagram of the protocol tested in this work. An insertion mutant library derived from *hpm91* was constructed according to (Kindle, 1990; Zhao et al., 2017) followed by tranformants selection via zeocin resistance. Since increased stability of PSII is essential to sustain sufur-deprived H_2_ photoproduction in Chlamydomonas (Chen et al., 2016; 2019; Volgusheva et al., 2013), we applied a two-step mutant screening, *i. e*. Y(II) measurements using Maxi-Imaging PAM chlorophyll fluorometer (Walz, Germany) as the first followed by H_2_-generating phenotype confirmation by GC analysis as described (Chen et al., 2016; 2019; Sun et al., 2013). Preliminary mutant screening identified more than two hundred transformants with 10% increase of Y(II) values than *hpm91*. Subsequent screening by GC analysis identified one of the mutants, named *hpm91-108*, with significantly higher H_2_ production than *hpm91* (Fig. 5B). At 120 h, H_2_ production in *hpm91-108* was 55. 9% higher than *hpm91*. This result is the direct experimental evidence of our previous propose (Chen et al., 2016; 2019) that *hpm91* is a valuable strain for re-engineering the organism towards algal-H_2_ photoproduction.

**Fig. 5.**
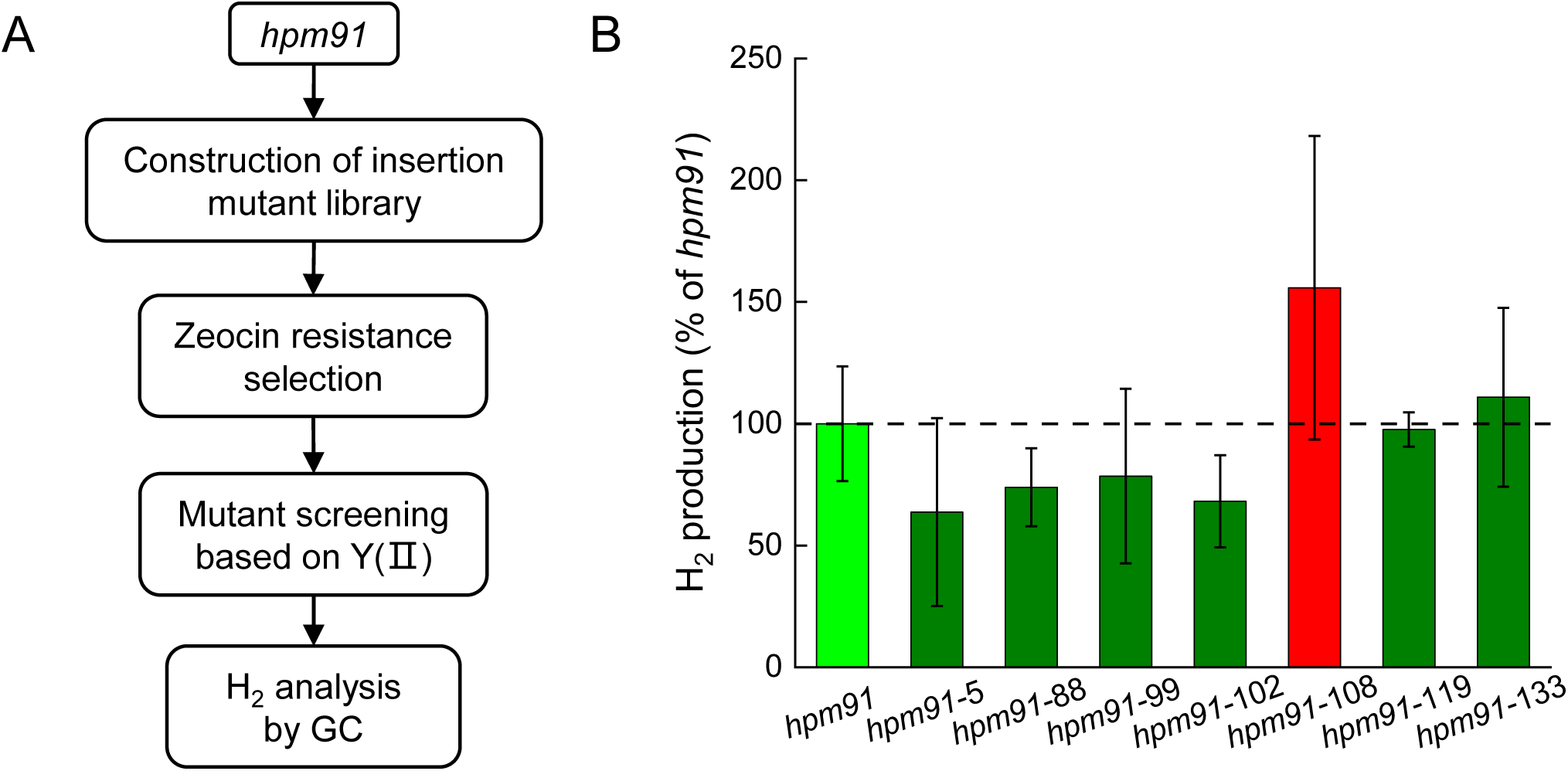
Screening for H_2_-production mutants excess to *hpm91. (A)* Outline of the screening method. *(B)* Phenotype confirmation of the *hpm91*-derived mutants. H_2_-producing capability of the selected mutants was determined with a GC-2014 gas chromatographer (Shimadzu; Japan) at 5 days of sulfur deprivation. Standard deviations were estimated from 3 biological replicates.

### Concluding remarks

In conclusion, analysis of the *hpm91* mutant has revealed a number of valuable properties towards development of sunlight-powered algal H_2_ production systems in the near future. First, it is largely up-scalable using the ‘two-step’ protocol of H_2_ induction by sulfur deprivation (Melis et al., 2000). In the both steps, up to 100-fold extention of PBR (10 L, mixotrophic growth) (Chen et al., 2019) and HPBR (10 L, sulfur-deprived H_2_ photoproduction) was achieved in the laboratory set-ups, leading to an average H_2_ output of 7287 ml/10L-HPBR around 26 days. Second, *hpm91* is robust during prolonged H_2_ production. In the absence of PGR5, *hpm91* shows competent viability than wild type and remains active over a long period of sulfur deprivation, most likely due to a decrease of intracellular ROS. Third, proteome of *hpm91* was modified metabolically (reinforced anti-ROS systems, compromised carbon metabolism, activation of anaplerotic route of ammonium assimilation and enhanced sulfur assmilation) and photosynthetically (optimal structure and function of PSII and PSI, reduced size of PSI antenna and CEF) towards sustained sulfur-deprived H_2_ photoproduction. Together with the successful isolation of new strain(s) with enhanced H_2_ capability relative to *hpm91*, we highlight that this mutant strain is not only scalable but also valuable in both basic and applied research aiming at elucidating molecular regulation mechanisms of H_2_ metabolism in the organism and developing advanced algal-H_2_ production biotechnology.

## Materials and methods

### Algal cultivation and H_2_ photoproduction

Wild-type *Chlamydomonas* strain CC400 was purchased from the Chlamydomonas Center (www.Chlamy.org) and the mutant strain *hpm91* was isolated in our laboratory and reported previously (Chen et al., 2016; 2019). Genetic analysis and phenotypic rescue of several fully complemented stains as well as immunoblot detection suggest that the loss of *PGR5* gene is responsible for the H_2_-overproduing phenotype of *hpm91* (Chen et al., 2016).

H_2_ production was induced using sulfur-deprivation method developed by Melis et al. (Melis et al., 2000) with minor modifications (Chen et al., 2010; Sun et al., 2013). Algal cells were grown in TAP medium (Gorman and Levine, 1965) at 25 °C under continuous light (60 µE m^-2^ s^-1^) until mid-exponential phase. Cells were pelleted and washed once with sulfur-depleted TAP medium followed by resuspending in the medium with desired cell density of 20- (for 10L-HPBR) and 25 μg ml^-1^ chlorophyll (*a* and *b*) (Arnon, 1949), respectively. H_2_ evolution was achieved via transferring the culture into small H_2_ photobioreactor (100 ml) for biochemical analysis and large gas-tight glass bottles (large-HPBR, upto 10 L) connected by a Teflon tube to storage glass cylinder for scaling-up studies followed by magnetic-bar-stiring cultivation at the same condition mentioned above (for small-HPBR) or upto 230 µE m^-2^ s^-1^ (for large HPBR), respectively. H_2_ gas accumulated in the headspace of the small HPBR was measured by gas chromatograph (GC-2014, Shimadzu, Japan) as previously (Chen et al., 2016; 2019; Sun et al., 2013; Zhao et al., 2013). The evolved H_2_ gas from the headspace of large HPBR was collected in inverted graduated cylinders and measured by the displacement of water (Chen et al., 2010).

### Intracellular ROS and cell growth analysis

Intracellular ROS content was determined using a MoFlo XDP high-speed flow cytometer (Beckman-Coulter, Inc. USA) by following the manufacturer’s instructions. Samples were prepared in an anoxia workstation (Longyao, LAI-3T; Shanghai, China) as (Chen et al., 2019). Cells were incubated with 10 mM H_2_DCFDA at 25 °C for 30 min in dark then examined by the flow cytometer. For detection of DCF green fluorescence, wavelength of excitation at 488 nm and emission at 510 to 550 nm was used. Rosup provided by a ROS assay kit (Beyotime Institute of Biotechnology, Haimen, China) was used as a positive control. Average fluorescence intensity of DCF from 3×10^5^ cells was recorded and data acquisition and analysis were carried out using Summit 5.2 software (Beckman-Coulter, Inc. USA). Morphology of algal cells was examined and photographed with a differential interference contrast microscopy (DIC) (Leica DM4500, Germany). Culture density were determined by cell counting using a hemocytometer.

### iTRAQ proteomics and data analysis

Protein extraction and sample preparation for iTRAQ labeling was done as described (Chen et al., 2010) with minor modifications (Ge et al., 2017). Frozen cells suspended in ice-cold extraction buffer were disrupted with glass beads (diameter 150-212 μm, Sigma) via vortexing 30 s/5 cycles/each with 1-min break on ice. After removed unbroken cells and insoluble debris, total proteins in the supernatant were precipitated with ice-cold 10% trichloroacetic acid (TCA) in acetone at -20°C. Protein pellets were washed with acetone by centrifugation and vacuum-dried. The proteins were resolubilized with 4% sodium dodecyl sulfate (SDS) in 0.1 M Tris-HCl, pH 7.6. Protein concentration was determined using BCA protein assay kit (Thermo Scientific, Rockford, IL).

Trypsin digestion of proteins was perfomed using the filter-aided sample preparation (FASP) method with slight modifications (Wisniewski et al., 2009). The resulting tryptic peptides were labeled with 8-plex iTRAQ reagents (AB Sciex Inc., MA, USA) alternatively by following manufacturer’s manual. The iTRAQ labeled samples were mixed together with equal ratios in amount, and the mixture was concentrated with a SpeedVac in subjection for fractionation by HPLC (Waters, e2695 separations) coupled with a phenomenex gemini-NX 5u C18 column (250 × 3.0 mm, 110 Å) (Torrance, CA, USA). The samples were then separated using a 97 min basic RP-LC gradient as described (Udeshi et al., 2013) and a flow rate of 0.4 mL/min was used. The separated samples were collected into 10 fractions and vacuum-dried prior to LC-MS/MS analysis by a TripleTOF 5600 mass spectrometer (AB SCIEX) coupled online to an Eksigent nanoLC Ultra in Information Dependent Mode. Peptides were separated on a C18 column (Acclaim PepMap C18, 250 mm x 75 μm x 5 μm, 100 Å, Dionex) with a 90 min nonlinear gradient of buffer B (100% ACN, 0.1% FA) from 3% to 30%. The gradient was set as 3%–8% B for 10 min, 8%–20% B for 60 min, 20%–30% B for 8 min, 30%–100% B for 2 min, and 100% B for 10 min at a flow rate of 300 nl/min. MS spectra survey scan was done across mass range of 350 to 1500 m/z and the spectra data were acquired at resolution 30000 with 250 ms accumulation per spectrum. 25 most intense ions from each MS scan were chosen for fragmentation from each MS spectrum with 2 sec minimum accumulation time for each precursor and dynamic exclusion for 18 sec. Tandem mass spectra were recorded in high sensitivity mode (resolution >15000) with rolling collision energy on and iTRAQ reagent collision energy adjustment on.

Peptide and protein identification and quantification was performed with the ProteinPilot 4.5 software (AB SCIEX) using the Paragon database search algorithm. The *Chlamydomonas* proteome sequences downloaded from UniProt (dated 2-07-2016) were used for the database searching with manual editing using the information (updated on 11-12-2019) in Uniprot database. The false discovery rate (FDR) analysis was performed using the software PSPEP integrated with the ProteinPilot. Confidence of quantitation for differentially expressed proteins was analyzed with the ProteinPilot Descriptive Statistics Template (PDST) (beta v3.07p, AB SCIEX). Gene ontology (GO) enrichment analysis of proteins was performed using DAVID Bioinformatic Resources 6.8 (https://david.ncifcrf.gov/summary.jsp) with the following parameters: ‘Gene List Enrichment’, ‘Chlamydomonas reinhardtii species’, ‘Uniprot Accession’. Functional annotation and classification of the differentially expressed proteins was mainly based on the search results of ‘Biological processes’ in DAVID analysis followed by manual editing using the updated Chlamydomonas genome data in NCBI (updated on 30-01-2018). Differentially expressed proteins were selected with *p* value <0.05 and cut-off >1.2 and <0.83 as up-and down-regulated, respectively.

### Generation and screening of *hpm91*-derived mutants

Insertion mutant library was constructed by transformation of *hpm91* using the glass bead method with KpnI-linearized plasmid pDble containing the *ble* gene conferring zeocin resistance (Kindle, 1990). Transformants growing on TAP plates with 10 μg mL^−1^ zeocin (Solarbio) were isolated. After cultured on TAP-S plates for one week, mutants were screened based on increased Y(Ⅱ) values relative to *hpm91* using Maxi-Imaging PAM chlorophyll fluorometer (Walz, Germany) as previously (Zhao et al., 2017) followed by H_2_ measurements by GC analysis (Chen et al., 2019; Sun et al., 2013).

## Data availability

All data needed to evaluate conclusions of this paper are present in the paper and in *SI Appendix*.

## Acknowledgements

This work was supported by funding from the National Natural Science Foundation of China (31470340), the Ministry of Science and Technology of China (2015CB150100) and CAS (XDB17030300, KGCX2-YW-373).

## Author contributions

M.C., Y.W. and F.H. conceived the project and designed the research; P.L., D.Y., M.C., J.Z., L.S. and X.H. carried out the experiments and data analysis; X. X., K. X., Y.Y. and Y. G. performed genomic sequencing and data analysis; P.L. and D.Y. prepared data files; P.L., D.Y., Y.W. and F.H. wrote the manuscript, with input from all authors. All authors approved the final manuscript.

## Competing interests

The authors declare no competing interests. This work is included in a patent application by Institute of Botany, Chinese Academy of Sciences. The patent 202110779011.4 covers a method of application.

## Notes

### Competing Interest Statement

The authors have declared no competing interest.

## References

Arnon, D.I. (1949). Copper enzymes in isolated chloroplasts. polyphenoloxidase in Beta vulgaris. Plant Physiol. 24, 1–15. https://doi.org/10.1104/pp.24.1.1.

Bayro-Kaiser, V., and Nelson, N. (2017). Microalgal hydrogen production: prospects of an essential technology for a clean and sustainable energy economy. Photosynth. Res. 133, 49–62. https://doi.org/10.1007/s11120-017-0350-6.

Chen F. Y.L., Yang, F., Wang, L., Guo, X.Q., Gao, F., Hua, C., Tan, C., Fang, L., Shan, R.Q., Zeng, W.J., Wang, B., Wang, R., Xu, X., Wei, X.F. (2020). CNGBdb: China National GeneBank DataBase. Hereditas(Beijing) 42, 799–809. https://doi.org/10.16288/j.yczz.20-080.

Chen, M., Liu, P., Zhang, F., Peng, L.W., and Huang, F. (2019). Photochemical characteristics of Chlamydomonas mutant hpm91 lacking proton gradient regulation 5 (PGR5) during sustained H2 photoproduction under sulfur deprivation. Int. J. Hydrogen Energ. 44, 31790–31799. https://doi.org/10.1016/j.ijhydene.2019.10.074.

Chen, M., Zhang, J., Zhao, L., Xing, J., Peng, L., Kuang, T., Rochaix, J.D., and Huang, F. (2016). Loss of algal proton gradient regulation 5 increases reactive oxygen species scavenging and H2 evolution. J. Integr. Plant Biol. 58, 943–946. https://doi.org/10.1111/jipb.12502.

Chen, M., Zhao, K., Sun, Y.L., Cui, S.X., Zhang, L.F., Yang, B., Wang, J., Kuang, T.Y., and Huang, F. (2010). Proteomic analysis of hydrogen photoproduction in sulfur-deprived Chlamydomonas cells. J. Proteome Res. 9, 3854–3866. https://doi.org/10.1021/pr100076c.

Chen, Y., Chen, Y., Shi, C., Huang, Z., Zhang, Y., Li, S., Li, Y., Ye, J., Yu, C., Li, Z., et al. (2018). SOAPnuke: a MapReduce acceleration-supported software for integrated quality control and preprocessing of high-throughput sequencing data. Gigascience 7, 1–6. https://doi.org/10.1093/gigascience/gix120.

Day, I.S., Reddy, V.S., Shad Ali, G., and Reddy, A.S. (2002). Analysis of EF-hand-containing proteins in Arabidopsis. Genome Biol. 3, RESEARCH0056.1. https://doi.org/10.1186/gb-2002-3-10-research0056.

Forestier, M., King, P., Zhang, L.P., Posewitz, M., Schwarzer, S., Happe, T., Ghirardi, M.L., and Seibert, M. (2003). Expression of two [Fe]-hydrogenases in Chlamydomonas reinhardtii under anaerobic conditions. Eur. J. Biochem. 270, 2750–2758. https://doi.org/10.1046/j.1432-1033.2003.03656.

Ge, H.T., Fang, L.F., Huang, X.H., Wang, J.L., Chen, W.Y., Liu, Y., Zhang, Y.Y., Wang, X.R., Xu, W., He, Q.F., and Wang, Y.C. (2017). Translating divergent environmental stresses into a common proteome response through the histidine kinase 33 (Hik33) in a model Cyanobacterium. Mol. Cell Proteomics 16, 1258–1274. https://doi.org/10.1074/mcp.M116.068080

Gfeller, R.P., and Gibbs, M. (1984). Fermentative metabolism of Chlamydomonas reinhardtii 1: I. analysis of fermentative products from starch in dark and light. Plant Physiol. 75, 212–218. https://doi.org/10.1104/pp.75.1.212.

Ghirardi, M.L. (2015). Implementation of photobiological H2 production: the O2 sensitivity of hydrogenases. Photosynth. Res. 125, 383–393. https://doi.org/10.1007/s11120-015-0158-1.

Gonzales-Ballester, D., and Grossman, A.R. (2009). Sulfur: From acquisition to assimilation. In The Chlamydomonas Sourcebook Second Edition, D.B. Stern, ed. (Elsevier), pp. 159–187.

Gorman, D.S., and Levine, R.P. (1965). Cytochrome f and plastocyanin - their sequence in photosynthetic electron transport chain of Chlamydomonas reinhardi. Proc. Natl. Acad. Sci. U.S.A. 54, 1665–1669. https://doi.org/10.1073/pnas.54.6.1665.

Greenbaum, E. (1988). Energetic efficiency of hydrogen photoevolution by algal water splitting. Biophys. J. 54, 365–368. https://doi.org/10.1016/S0006-3495(88)82968-0.

Guo, X.Q., Chen, F.Z., Gao, F., Li, L., Liu, K., You, L.J., Hua, C., Yang, F., Liu, W.L., Peng, C.H., et al. (2020). CNSA: a data repository for archiving omics data. Database 2020, baaa055 https://doi.org/10.1093/database/baaa055.

GutierrezMarcos, J.F., Roberts, M.A., Campbell, E.I., and Wray, J.L. (1996). Three members of a novel small gene-family from Arabidopsis thaliana able to complement functionally an Escherichia coli mutant defective in PAPS reductase activity encode proteins with a thioredoxin-like domain and ‘‘APS reductase’’ activity. Proc. Natl. Acad. Sci. U.S.A. 93, 13377–13382. https://doi.org/10.1073/pnas.93.23.13377.

Haber, Z., Lampl, N., Meyer, A.J., Zelinger, E., Hipsch, M., and Rosenwasser, S. (2021). Resolving diurnal dynamics of the chloroplastic glutathione redox state in Arabidopsis reveals its photosynthetically derived oxidation. Plant Cell 33, 1828–1844. https://doi.org/10.1093/plcell/koab068.

Hemschemeier, A., Melis, A., and Happe, T. (2009). Analytical approaches to photobiological hydrogen production in unicellular green algae. Photosynth. Res. 102, 523–540. https://doi.org/10.1007/s11120-009-9415-5.

Iwai, M., Takizawa, K., Tokutsu, R., Okamuro, A., Takahashi, Y., and Minagawa, J. (2010). Isolation of the elusive supercomplex that drives cyclic electron flow in photosynthesis. Nature 464, 1210–1213. https://doi.org/10.1038/nature08885.

Johnson, X., Steinbeck, J., Dent, R.M., Takahashi, H., Richaud, P., Ozawa, S.I., Houille-Vernes, L., Petroutsos, D., Rappaport, F., Grossman, A.R., et al. (2014). Proton gradient regulation 5-mediated cyclic electron flow under ATP-or redox-limited conditions: A study of ?ATPase pgr5 and ?rbcL pgr5 mutants in the green alga Chlamydomonas reinhardtii. Plant Physiol. 165, 438–452. https://doi.org/10.1104/pp.113.233593.

Kawasaki, H., and Kretsinger, R.H. (2017). Structural and functional diversity of EF-hand proteins: Evolutionary perspectives. Protein Sci. 26, 1898–1920. https://doi.org/10.1002/pro.3233.

Kindle, K.L. (1990). High-Frequency nuclear transformation of Chlamydomonas reinhardtii. Proc. Natl. Acad. Sci. U.S.A. 87, 1228–1232. https://doi.org/10.1073/pnas.87.3.1228.

Kosourov, S., Tsygankov, A., Seibert, M., and Ghirardi, M.L. (2002). Sustained hydrogen photoproduction by Chlamydomonas reinhardtii: Effects of culture parameters. Biotechnol. Bioeng 78, 731–740. https://doi.org/10.1002/bit.10254.

Kosourov, S.N., Ghirardi, M.L., and Seibert, M. (2011). A truncated antenna mutant of Chlamydomonas reinhardtii can produce more hydrogen than the parental strain. Int. J. Hydrogen Energ. 36, 2044–2048. https://doi.org/10.1016/j.ijhydene.2010.10.041.

Kruse, O., Rupprecht, J., Bader, K.P., Thomas-Hall, S., Schenk, P.M., Finazzi, G., and Hankamer, B. (2005). Improved photobiological H2 production in engineered green algal cells. J. Biol. Chem. 280, 34170–34177. https://doi.org/10.1074/jbc.M503840200.

Li, H. (2013). Aligning sequence reads, clone sequences and assembly contigs with BWA-MEM. 1303.3997. https://doi.org/10.6084/M9.FIGSHARE.963153.V1.

Lu, Y., Gan, Q., Iwai, M., Alboresi, A., Burlacot, A., Dautermann, O., Takahashi, H., Crisanto, T., Peltier, G., Morosinotto, T., et al. (2021). Role of an ancient light-harvesting protein of PSI in light absorption and photoprotection. Nat. Commun. 12, 679. https://doi.org/10.1038/s41467-021-20967-1.

Merchant, S.S., Prochnik, S.E., Vallon, O., Harris, E.H., Karpowicz, S.J., Witman, G.B., Terry, A., Salamov, A., Fritz-Laylin, L.K., Marechal-Drouard, L., et al. (2007). The Chlamydomonas genome reveals the evolution of key animal and plant functions. Science 318, 245–251. https://doi.org/10.1126/science.1143609.

Melis, A., Zhang, L.P., Forestier, M., Ghirardi, M.L., and Seibert, M. (2000). Sustained photobiological hydrogen gas production upon reversible inactivation of oxygen evolution in the green alga Chlamydomonas reinhardtii. Plant Physiol. 122, 127–135. https://doi.org/10.1104/pp.122.1.127.

Moyano, E., Cardenas, J., and Munozblanco, J. (1995). Involvement of NAD(P)^+^-glutamate dehydrogenase isoenzymes in carbon and nitrogen metabolism in Chlamydomonas reinhardtii. Physiol. Plantarum 94, 553–559. https://doi.org/10.1034/j.1399-3054.1995.940403.x.

Munekage, Y., Hojo, M., Meurer, J., Endo, T., Tasaka, M., and Shikanai, T. (2002). PGR5 is involved in cyclic electron flow around photosystem I and is essential for photoprotection in Arabidopsis. Cell 110, 361–371. https://doi.org/10.1016/S0092-8674(02)00867-X.

Nishiyama, H., Yamada, T., Nakabayashi, M., Maehara, Y., Yamaguchi, M., Kuromiya, Y., Nagatsuma, Y., Tokudome, H., Akiyama, S., Watanabe, T., et al. (2021). Photocatalytic solar hydrogen production from water on a 100-m^2^ scale. Nature 598, 304–307. https://doi.org/10.1038/s41586-021-03907-3.

Ohashi, Y., Shi, W., Takatani, N., Aichi, M., Maeda, S., Watanabe, S., Yoshikawa, H., and Omata, T. (2011). Regulation of nitrate assimilation in cyanobacteria. J. Exp. Bot. 62, 1411–1424. https://doi.org/10.1093/jxb/erq427.

Philipps, G., Krawietz, D., Hemschemeier, A., and Happe, T. (2011). A pyruvate formate lyase-deficient Chlamydomonas reinhardtii strain provides evidence for a link between fermentation and hydrogen production in green algae. Plant J. 66, 330–340. https://doi.org/10.1111/j.1365-313X.2011.04494.x.

Ravina, C.G., Chang, C.I., Tsakraklides, G.P., McDermott, J.P., Vega, J.M., Leustek, T., Gotor, C., and Davies, J.P. (2002). The sac mutants of Chlamydomonas reinhardtii reveal transcriptional and posttranscriptional control of cysteine biosynthesis. Plant Physiol. 130, 2076–2084. https://doi.org/10.1104/pp.012484.

Schonfeld, C., Wobbe, L., Borgstadt, R., Kienast, A., Nixon, P.J., and Kruse, O. (2004). The nucleus-encoded protein MOC1 is essential for mitochondrial light acclimation in Chlamydomonas reinhardtii. J. Biol. Chem. 279, 50366–50374. https://doi.org/10.1074/jbc.M408477200.

Schwenkert, S., Fernie, A.R., Geigenberger, P., Leister, D., Mohlmann, T., Naranjo, B., and Neuhaus, H.E. (2022). Chloroplasts are key players to cope with light and temperature stress. Trends Plant Sci. https://doi.org/10.1016/j.tplants.2021.12.004.

Scoma, A., Krawietz, D., Faraloni, C., Giannelli, L., Happe, T., and Torzillo, G. (2012). Sustained H2 production in a Chlamydomonas reinhardtii D1 protein mutant. J. Biotechnol. 157, 613–619. https://doi.org/10.1016/j.jbiotec.2011.06.019.

Setya, A., Murillo, M., and Leustek, T. (1996). Sulfate reduction in higher plants: Molecular evidence for a novel 5’-adenylylsulfate reductase. Proc. Natl. Acad. Sci. U.S.A. 93, 13383–13388. https://doi.org/10.1073/pnas.93.23.13383.

Shen, J.R. (2015). The structure of Photosystem II and the mechanism of water oxidation in photosynthesis. Annu. Rev. Plant Biol. 66, 23–48. https://doi.org/10.1146/annurev-arplant-050312-120129.

Shen, L.L., Huang, Z.H., Chang, S.H., Wang, W.D., Wang, J.F., Kuang, T.Y., Han, G.Y., Shen, J.R., and Zhang, X. (2019). Structure of a C2S2M2N2 type PSII-LHCII supercomplex from the green alga Chlamydomonas reinhardtii. Proc. Natl. Acad. Sci. U.S.A. 116, 21246–21255. https://doi.org/10.1073/pnas.1912462116.

Steinbeck, J., Nikolova, D., Weingarten, R., Johnson, X., Richaud, P., Peltier, G., Hermann, M., Magneschi, L., and Hippler, M. (2015). Deletion of proton gradient regulation 5 (PGR5) and PGR5-like 1 (PGRL1) proteins promote sustainable light-driven hydrogen production in Chlamydomonas reinhardtii due to increased PSII activity under sulfur deprivation. Front. Plant Sci. 6. 892 https://doi.org/10.3389/fpls.2015.00892.

Su, X.D., Ma, J., Pan, X.W., Zhao, X.L., Chang, W.R., Liu, Z.F., Zhang, X.Z., and Li, M. (2019). Antenna arrangement and energy transfer pathways of a green algal photosystem-I-LHCI supercomplex. Nat. Plants 5, 273–281. https://doi.org/10.1038/s41477-019-0380-5.

Suga, M., Ozawa, S.I., Yoshida-Motomura, K., Akita, F., Miyazaki, N., and Takahashi, Y. (2019). Structure of the green algal photosystem I supercomplex with a decameric light-harvesting complex I. Nat. Plants 5, 626–636. https://doi.org/10.1038/s41477-019-0438-4.

Sun, Y.L., Chen, M., Yang, H.M., Zhang, J., Kuang, T.Y., and Huang, F. (2013). Enhanced H2 photoproduction by down-regulation of ferredoxin-NADP^+^ reductase (FNR) in the green alga Chlamydomonas reinhardtii. Int. J. Hydrogen Energ. 38, 16029–16037. https://doi.org/10.1016/j.ijhydene.2013.10.011.

Suorsa, M., Jarvi, S., Grieco, M., Nurmi, M., Pietrzykowska, M., Rantala, M., Kangasjarvi, S., Paakkarinen, V., Tikkanen, M., Jansson, S., and Aro, E.M. (2012). Proton gradient regulation5 is essential for proper acclimation of arabidopsis photosystem I to naturally and artificially fluctuating light conditions. Plant Cell 24, 2934–2948. https://doi.org/10.1105/tpc.112.097162.

Takahashi, H., Clowez, S., Wollman, F.A., Vallon, O., and Rappaport, F. (2013). Cyclic electron flow is redox-controlled but independent of state transition. Nat. Commun. 4. 1954 https://doi.org/10.1038/ncomms2954.

Thorvaldsdottir, H., Robinson, J.T., and Mesirov, J.P. (2013). Integrative Genomics Viewer (IGV): high-performance genomics data visualization and exploration. Brief. Bioinform. 14, 178–192. https://doi.org/10.1093/bib/bbs017.

Tolleter, D., Ghysels, B., Alric, J., Petroutsos, D., Tolstygina, I., Krawietz, D., Happe, T., Auroy, P., Adriano, J.M., Beyly, A., et al. (2011). Control of hydrogen photoproduction by the proton gradient generated by cyclic electron flow in Chlamydomonas reinhardtii. Plant Cell 23, 2619–2630. https://doi.org/10.1105/tpc.111.086876.

Trosch, R., Barahimipour, R., Gao, Y., Badillo-Corona, J.A., Gotsmann, V.L., Zimmer, D., Muhlhaus, T., Zoschke, R., and Willmund, F. (2018). Commonalities and differences of chloroplast translation in a green alga and land plants. Nat. Plants 4, 564–575. https://doi.org/10.1038/s41477-018-0211-0.

Udeshi, N.D., Svinkina, T., Mertins, P., Kuhn, E., Mani, D.R., Qiao, J.W., and Carr, S.A. (2013). Refined preparation and use of anti-diglycine remnant (K-epsilon-GG) antibody enables routine quantification of 10,000s of ubiquitination sites in single proteomics experiments. Mol. Cell Proteomics 12, 825–831. https://doi.org/10.1074/mcp.O112.027094.

Umena, Y., Kawakami, K., Shen, J.R., and Kamiya, N. (2011). Crystal structure of oxygen-evolving photosystem II at a resolution of 1.9 angstrom. Nature 473, 55–60. https://doi.org/10.1038/nature09913.

Volgusheva, A., Styring, S., and Mamedov, F. (2013). Increased photosystem II stability promotes H2 production in sulfur-deprived Chlamydomonas reinhardtii. Proc. Natl. Acad. Sci. U. S. A. 110, 7223–7228. https://doi.org/10.1073/pnas.1220645110.

Wang, H., Alvarez, S., and Hicks, L.M. (2012). Comprehensive comparison of iTRAQ and label-free LC-based quantitative proteomics approaches using two Chlamydomonas reinhardtii strains of interest for biofuels engineering. J. Proteome Res. 11, 487–501. https://doi.org/10.1021/pr2008225.

Weigelt, K., Kuster, H., Rutten, T., Fait, A., Fernie, A.R., Miersch, O., Wasternack, C., Emery, R.J.N., Desel, C., Hosein, F., et al. (2009). ADP-glucose pyrophosphorylase-deficient pea embryos reveal specific transcriptional and metabolic changes of carbon-nitrogen metabolism and stress responses. Plant Physiol. 149, 395–411. https://doi.org/10.1104/pp.108.129940.

Wisniewski, J.R., Zougman, A., Nagaraj, N., and Mann, M. (2009). Universal sample preparation method for proteome analysis. Nat. Methods 6, 359–362. https://doi.org/10.1038/nmeth.1322

Yacoby, I., Pochekailov, S., Toporik, H., Ghirardi, M.L., King, P.W., and Zhang, S.G. (2011). Photosynthetic electron partitioning between [FeFe]-hydrogenase and ferredoxin:NADP^+^-oxidoreductase (FNR) enzymes in vitro. Proc. Natl. Acad. Sci. U. S. A. 108, 9396–9401. https://doi.org/10.1073/pnas.1103659108.

Yang, W.Q., Catalanotti, C., Wittkopp, T.M., Posewitz, M.C., and Grossman, A.R. (2015). Algae after dark: mechanisms to cope with anoxic/hypoxic conditions. Plant J. 82, 481–503. https://doi.org/10.1111/tpj.12823.

Zhang, C.C., Zhou, C.Z., Burnap, R.L., and Peng, L. (2018). Carbon/Nitrogen metabolic balance: Lessons from Cyanobacteria. Trends Plant Sci. 23, 1116–1130. https://doi.org/10.1016/j.tplants.2018.09.008.

Zhang, L.P., Happe, T., and Melis, A. (2002). Biochemical and morphological characterization of sulfur-deprived and H2-producing Chlamydomonas reinhardtii (green alga). Planta 214, 552–561. https://doi.org/10.1007/s004250100660.

Zhang, Z.D., Shrager, J., Jain, M., Chang, C.W., Vallon, O., and Grossman, A.R. (2004). Insights into the survival of Chlamydomonas reinhardtii during sulfur starvation based on microarray analysis of gene expression. Eukaryot. Cell 3, 1331–1348. https://doi.org/10.1128/Ec.3.5.1331-1348.2004.

Zhao, L., Chen, M., Cheng, D.M., Yang, H.M., Sun, Y.L., Zhou, H.Y., and Huang, F. (2013). Different B-Type Methionine Sulfoxide Reductases in Chlamydomonas May Protect the Alga against High-Light, Sulfur-Depletion, or Oxidative Stress. J. Integr. Plant Biol. 55, 1054–1068. https://doi.org/10.1111/jipb.12104.

Zhao, L., Cheng, D.M., Huang, X.H., Chen, M., Dall’Osto, L., Xing, J.L., Gao, L.Y., Li, L.Y., Wang, Y., Bassi, R., et al. (2017). A light harvesting complex-like protein in maintenance of photosynthetic components in Chlamydomonas. Plant Physiol. 174, 2419–2433. https://doi.org/10.1104/pp.16.01465.

