## Supplemental data files for "Chlamydomonas mutant *hpm91* lacking PGR5 is a scalable and valuable strain for algal hydrogen (H_2_) production"

**This PDF file includes:**

Figures S1 to S3
Tables S1 to S4
Legend for Movie S1
Legends for Datasets S1 to S6

**Other supplementary materials for this manuscript include the following:**

Movie S1
Datasets S1 to S6

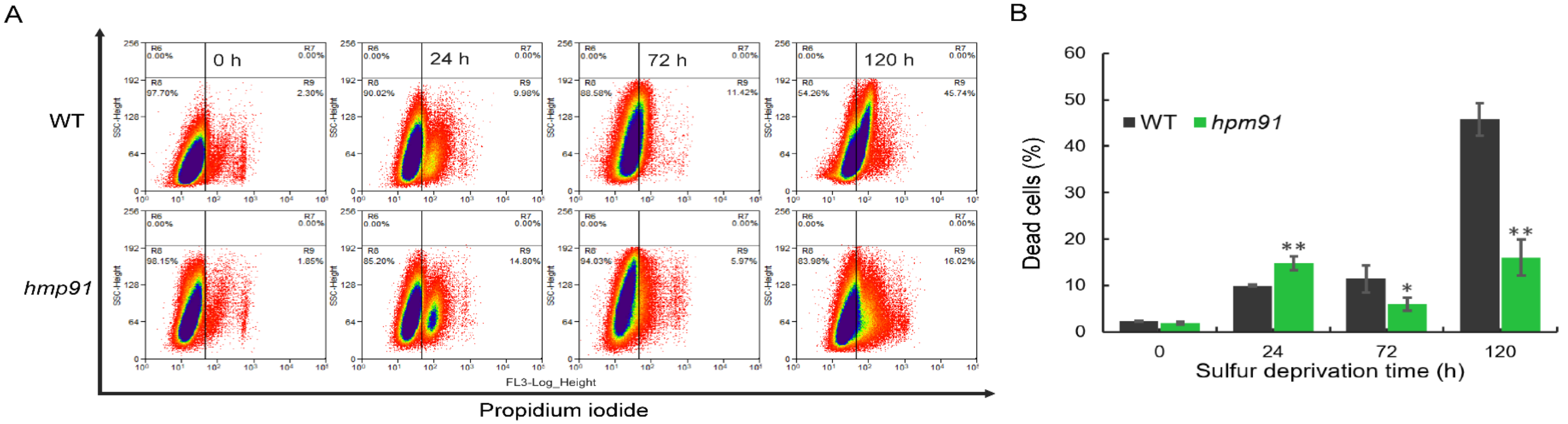

**Fig. S1.** Comparison of percentage of dead cells in the culture of *hpm91* and wild type during 120 h of sulfur-deprived $H_2$ production. Cells were incubated with propidium iodide (PI; 10 $\mu g \cdot mL^{-1}$) in an anoxia workstation (Longyao, LAI-3T; Shanghai, China). Fluorescence of PI inside the cells was detected with excitation at 488 nm and emission 510 to 550 nm. Data acquisition and analysis was carried out using Summit 5.2 software (Beckman Coulter, Inc. USA). The percentage of membrane damaged cells (dead cells) was a proportion of the total population. Experiments were repeated three times with similar results. * and ** refer to $p$-values $<0.05$ and $<0.01$ in Student's t-test, respectively.

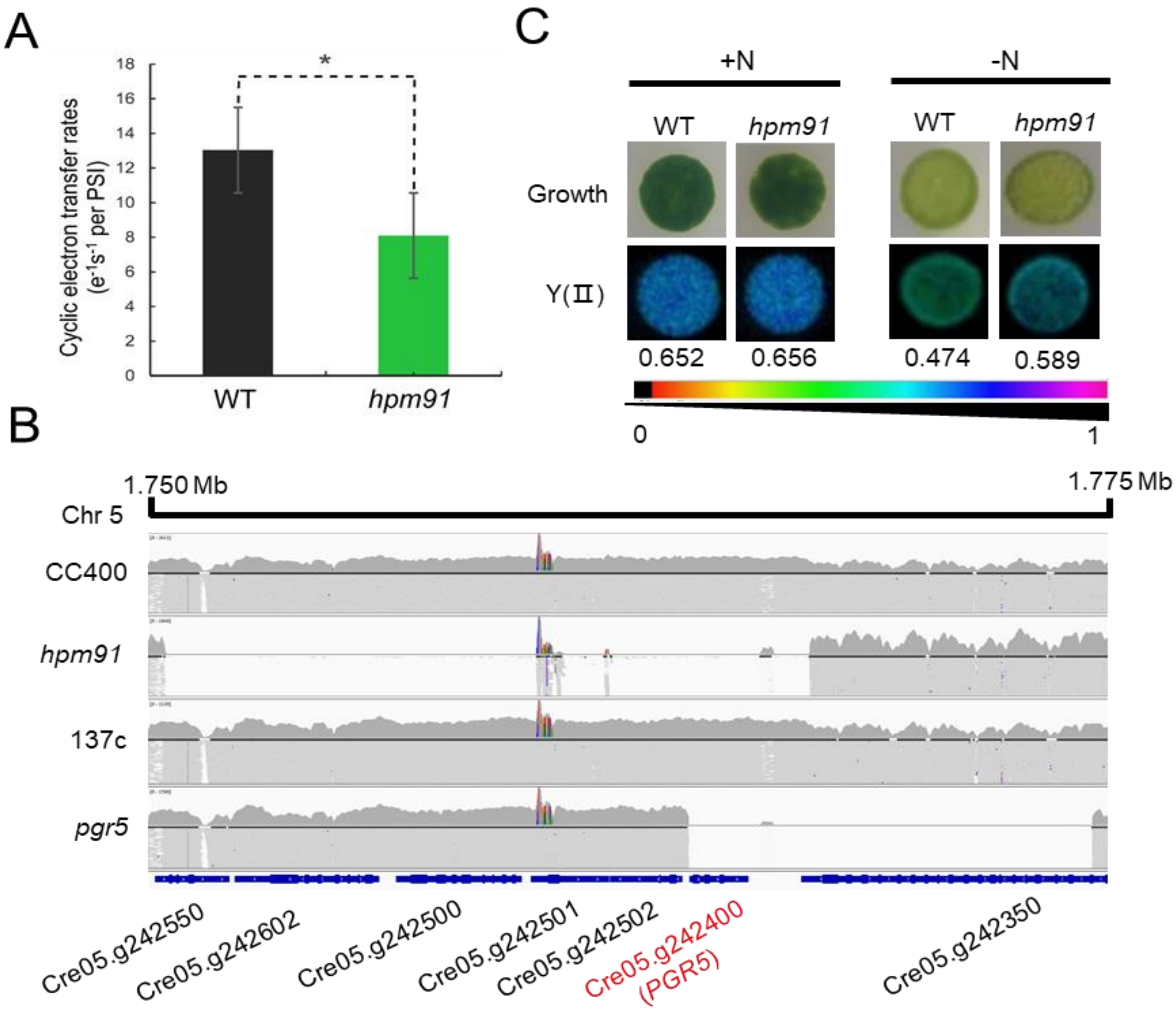

A
Cyclic electron transfer rates ($e^{-1}s^{-1}$ per PSI)
18
16
14
12
10
8
6
4
2
0
*
WT
hpm91
B
1.750 Mb
1.775 Mb
Chr 5
CC400
hpm91
137c
pgr5
Cre05.g242550
Cre05.g242602
Cre05.g242500
Cre05.g242501
Cre05.g242502
Cre05.g242400 (PGR5)
Cre05.g242350
C
+N
-N
WT
hpm91
WT
hpm91
Growth
Y(Ⅱ)
0.652
0.656
0.474
0.589
0
1

**Fig. S2.** Functional and genomic verification of *PGR5* deletion in *hpm91*. *(A)* Comparison of CEF in wild type and *hpm91*. Measurements were performed with a JTS-10 spectrometer (BioLogic, France) using samples prepared according to (Takahashi et al., 2013). Standard deviations were estimated from 3 biological replicates. Experiments were repeated three times with similar results. *refers to p-values $<$0.05 in Student's t-test. *(B)* Genomic coverage of four strains of *hpm91*, CC400, 137c and *pgr5* (these two strains were also purchased from Chlamydomonas Center, www.Chlamy.org). Genomic DNA was isolated using the Plant Genomic DNA Kit (TiangenBiotech; Beijing, China) as described in (Chen et al., 2016). High-throughput sequencing was carried out in DNBSEQ-T1 platform (BGI-Shenzhen, China) with clean data obtained via SOAPnuke software (Chen et al., 2018) followed by mapping to Chlamydomonas reference genome (Merchant et al., 2007) using BWA-mem algorithms (Li, 2013) and deposited at the CNGBdb database (Chen et al., 2020; Guo et al., 2020) (https://db.cngb.org/search; accession No. CNP0002674). Further analysis of genomic region containing *PGR5* was done with IGV software (Thorvaldsdottir et al., 2013). The thick blue lines in the bottom indicates genes on the chromosome 5: 1.750-1.775 Mb. Each grey line indicates the reads coverage of the four strains, which reveals two large deletions in the strains of *hpm91* and *pgr5* both with the gene loss of *PGR5*. *(C)* Loss of PGR5 led to enhanced tolerance to N-starvation. Mid-exponential phase cells with density of 2 x $10^6$ cells $mL^{-1}$ were spotted on TAP plates (+N, -N) and grown for 5 days. Y(II) was measured with as Maxi-Imaging PAM (Walz, Germany) as described in (Zhao et al., 2017). Experiments were repeated three times with similar results.

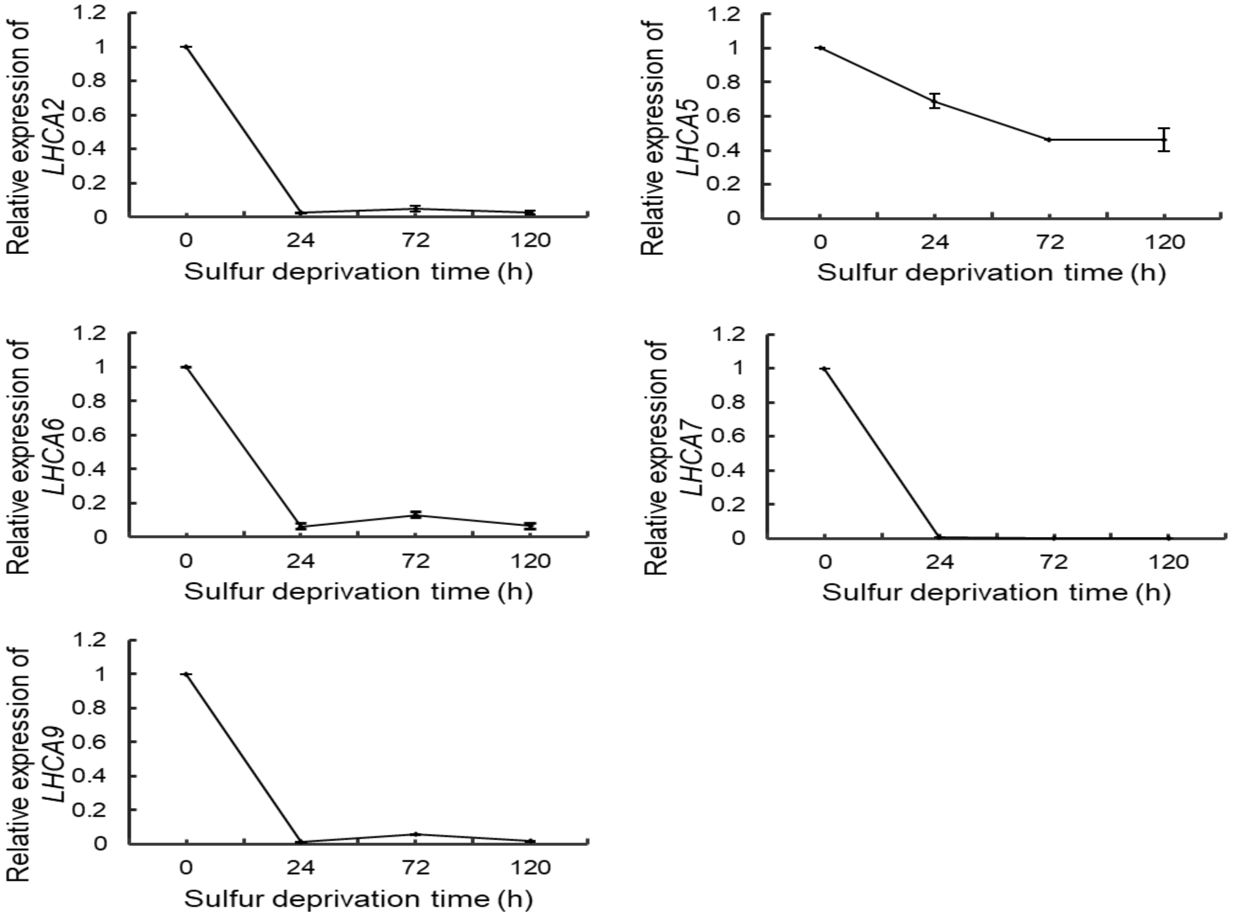

**Fig. S3.** qRT-PCR analysis of relative gene expression of the LHCAs in *hpm91* during 120 h of sulfur-deprived $H_2$ production. Measurements were done according to (Zhao et al., 2017). Standard deviations were estimated from 3 biological replicates each with 3 technical replicates (n = 9). Similar results were obtained in at least three independent experiments. The CBLP gene was used as a control.

**Table S1.** Comparison of $H_2$ output from *hpm91* in 10L-HPBR under different light intensities.

| No. | | Light irradiance (μE $m^{-2}$ $s^{-1}$) | Chlorophyll (μg/ml) | SD | Duration time (d) | SD | Collected $H_2$ (ml) | SD |
|---|---|---|---|---|---|---|---|---|
| Experiment | 1 | 130 | 20.0 | | 33 | | 5315 | |
| | 2 | 130 | 18.4 | | 25 | | 4465 | |
| | 3 | 130 | 19.1 | | 27 | | 4718 | |
| Average | | | 19.2 | 0.8 | 28 | 4 | 4832 | 436 |
| Experiment | 1 | 230 | 18.3 | | 33 | | 8395 | |
| | 2 | 230 | 18.3 | | 21 | | 6505 | |
| | 3 | 230 | 18.1 | | 23 | | 6960 | |
| Average | | | 18.2 | 0.1 | 26 | 6 | 7287 | 986 |

**Table S2.** List of differentially expressed proteins in wild type delineated from Dataset S5.

| Ranking | GOPB | Uniprot accession | Gene ID | Protein name | Ratio (WTt/WT0) | | | *p*-value | | |
|---|---|---|---|---|---|---|---|---|---|---|
| | | | | | 24 h | 72 h | 120 h | 24 h | 72 h | 120 h |
| 1 | Translation | A8J9T0 | RPS12 | 40S ribosomal protein S12 | 2.18 | 1.34 | 0.40 | 2.50E-01 | 2.97E-01 | 2.45E-02 |
| | | A8IXG3 | RPS21 | 40S ribosomal protein S21 | 1.10 | 0.63 | 0.72 | 7.51E-01 | 2.43E-02 | 4.78E-01 |
| | | A8I0I1 | RPS24 | 40S ribosomal protein S24 | 1.63 | 0.66 | 0.30 | 3.45E-01 | 4.00E-02 | 4.31E-02 |
| | | A8J576 | RPS27-A | 40S ribosomal protein S27 | 0.43 | 0.14 | 0.07 | 1.51E-03 | 1.08E-07 | 8.81E-06 |
| | | A8J0V6 | RPS27-B | 40S ribosomal protein S27 | 2.96 | 2.96 | 2.08 | 1.51E-02 | 7.29E-03 | 2.67E-02 |
| | | A8HS48 | CHLREDRAFT_168484 | 40S ribosomal protein S3a | 1.33 | 0.39 | 0.10 | 6.15E-01 | 9.77E-05 | 1.07E-05 |
| | | A8HVQ1 | RPS8 | 40S ribosomal protein S8 | 0.32 | 0.16 | 0.08 | 4.41E-04 | 1.60E-06 | 5.04E-06 |
| | | A8IB25 | CHLREDRAFT_126059 | 40S ribosomal protein SA | 1.13 | 0.58 | 0.22 | 5.50E-01 | 3.48E-05 | 2.53E-03 |
| | | A8IUV7 | RPL13 | 60S ribosomal protein L13 | 1.72 | 0.34 | 0.09 | 2.16E-01 | 1.76E-07 | 1.31E-06 |
| | | A8IKZ2 | RPL18 | 60S ribosomal protein L18 | 1.21 | 0.35 | 0.09 | 4.94E-01 | 5.94E-05 | 8.35E-06 |
| | | A8IQC1 | RPL27 | 60S ribosomal protein L27 | 14.85 | 6.63 | 3.01 | 4.76E-04 | 2.12E-02 | 5.73E-02 |
| | | Q8GUQ9 | RPL38 | 60S ribosomal protein L38 | 4.17 | 0.91 | 0.63 | 3.16E-02 | 7.11E-01 | 2.99E-01 |
| | | A8J1B6 | ANT1 | Adenine nucleotide translocator | 0.49 | 0.48 | 0.36 | 3.79E-02 | 1.79E-02 | 5.40E-04 |
| | | Q84X74 | MPC1b\|MPC1a | CR057 protein | 1.42 | 4.27 | 2.72 | 6.31E-02 | 2.67E-02 | 2.53E-02 |
| | | A8JHX9 | EFG2 | Elongation factor 2 | 1.77 | 1.14 | 0.20 | 1.19E-01 | 6.39E-01 | 1.70E-03 |
| | | A8JCA8 | EFG5 | Elongation factor EF-Tu-like protein | 1.15 | 2.23 | 2.48 | 2.36E-01 | 6.07E-03 | 7.58E-03 |
| | | A8IAC7 | EIF5Ba | Eukaryotic initiation factor | 0.51 | 0.83 | 0.64 | 2.69E-03 | 3.04E-01 | 5.59E-02 |
| | | A8HX38 | EEF1 | Eukaryotic translation elongation factor 1 alpha 1 | 1.13 | 0.67 | 0.10 | 3.74E-01 | 2.16E-01 | 2.37E-06 |
| | | A8J8J8 | CHLREDRAFT_120661 | GTPase Der | 0.44 | 0.43 | 0.44 | 1.68E-03 | 5.38E-04 | 5.25E-02 |
| | | A8JCK3 | GBA1 | GTP binding protein TypA | 0.48 | 0.63 | 1.11 | 3.77E-04 | 2.70E-01 | 8.43E-01 |
| | | A8IFH4 | LPA1 | Membrane GTP-binding protein LepA | 0.39 | 0.51 | 0.44 | 2.40E-02 | 4.70E-02 | 1.66E-02 |
| | | A8JB67 | NHP2 | Nucleolar protein small subunit of H/ACA snoRNPs | 0.70 | 0.37 | 0.15 | 1.23E-01 | 1.15E-05 | 3.91E-05 |
| | | A8J9X5 | CHLREDRAFT_18599 | Predicted protein | 0.86 | 0.54 | 0.74 | 5.62E-01 | 7.32E-02 | 1.98E-03 |
| | | A8I9M5 | CHLREDRAFT_141578 | Predicted protein | 0.14 | 0.14 | 0.16 | 7.34E-08 | 5.99E-07 | 5.02E-04 |
| | | A8JHJ5 | CHLRE_15g641200v5 | Predicted protein | 0.08 | 0.15 | 0.12 | 2.52E-07 | 7.48E-06 | 3.48E-06 |
| | | A8I2G3 | CHLREDRAFT_112251 | Predicted protein | 0.57 | 0.48 | 0.51 | 3.46E-03 | 2.43E-03 | 5.58E-02 |
| | | A8HXD0 | CHLREDRAFT_134139 | Predicted protein | 0.96 | 0.70 | 0.72 | 3.27E-01 | 7.35E-03 | 1.42E-01 |
| | | A8HZV5 | CHLREDRAFT_187641 | Predicted protein | 0.50 | 0.46 | 0.47 | 3.73E-02 | 5.22E-02 | 1.87E-03 |
| | | A8I4T2 | RPL10a | Ribosomal protein | 21.35 | 14.90 | 11.18 | 1.64E-02 | 1.74E-02 | 3.30E-02 |
| | | A8IZK3 | RPL10 | Ribosomal protein L10 | 1.19 | 0.30 | 0.12 | 5.62E-01 | 3.35E-05 | 2.26E-04 |
| | | A8HQ81 | RPL11 | Ribosomal protein L11 | 5.40 | 1.87 | 1.11 | 5.51E-04 | 3.92E-04 | 7.82E-01 |
| | | A8J597 | RPL12 | Ribosomal protein L12 | 1.75 | 0.47 | 0.25 | 2.96E-01 | 4.09E-03 | 5.09E-03 |
| | | A8HS59 | RPL17 | Ribosomal protein L17 | 3.66 | 0.97 | 0.46 | 3.89E-02 | 8.94E-01 | 1.28E-01 |

| | | | | | | | | |
|---|---|---|---|---|---|---|---|---|
| A8IA18 | RPL19 | Ribosomal protein L19 | 0.50 | 0.11 | 0.05 | 6.68E-03 | 1.80E-05 | 1.84E-09 |
| A8J951 | RPL21 | Ribosomal protein L21 | 3.97 | 1.37 | 0.41 | 4.16E-02 | 4.04E-01 | 1.38E-01 |
| A8J239 | RPL23a | Ribosomal protein L23a | 1.10 | 0.29 | 0.13 | 7.90E-01 | 9.73E-05 | 2.66E-05 |
| A8J1A3 | RPL24 | Ribosomal protein L24 | 1.02 | 0.44 | 0.14 | 9.46E-01 | 2.84E-05 | 1.12E-06 |
| A8I2T0 | RPL27a | Ribosomal protein L27a | 0.45 | 0.11 | 0.04 | 1.81E-03 | 3.91E-07 | 2.59E-09 |
| A8ID84 | RPL3 | Ribosomal protein L3 | 0.40 | 0.18 | 0.04 | 2.42E-03 | 4.92E-08 | 5.94E-09 |
| A8ICT1 | RPL30 | Ribosomal protein L30 | 2.73 | 0.80 | 0.38 | 2.29E-01 | 2.64E-01 | 2.76E-02 |
| A8ILG8 | RPL31 | Ribosomal protein L31 | 0.98 | 0.22 | 0.11 | 9.63E-01 | 4.25E-04 | 3.86E-05 |
| A8J2G4 | RPL32 | Ribosomal protein L32 | 1.13 | 0.31 | 0.14 | 7.85E-01 | 6.98E-04 | 2.82E-04 |
| A8J8P4 | RPL34 | Ribosomal protein L34 | 0.38 | 0.23 | 0.12 | 3.76E-04 | 2.98E-07 | 8.59E-05 |
| A8HNX3 | RPL35 | Ribosomal protein L35 | 8.38 | 2.40 | 1.59 | 7.12E-03 | 1.92E-02 | 4.00E-01 |
| A8HY08 | RPL37a | Ribosomal protein L37a | 0.71 | 0.37 | 0.18 | 1.26E-01 | 2.10E-03 | 1.43E-05 |
| A8J0I0 | RPL4 | Ribosomal protein L4 | 0.49 | 0.18 | 0.06 | 1.20E-02 | 2.49E-06 | 2.43E-07 |
| A8HP55 | RPL5 | Ribosomal protein L5 | 2.95 | 1.47 | 0.69 | 2.45E-02 | 3.91E-02 | 3.78E-01 |
| A8HP90 | RPL6 | Ribosomal protein L6 | 2.22 | 0.67 | 0.39 | 1.37E-01 | 2.76E-02 | 2.04E-02 |
| A8J567 | RPL7a | Ribosomal protein L7a | 6.85 | 2.95 | 1.22 | 6.39E-02 | 4.38E-02 | 7.86E-01 |
| A8JHC3 | RPS11 | Ribosomal protein S11 | 0.27 | 0.12 | 0.04 | 1.50E-05 | 2.75E-07 | 2.03E-07 |
| A8IGY1 | RPS13 | Ribosomal protein S13 | 1.48 | 0.36 | 0.09 | 4.44E-01 | 3.24E-05 | 4.92E-10 |
| A8J493 | RPS15 | Ribosomal protein S15 | 2.67 | 0.92 | 0.34 | 8.80E-02 | 7.74E-01 | 8.07E-03 |
| A8JE07 | RPS15a | Ribosomal protein S15a | 2.34 | 0.85 | 0.30 | 2.48E-02 | 5.27E-01 | 3.07E-03 |
| A8JGK1 | RPS17 | Ribosomal protein S17 | 27.56 | 11.70 | 5.88 | 2.49E-02 | 1.30E-02 | 3.15E-01 |
| A8HVP2 | RPS18 | Ribosomal protein S18 | 20.38 | 9.99 | 5.76 | 2.46E-02 | 1.87E-02 | 1.48E-01 |
| A8I403 | RPS19 | Ribosomal protein S19 | 11.02 | 4.78 | 2.51 | 3.40E-03 | 5.98E-03 | 2.79E-01 |
| A8HME4 | RPS2 | Ribosomal protein S2 | 0.98 | 0.32 | 0.10 | 8.87E-01 | 8.20E-07 | 1.72E-05 |
| A8J8M9 | RPS20 | Ribosomal protein S20 | 1.26 | 0.39 | 0.16 | 4.47E-01 | 1.46E-09 | 4.01E-04 |
| A8IS22 | RPS26 | Ribosomal protein S26 | 0.67 | 0.18 | 0.09 | 2.23E-01 | 2.60E-04 | 6.17E-06 |
| A8HVK4 | RPS27a | Ribosomal protein S27a | 0.22 | 0.07 | 0.03 | 2.28E-03 | 2.65E-07 | 1.02E-07 |
| A8IKP1 | RPS28 | Ribosomal protein S28 | 9.86 | 7.71 | 4.05 | 1.14E-01 | 2.08E-01 | 4.77E-02 |
| A8I4P5 | RPS3 | Ribosomal protein S3 | 1.62 | 0.42 | 0.18 | 2.89E-01 | 3.64E-04 | 4.32E-04 |
| A8JF66 | RPS30 | Ribosomal protein S30 | 0.75 | 0.27 | 0.12 | 1.63E-01 | 3.19E-04 | 6.60E-05 |
| A8IMP6 | RPS4 | Ribosomal protein S4 | 1.53 | 0.29 | 0.06 | 4.73E-01 | 2.58E-06 | 4.68E-08 |
| A8J2I5 | RPS5 | Ribosomal protein S5 | 16.37 | 10.13 | 6.52 | 1.79E-02 | 2.01E-02 | 1.56E-01 |
| A8HYC0 | UTP1 | Nucleolar protein component of the U3 processome | 1.13 | 0.70 | 0.86 | 8.20E-01 | 1.77E-02 | 6.70E-01 |
| A8JHU2 | RPL36 | 60S ribosomal protein L36 | 6.91 | 2.24 | 0.85 | 4.49E-02 | 2.38E-01 | 7.66E-01 |
| A8J5Z0 | RPP0 | Acidic ribosomal protein P0 | 1.08 | 0.36 | 0.09 | 7.59E-01 | 1.37E-03 | 5.04E-07 |
| A8J0R4 | RPP2 | Acidic ribosomal protein P2 | 3.56 | 3.63 | 3.14 | 2.96E-02 | 3.15E-02 | 1.08E-01 |
| A8IML7 | EIF3X | Hypothetical translation initiation factor (Fragment) | 3.17 | 1.70 | 0.87 | 4.27E-02 | 1.87E-01 | 6.67E-01 |
| A8I982 | RPL15 | Ribosomal protein L15 | 1.15 | 0.34 | 0.07 | 6.81E-01 | 1.49E-04 | 8.61E-07 |

| | | | | | | | | |
|---|---|---|---|---|---|---|---|---|
| A8JI94 | RPL22 | Ribosomal protein L22 | 21.08 | 11.10 | 6.18 | 1.75E-03 | 5.70E-03 | 3.78E-02 |
| A8HMG7 | RPL26 | Ribosomal protein L26 | 2.04 | 0.42 | 0.14 | 6.81E-02 | 6.18E-03 | 8.88E-06 |
| A8I0Y2 | RPL35a | Ribosomal protein L35a | 1.10 | 0.29 | 0.08 | 6.18E-01 | 1.69E-06 | 2.89E-07 |
| A8IVE2 | RPL7 | Ribosomal protein L7 | 10.68 | 5.29 | 2.56 | 4.76E-04 | 2.87E-04 | 1.39E-01 |
| A8IVK1 | RPL8 | Ribosomal protein L8 | 0.49 | 0.20 | 0.13 | 4.86E-02 | 3.26E-05 | 8.57E-05 |
| A8ILL5 | EIF2Ab | Eukaryotic initiation factor (Fragment) | 5.54 | 3.11 | 2.30 | 2.26E-02 | 9.66E-04 | 1.91E-01 |
| A8JHM2 | EIF3C | Eukaryotic initiation factor (Fragment) | 0.57 | 0.33 | 0.20 | 6.46E-02 | 1.10E-04 | 2.20E-04 |
| A8IP17 | CHLREDRAFT_188942 | Eukaryotic initiation factor 4A-like protein | 0.60 | 0.54 | 0.15 | 1.64E-02 | 4.55E-05 | 4.34E-05 |
| A8J9A9 | EIF3F | Eukaryotic initiation factor | 2.07 | 1.38 | 1.26 | 2.32E-02 | 2.60E-01 | 4.69E-01 |
| A8J4L1 | CHLREDRAFT_119356 | Predicted protein (Fragment) | 0.64 | 0.58 | 0.37 | 3.57E-02 | 3.74E-02 | 5.58E-03 |
| A8J2I8 | CHLREDRAFT_104111 | Predicted protein (Fragment) | 0.85 | 0.74 | 0.44 | 1.17E-01 | 2.99E-01 | 3.67E-02 |
| A8IBY2 | CHLREDRAFT_111269 | Predicted protein (Fragment) | 0.86 | 0.43 | 0.34 | 4.01E-01 | 4.51E-03 | 1.86E-03 |
| A8HTK2 | CPLD10 | Predicted protein | 0.78 | 0.66 | 0.67 | 4.35E-03 | 1.07E-02 | 7.41E-02 |
| Q7YKX3 | rps11 | 30S ribosomal protein S11 chloroplastic | 0.41 | 0.29 | 0.26 | 9.93E-04 | 2.08E-04 | 3.06E-03 |
| Q9GGE2 | rps14 | 30S ribosomal protein S14 chloroplastic | 2.17 | 1.07 | 1.02 | 4.56E-06 | 7.42E-01 | 5.00E-01 |
| O20032 | rps18 | 30S ribosomal protein S18 chloroplastic | 2.90 | 0.34 | 0.27 | 3.01E-02 | 1.50E-02 | 4.69E-03 |
| P59776 | rps19 | 30S ribosomal protein S19 chloroplastic | 0.38 | 0.11 | 0.17 | 5.15E-02 | 5.17E-06 | 9.81E-04 |
| O47027 | rps2-1 | 30S ribosomal protein S2 chloroplastic | 0.25 | 0.16 | 0.12 | 3.46E-04 | 3.76E-06 | 7.48E-05 |
| Q08365 | rps3 | 30S ribosomal protein S3 chloroplastic | 0.33 | 0.12 | 0.13 | 8.56E-04 | 3.88E-06 | 9.15E-05 |
| P48270 | rps4 | 30S ribosomal protein S4 chloroplastic | 0.96 | 0.29 | 0.14 | 6.97E-01 | 3.80E-04 | 3.30E-06 |
| P48267 | rps7 | 30S ribosomal protein S7 chloroplastic | 4.71 | 1.80 | 1.70 | 3.25E-03 | 2.77E-02 | 1.03E-01 |
| P59775 | rps8 | 30S ribosomal protein S8 chloroplastic | 1.21 | 0.59 | 0.52 | 6.18E-01 | 1.93E-02 | 1.60E-01 |
| O20029 | rps9 | 30S ribosomal protein S9 chloroplastic | 0.31 | 0.17 | 0.11 | 7.67E-04 | 7.22E-06 | 1.59E-05 |
| P11094 | rpl14 | 50S ribosomal protein L14 chloroplastic | 0.68 | 0.12 | 0.20 | 3.22E-02 | 1.41E-08 | 1.75E-03 |
| Q8HTL2 | rpl2 | 50S ribosomal protein L2 chloroplastic | 0.19 | 0.05 | 0.05 | 2.04E-04 | 3.55E-07 | 1.83E-07 |
| P26565 | rpl20 | 50S ribosomal protein L20 chloroplastic | 1.82 | 0.27 | 0.15 | 2.06E-01 | 8.73E-04 | 5.41E-06 |
| Q8HTL3 | rpl23 | 50S ribosomal protein L23 chloroplastic | 0.76 | 0.12 | 0.13 | 9.08E-02 | 7.08E-06 | 1.07E-04 |
| A8J3Z3 | PRPL31 | 50S ribosomal protein L31 | 0.34 | 0.08 | 0.15 | 1.30E-05 | 1.49E-07 | 1.26E-04 |
| A8JEP1 | PRPL35 | 50S ribosomal protein L35 | 1.33 | 0.49 | 0.22 | 4.85E-01 | 4.83E-02 | 9.30E-05 |
| P59774 | rpl36 | 50S ribosomal protein L36 chloroplastic | 0.20 | 0.11 | 0.06 | 2.14E-03 | 1.78E-05 | 2.13E-05 |
| Q8HTL1 | rpl5 | 50S ribosomal protein L5 chloroplastic | 2.79 | 0.68 | 0.37 | 1.09E-02 | 6.66E-02 | 1.61E-03 |
| A8I8Z4 | PRPL1 | Plastid ribosomal protein L1 | 0.40 | 0.06 | 0.04 | 8.41E-04 | 8.88E-07 | 7.65E-10 |
| A8ICE4 | PRPL11 | Plastid ribosomal protein L11 | 1.22 | 0.19 | 0.21 | 4.61E-01 | 2.52E-05 | 1.48E-05 |
| A8HWZ6 | PRPL13 | Plastid ribosomal protein L13 | 0.32 | 0.08 | 0.08 | 3.65E-04 | 1.59E-08 | 4.22E-07 |
| A8JAL6 | PRPL15 | Plastid ribosomal protein L15 | 0.24 | 0.04 | 0.05 | 3.39E-04 | 2.86E-07 | 2.65E-07 |
| A8I3M4 | PRPL17 | Plastid ribosomal protein L17 | 1.34 | 0.27 | 0.59 | 2.95E-02 | 3.70E-04 | 2.55E-02 |
| A8HNJ8 | PRPL18 | Plastid ribosomal protein L18 | 0.49 | 0.08 | 0.07 | 2.13E-02 | 2.02E-09 | 9.00E-08 |
| A8IW44 | PRPL19 | Plastid ribosomal protein L19 | 0.48 | 0.10 | 0.15 | 9.39E-03 | 1.18E-07 | 1.67E-04 |

| | | | | | | | | |
|---|---|---|---|---|---|---|---|---|
| A8IP00 | PRPL21 | Plastid ribosomal protein L21 | 0.54 | 0.19 | 0.37 | 3.92E-02 | 5.71E-03 | 5.38E-02 |
| A8J9D9 | PRPL24 | Plastid ribosomal protein L24 | 1.49 | 0.23 | 0.24 | 4.87E-01 | 2.09E-05 | 2.94E-04 |
| A8INR7 | PRPL27 | Plastid ribosomal protein L27 | 0.26 | 0.14 | 0.15 | 8.50E-04 | 8.24E-05 | 9.15E-05 |
| A8HWS8 | PRPL28 | Plastid ribosomal protein L28 | 0.21 | 0.05 | 0.06 | 5.06E-04 | 3.61E-07 | 1.30E-06 |
| A8JE35 | PRPL3 | Plastid ribosomal protein L3 | 0.23 | 0.04 | 0.03 | 2.72E-04 | 4.87E-08 | 4.45E-10 |
| A8IUC3 | PRPL32 | Plastid ribosomal protein L32 | 2.41 | 0.35 | 0.33 | 1.33E-01 | 5.31E-03 | 4.08E-03 |
| A8I1D3 | PRPL33 | Plastid ribosomal protein L33 | 1.03 | 0.41 | 0.43 | 7.78E-01 | 5.97E-05 | 4.32E-03 |
| Q84U22 | PRPL4 | Plastid ribosomal protein L4 | 2.76 | 0.59 | 0.40 | 2.10E-01 | 1.84E-02 | 7.93E-02 |
| A8J503 | PRPL6 | Plastid ribosomal protein L6 | 0.94 | 0.16 | 0.15 | 8.44E-01 | 3.37E-05 | 6.67E-05 |
| A8HTY0 | PRPL7/L12 | Plastid ribosomal protein L7/L12 | 3.16 | 1.76 | 1.34 | 1.35E-02 | 7.12E-03 | 2.51E-01 |
| A8IYS1 | PRPL9 | Plastid ribosomal protein L9 | 0.79 | 0.12 | 0.07 | 5.29E-01 | 1.65E-06 | 7.78E-10 |
| A8JGS2 | PRPS17 | Plastid ribosomal protein S17 | 0.45 | 0.08 | 0.09 | 3.05E-02 | 2.19E-10 | 3.44E-06 |
| A8JDN4 | PRPS20 | Plastid ribosomal protein S20 | 6.41 | 2.20 | 3.12 | 3.86E-04 | 8.18E-02 | 1.28E-02 |
| A8J8M5 | PRPS5 | Plastid ribosomal protein S5 | 0.18 | 0.05 | 0.07 | 1.71E-05 | 9.97E-09 | 2.59E-07 |
| A8IMN3 | PSRP-6 | Plastid-specific ribosomal protein 6 | 0.11 | 0.05 | 0.05 | 3.79E-05 | 1.07E-07 | 6.60E-08 |
| Q8HUH1 | rps2-2 | Putative 30S ribosomal S2-like protein | 1.87 | 1.15 | 1.08 | 1.02E-07 | 4.75E-01 | 7.72E-01 |
| A8IA39 | EFG1 | Chloroplast elongation factor G | 0.36 | 0.45 | 0.15 | 1.35E-03 | 6.65E-03 | 9.62E-05 |
| A8HTK7 | TBA1 | PsbA translation factor | 1.05 | 0.73 | 0.51 | 7.33E-01 | 9.95E-02 | 5.55E-03 |
| A8IPJ0 | MITC11 | Mitochondrial carrier protein | 0.50 | 0.67 | 0.47 | 3.18E-03 | 8.80E-02 | 1.49E-04 |
| A8IXI7 | MITC10 | Mitochondrial carrier protein | 0.72 | 0.93 | 0.76 | 2.70E-02 | 3.64E-01 | 3.24E-02 |
| A8HXM1 | MRPL29 | Mitochondrial ribosomal protein L29 | 9.54 | 2.56 | 2.79 | 3.83E-02 | 2.01E-01 | 2.66E-01 |
| A8HPX6 | MRPL3 | Mitochondrial ribosomal protein L3 | 0.86 | 0.67 | 0.54 | 1.18E-01 | 3.64E-04 | 7.34E-05 |
| A8JFK9 | MRPL7/L12 | Mitochondrial ribosomal protein L7/L12 | 2.68 | 2.68 | 2.28 | 0.00E+00 | 1.84E-04 | 1.99E-03 |
| A8JCF4 | MRPS17 | Mitochondrial ribosomal protein S17 | 0.70 | 0.44 | 0.50 | 3.46E-02 | 1.94E-03 | 6.40E-02 |
| A8JGX6 | MRPS6 | Mitochondrial ribosomal protein S6 | 0.76 | 0.75 | 0.87 | 1.39E-01 | 4.84E-02 | 6.05E-01 |
| A8J1X0 | UCP1 | Uncoupling protein | 0.62 | 0.63 | 0.51 | 3.78E-04 | 1.19E-02 | 7.34E-03 |
| A8HXY9 | UCP2 | Uncoupling protein | 2.05 | 2.44 | 2.27 | 1.36E-01 | 1.39E-02 | 1.61E-02 |
| A8J3F7 | MITC14 | Mitochondrial substrate carrier protein | 0.67 | 0.34 | 0.27 | 2.05E-01 | 3.24E-03 | 2.61E-03 |
| A8J2A5 | MSCP1 | Mitochondrial substrate carrier protein | 0.60 | 0.35 | 0.52 | 3.02E-02 | 4.42E-04 | 3.19E-02 |
| A8J1B6 | ANT1 | Adenine nucleotide translocator | 0.49 | 0.48 | 0.36 | 3.79E-02 | 1.79E-02 | 5.40E-04 |
| A8ID84 | RPL3 | Ribosomal protein L3 | 0.40 | 0.18 | 0.04 | 2.42E-03 | 4.92E-08 | 5.94E-09 |
| A8HP55 | RPL5 | Ribosomal protein L5 | 2.95 | 1.47 | 0.69 | 2.45E-02 | 3.91E-02 | 3.78E-01 |
| A8HP90 | RPL6 | Ribosomal protein L6 | 2.22 | 0.67 | 0.39 | 1.37E-01 | 2.76E-02 | 2.04E-02 |
| A8HQ81 | RPL11 | Ribosomal protein L11 | 5.40 | 1.87 | 1.11 | 5.51E-04 | 3.92E-04 | 7.82E-01 |
| A8J597 | RPL12 | Ribosomal protein L12 | 1.75 | 0.47 | 0.25 | 2.96E-01 | 4.09E-03 | 5.09E-03 |
| P26565 | rpl20 | 50S ribosomal protein L20 chloroplastic | 1.82 | 0.27 | 0.15 | 2.06E-01 | 8.73E-04 | 5.41E-06 |
| Q8HTL3 | rpl23 | 50S ribosomal protein L23 chloroplastic | 0.76 | 0.12 | 0.13 | 9.08E-02 | 7.08E-06 | 1.07E-04 |
| A8J239 | RPL23a | Ribosomal protein L23a | 1.10 | 0.29 | 0.13 | 7.90E-01 | 9.73E-05 | 2.66E-05 |

| | | | | | | | | | | |
|---|---|---|---|---|---|---|---|---|---|---|
| | | A8ICE4 | PRPL11 | Plastid ribosomal protein L11 | 1.22 | 0.19 | 0.21 | 4.61E-01 | 2.52E-05 | 1.48E-05 |
| | | Q8HTL1 | rpl5 | 50S ribosomal protein L5 chloroplastic | 2.79 | 0.68 | 0.37 | 1.09E-02 | 6.66E-02 | 1.61E-03 |
| | | A8J1A3 | RPL24 | Ribosomal protein L24 | 1.02 | 0.44 | 0.14 | 9.46E-01 | 2.84E-05 | 1.12E-06 |
| | | A8IZK3 | RPL10 | Ribosomal protein L10 | 1.19 | 0.30 | 0.12 | 5.62E-01 | 3.35E-05 | 2.26E-04 |
| | | A8I2G3 | CHLREDRAFT_112251 | Predicted protein | 0.57 | 0.48 | 0.51 | 3.46E-03 | 2.43E-03 | 5.58E-02 |
| | | A8HXD0 | CHLREDRAFT_134139 | Predicted protein | 0.96 | 0.70 | 0.72 | 3.27E-01 | 7.35E-03 | 1.42E-01 |
| | | A8J8J8 | CHLREDRAFT_120661 | GTPase Der | 0.44 | 0.43 | 0.44 | 1.68E-03 | 5.38E-04 | 5.25E-02 |
| | | A8HZV5 | CHLREDRAFT_187641 | Predicted protein | 0.50 | 0.46 | 0.47 | 3.73E-02 | 5.22E-02 | 1.87E-03 |
| 2 | Photosynthesis | P10898 | psbC | Photosystem II CP43 chlorophyll apoprotein | 0.31 | 2.01 | 1.38 | 2.50E-03 | 1.29E-01 | 5.10E-01 |
| | | P06007 | psbD | Photosystem II D2 protein | 0.12 | 1.14 | 1.02 | 2.97E-07 | 2.99E-01 | 9.28E-01 |
| | | P22666 | psbH | Photosystem II reaction center protein H | 12.59 | 15.21 | 12.10 | 5.62E-04 | 1.21E-02 | 1.03E-02 |
| | | Q06480 | psbN | Protein psbN | 1.39 | 1.02 | 1.13 | 3.73E-03 | 8.68E-01 | 4.89E-04 |
| | | A8J0E4 | PsbO | Oxygen-evolving enhancer protein 1 of photosystem II | 0.94 | 0.77 | 0.69 | 2.48E-01 | 2.71E-05 | 1.94E-06 |
| | | A8IXU9 | CGL30/PsbP | Photosystem II thylakoid lumenal 29.8 kDa protein PsbP | 7.48 | 7.99 | 7.70 | 5.69E-02 | 9.00E-02 | 4.35E-02 |
| | | A8IYH9 | PsbP1 | Oxygen-evolving enhancer protein 2 of photosystem II | 1.50 | 0.74 | 0.56 | 3.85E-01 | 4.36E-02 | 8.67E-02 |
| | | A8J3S9 | PsbP2 | PsbP-like protein | 1.01 | 0.51 | 0.67 | 9.11E-01 | 5.26E-02 | 4.86E-03 |
| | | A8IKE6 | PsbP3 | OEE2-like protein of thylakoid lumen | 1.05 | 0.42 | 0.36 | 8.85E-01 | 6.81E-04 | 1.04E-04 |
| | | A8IHH9 | PsbP6 | Lumen targeted protein | 1.12 | 0.59 | 0.43 | 6.13E-01 | 1.36E-02 | 1.46E-02 |
| | | A8JEV1 | PsbQ | Oxygen evolving enhancer protein 3 | 13.41 | 14.63 | 11.40 | 1.15E-02 | 5.20E-04 | 1.08E-03 |
| | | A8HXG5 | PsbR | 10 kDa photosystem II polypeptide | 0.78 | 0.42 | 0.27 | 2.42E-02 | 9.25E-03 | 1.62E-02 |
| | | A8JFQ7 | PsbW | Photosystem II reaction center W protein | 0.17 | 0.08 | 0.10 | 4.09E-05 | 1.33E-05 | 5.05E-05 |
| | | A8III5 | Psb28 | Photosystem II reaction center psb28 protein | 0.67 | 0.19 | 0.28 | 1.97E-03 | 9.04E-05 | 2.27E-05 |
| | | P12154 | psaA | Photosystem I P700 chlorophyll a apoprotein A1 | 0.08 | 3.02 | 3.59 | 2.68E-06 | 1.18E-02 | 3.27E-02 |
| | | P09144 | psaB | Photosystem I P700 chlorophyll a apoprotein A2 | 0.13 | 4.23 | 4.34 | 1.93E-05 | 5.52E-03 | 5.32E-03 |
| | | Q5NKW4 | PsaD | Photosystem I reaction center subunit II 20 kDa | 14.68 | 19.73 | 20.90 | 1.38E-03 | 1.29E-04 | 2.92E-07 |
| | | A8J4S1 | PsaF | Photosystem I reaction center subunit III | 2.31 | 0.99 | 0.69 | 1.59E-01 | 9.86E-01 | 3.07E-02 |
| | | A8JHN9 | PsaG | Photosystem I reaction center subunit V | 0.30 | 0.10 | 0.15 | 1.32E-03 | 2.72E-08 | 2.65E-05 |
| | | A8IH77 | PsaH | Subunit H of photosystem I | 12.48 | 9.65 | 9.76 | 1.05E-02 | 3.24E-02 | 1.49E-02 |
| | | P59777 | psaJ | Photosystem I reaction center subunit IX | 0.26 | 0.12 | 0.23 | 2.80E-06 | 1.73E-04 | 1.62E-02 |
| | | A8J6K8 | PsaK | Photosystem I reaction center subunit psaK | 0.51 | 0.18 | 0.12 | 2.34E-03 | 4.70E-05 | 5.15E-06 |
| | | A8I835 | PsaN | Photosystem I reaction center subunit N | 0.47 | 0.41 | 0.08 | 9.95E-04 | 3.27E-04 | 8.99E-07 |
| | | O20030 | ycf4 | Photosystem I assembly protein Ycf4 | 0.08 | 0.08 | 0.14 | 2.20E-08 | 6.55E-08 | 2.94E-06 |
| | | A8J9G6 | Cyc 6 | Cytochrome c6 | 3.06 | 0.46 | 1.20 | 3.13E-01 | 3.89E-02 | 8.36E-01 |
| | | P23577 | petA | Apocytochrome f | 1.77 | 0.71 | 0.91 | 2.57E-01 | 2.18E-02 | 5.15E-01 |
| | | Q00471 | petB | Cytochrome b6 | 0.27 | 0.42 | 0.61 | 2.97E-04 | 4.23E-05 | 2.67E-03 |
| | | A8IXV0 | CHLRE_03g198850v5 | Thylakoid lumen protein | 1.29 | 0.78 | 0.45 | 2.62E-01 | 2.74E-01 | 1.50E-02 |
| | | A8HZ72 | CGLD14 | Predicted protein | 0.48 | 0.18 | 0.38 | 1.92E-02 | 5.84E-04 | 2.94E-02 |
| | | A8HQJ5 | TEF30 | Predicted protein | 0.85 | 0.71 | 0.79 | 1.81E-01 | 7.00E-03 | 6.76E-02 |

| | | | | | | | | |
|---|---|---|---|---|---|---|---|---|
| P00877 | Rbcl | Ribulose bisphosphate carboxylase large chain | 0.54 | 0.56 | 0.37 | 8.04E-02 | 7.53E-02 | 4.90E-02 |
| A8IYP4 | PRK1 | Phosphoribulokinase | 1.04 | 0.12 | 0.11 | 8.34E-01 | 2.66E-06 | 1.77E-09 |
| A8IKQ0 | FBP1 | Fructose-1,6-bisphosphatase | 0.66 | 0.28 | 0.07 | 1.41E-05 | 1.39E-04 | 5.99E-08 |
| A8I531 | CHLD | Magnesium chelatase subunit D | 0.20 | 0.15 | 0.12 | 9.07E-05 | 2.18E-06 | 3.19E-07 |
| A8IMZ5 | CHLI1 | Magnesium chelatase subunit I | 0.12 | 0.07 | 0.06 | 2.84E-09 | 5.58E-09 | 1.21E-07 |
| A8IKQ6 | CHLI2 | Magnesium chelatase subunit I | 0.18 | 0.09 | 0.16 | 5.76E-05 | 2.07E-07 | 7.13E-07 |
| A8IKK7 | CTH1\|CTH1B | Copper target 1 protein | 0.31 | 0.33 | 0.38 | 1.20E-02 | 1.50E-02 | 1.75E-02 |
| A8ITX0 | CRD1 | Copper response defect 1 protein | 1.55 | 0.40 | 0.60 | 7.13E-01 | 4.04E-02 | 4.62E-01 |
| A8HNE8 | CHLRE_01g050950v5 | Geranylgeranyl reductase | 0.23 | 0.13 | 0.09 | 5.09E-04 | 7.99E-06 | 2.98E-06 |
| A8HPJ2 | POR | Light-dependent protochlorophyllide reductase | 0.20 | 0.09 | 0.05 | 5.50E-05 | 1.39E-06 | 6.02E-12 |
| Q9AXF6 | LHCBM7 | Chlorophyll a-b binding protein of LHCII | 0.69 | 0.22 | 0.95 | 3.29E-04 | 1.81E-03 | 2.58E-02 |
| Q9ZSJ4 | LHCBM5 | Chlorophyll a-b binding protein of LHCII | 3.42 | 3.47 | 3.55 | 1.68E-02 | 1.06E-01 | 1.90E-03 |
| A8JCU4 | LHCBM1 | Chlorophyll a-b binding protein of LHCII | 4.45 | 5.59 | 5.91 | 1.40E-02 | 1.32E-01 | 1.36E-02 |
| A8J6D1 | CP29 | Chlorophyll a-b binding protein of photosystem II | 0.78 | 0.44 | 0.18 | 3.47E-01 | 2.26E-02 | 4.46E-07 |
| A8J287 | LHCBM6 | Chloropyll a-b binding protein of LHCII type I chloroplast | 0.79 | 0.53 | 0.59 | 1.77E-01 | 3.93E-02 | 4.87E-02 |
| A8J270 | LHCBM8 | Chlorophyll a-b binding protein of LHCII | 0.59 | 0.32 | 0.44 | 1.69E-01 | 5.95E-03 | 3.24E-02 |
| Q8S3T9 | LHCBM9 | Chlorophyll a-b binding protein of LHCII | 24.82 | 23.50 | 26.59 | 7.94E-05 | 4.68E-03 | 1.40E-04 |
| P93664 | LHCSR1 | Light-harvesting complex stress-related protein 1, chloroplastic | 9.42 | 28.18 | 39.15 | 5.31E-02 | 1.52E-02 | 1.49E-04 |
| A8J431 | LHCSR3 | Stress-related chlorophyll a/b binding protein 2 | 0.28 | 0.22 | 0.50 | 1.67E-03 | 4.82E-04 | 1.09E-01 |
| A8JF10 | LHCA3 | Light-harvesting chlorophyll-a/b protein of photosystem I type III | 0.72 | 0.61 | 0.59 | 3.46E-01 | 4.66E-02 | 1.45E-02 |
| A8I000 | LHCA4 | Light-harvesting protein of photosystem I | 4.15 | 4.68 | 5.35 | 5.31E-03 | 1.24E-01 | 1.07E-02 |
| A8ISG0 | LHCA7 | Light-harvesting protein of photosystem I | 0.55 | 0.44 | 0.70 | 3.43E-03 | 1.35E-02 | 5.96E-02 |
| A8ITV3 | LHCA9 | Light-harvesting protein of photosystem I | 0.62 | 0.34 | 0.48 | 9.44E-06 | 8.72E-04 | 1.61E-03 |
| Q75VY8 | LHCA5 | Light-harvesting chlorophyll-a/b protein of photosystem I | 0.62 | 0.65 | 1.12 | 1.18E-02 | 7.37E-02 | 6.31E-01 |
| Q75VY7 | LHCA8 | Light-harvesting chlorophyll-a/b protein of photosystem I | 2.54 | 2.27 | 3.87 | 1.04E-01 | 2.65E-01 | 2.54E-02 |
| A8J0A7 | ELI3 | Early light-inducible protein | 1.32 | 2.92 | 3.33 | 7.03E-01 | 3.19E-02 | 3.73E-02 |
| A8HMS2 | CAO | Chlorophyll a oxygenase | 0.78 | 0.43 | 0.65 | 6.75E-01 | 1.26E-02 | 3.06E-01 |
| A8JFJ1 | CHLG | Chlorophyll synthetase | 0.40 | 0.65 | 0.74 | 2.28E-02 | 1.03E-01 | 1.47E-01 |
| A8J7H3 | GSA | Chlorophyll synthetase | 0.06 | 0.04 | 0.06 | 1.27E-08 | 1.32E-09 | 1.23E-06 |
| A8I7P5 | CHLH1 | Magnesium chelatase subunit H | 0.06 | 0.12 | 0.14 | 6.32E-07 | 7.25E-06 | 3.51E-05 |
| A8JGJ6 | CHLM | Mg protoporphyrin IX S-adenosyl methionine O-methyl transferase | 0.28 | 0.18 | 0.42 | 7.01E-07 | 2.76E-04 | 2.98E-03 |
| A8J9S8 | PNP1 | Polyribonucleotide phosphorylase PNPase (Fragment) | 0.51 | 0.71 | 0.40 | 8.83E-04 | 3.27E-02 | 1.67E-04 |
| A8J9E9 | CHLREDRAFT_196597 | Carotenoid isomerase (Fragment) | 1.19 | 2.11 | 2.05 | 4.02E-01 | 1.55E-02 | 2.92E-03 |
| Q6J214 | PSY | Chloroplast phytoene synthase | 0.26 | 0.58 | 0.93 | 8.42E-04 | 5.61E-03 | 8.61E-01 |
| A8J261 | HST1 | Homogentisate solanesyltransferase (Fragment) | 0.42 | 0.53 | 0.49 | 4.70E-03 | 7.21E-03 | 2.44E-02 |
| A8HWI0 | LCYE | Lycopene epsilon cyclase | 0.14 | 0.21 | 0.38 | 3.51E-05 | 1.90E-04 | 1.49E-04 |
| A8I647 | ZDS1 | Zeta-carotene desaturase | 0.21 | 0.23 | 0.17 | 1.05E-05 | 1.01E-04 | 9.32E-06 |

| | | | | | | | | | | |
|---|---|---|---|---|---|---|---|---|---|---|
| 3 | Redox homeostasis | A8IUG8 | CDSP32 | Plastidic thioredoxin-like protein | 0.90 | 0.76 | 0.35 | 5.06E-01 | 6.09E-03 | 1.48E-04 |
| | | A8J0Q8 | CITRX | Thioredoxin-related protein | 4.51 | 4.51 | 3.24 | 4.33E-03 | 4.25E-03 | 1.16E-01 |
| | | A8J594 | DLC3 | Flagellar outer dynein arm 16 kDa light chain LC3 | 3.34 | 3.22 | 1.65 | 4.94E-02 | 1.52E-01 | 8.43E-02 |
| | | A8HPL8 | DLD2 | Dihydrolipoamide dehydrogenase | 0.70 | 0.59 | 0.28 | 7.27E-02 | 2.07E-02 | 1.66E-04 |
| | | A8J1T4 | GCSL | Dihydrolipoyl dehydrogenase | 2.31 | 4.07 | 4.23 | 5.44E-02 | 1.55E-02 | 7.15E-03 |
| | | A8JHA9 | GRX1 | Glutaredoxin CPYC type | 3.21 | 3.64 | 3.10 | 5.87E-04 | 3.04E-08 | 9.41E-04 |
| | | A8IYH1 | GRX2 | Glutaredoxin CPYC type | 1.71 | 2.45 | 3.47 | 9.28E-02 | 1.72E-01 | 4.41E-02 |
| | | A8J916 | CHLRE_06g278183v5 | Glutaredoxin-like protein | 1.12 | 2.03 | 2.25 | 3.76E-01 | 5.84E-02 | 2.53E-02 |
| | | A8JH05 | GRX3 | Glutaredoxin CGFS type | 0.80 | 0.64 | 0.59 | 1.53E-01 | 2.43E-03 | 9.55E-02 |
| | | A8JIA7 | GRX4 | Glutaredoxin CGFS type | 0.78 | 1.05 | 1.12 | 5.34E-03 | 8.55E-01 | 3.13E-02 |
| | | A8HN52 | GRX6 | Glutaredoxin CGFS type | 0.51 | 0.15 | 0.12 | 2.43E-02 | 4.70E-06 | 6.67E-05 |
| | | A8J0E5 | GSHR2 | Glutathione reductase | 1.56 | 2.78 | 2.33 | 1.84E-02 | 1.51E-04 | 3.36E-02 |
| | | A8J6A7 | MET16/APR1 | Adenylylphosphosulfate reductase | 8.99 | 18.99 | 24.48 | 1.92E-05 | 5.61E-04 | 7.93E-04 |
| | | A8J9N7 | NRX4 | Nucleoredoxin | 1.19 | 1.65 | 1.69 | 1.90E-01 | 3.56E-03 | 1.72E-01 |
| | | A8HNQ7 | NTRC1 | Thioredoxin reductase | 0.35 | 0.53 | 1.19 | 3.21E-03 | 1.57E-02 | 5.05E-01 |
| | | O48949 | PDI | Protein disulfide isomerase | 9.02 | 8.22 | 10.25 | 8.36E-02 | 1.05E-01 | 4.16E-02 |
| | | A8HQT1 | PDI2 | Protein disulfide isomerase | 6.88 | 5.12 | 2.64 | 1.51E-02 | 8.55E-02 | 3.44E-02 |
| | | A8JBH7 | PDI3 | Protein disulfide isomerase | 1.43 | 0.42 | 0.38 | 1.20E-01 | 3.44E-02 | 1.49E-04 |
| | | A8IHI1 | PDI4 | Protein disulfide isomerase | 0.58 | 2.13 | 1.91 | 1.57E-02 | 2.01E-01 | 7.67E-02 |
| | | A8HZQ4 | PRX3 | Peroxiredoxin type II | 0.74 | 0.54 | 0.43 | 9.78E-02 | 3.21E-03 | 1.71E-04 |
| | | A8HPG8 | PRX5 | Peroxiredoxin type II | 4.75 | 4.11 | 3.29 | 6.62E-02 | 2.90E-02 | 3.55E-02 |
| | | A8ICK6 | PRX6 | Thioredoxin dependent peroxidase | 0.73 | 0.42 | 0.44 | 1.95E-01 | 8.74E-04 | 2.03E-03 |
| | | A8JIT5 | PRX7 | Peroxiredoxin | 2.56 | 3.38 | 3.54 | 4.76E-02 | 4.43E-02 | 3.40E-02 |
| | | A8JFC3 | SCO1 | Cytochrome c oxidase assembly factor | 1.60 | 1.87 | 2.13 | 2.78E-02 | 2.44E-02 | 2.45E-02 |
| | | Q9FE86 | PRX1 | 2-cys peroxiredoxin chloroplastic | 17.92 | 17.27 | 17.88 | 5.92E-03 | 7.49E-05 | 7.60E-04 |
| | | A8IZR5 | TRXh | Thioredoxin | 9.35 | 9.45 | 11.36 | 2.54E-02 | 3.55E-02 | 4.68E-02 |
| | | A8HP58 | TRXm | Thioredoxin | 1.39 | 1.18 | 0.17 | 4.53E-01 | 6.23E-01 | 7.60E-05 |
| | | Q84XS0 | TRXo | Thioredoxin o | 1.29 | 0.97 | 0.20 | 7.17E-02 | 9.14E-01 | 6.66E-05 |
| | | Q84XR9 | TRXx | Thioredoxin x | 9.23 | 8.98 | 9.09 | 1.58E-01 | 1.62E-02 | 4.23E-02 |
| | | A8IQA9 | CHLREDRAFT_11164 | Thioredoxin-like protein | 1.06 | 0.75 | 0.92 | 8.41E-01 | 2.32E-03 | 7.00E-01 |
| | | A8I0I4 | CHLRE_10g456250v5 | Thioredoxin-like protein | 0.56 | 0.60 | 0.67 | 9.46E-04 | 5.40E-03 | 9.08E-04 |
| | | A8JA70 | CHLRE_16g687294v5 | Ferredoxin thioredoxin reductase variable chain | 2.08 | 0.55 | 0.34 | 7.57E-02 | 2.07E-04 | 9.15E-04 |
| | | A8HUI8 | CHLREDRAFT_188073 | Predicted protein | 1.08 | 1.62 | 2.00 | 6.05E-01 | 2.81E-01 | 2.57E-03 |
| | | A8JDA2 | CHLRE_17g715500v5 | Predicted protein | 2.01 | 1.36 | 0.90 | 2.56E-02 | 1.82E-01 | 1.40E-01 |
| | | A8IC32 | CHLREDRAFT_141005 | Predicted protein | 0.47 | 0.67 | 0.97 | 2.39E-02 | 1.56E-01 | 8.75E-01 |
| | | A8JG35 | CHLREDRAFT_160132 | Predicted protein | 0.96 | 0.63 | 0.28 | 2.10E-01 | 1.79E-03 | 9.24E-05 |
| | | A8HUF5 | CHLREDRAFT_182752 | Predicted protein | 1.05 | 0.92 | 0.52 | 8.00E-01 | 6.64E-01 | 1.04E-02 |
| | | A8IHN7 | CHLREDRAFT_188533 | Predicted protein | 0.60 | 1.07 | 1.14 | 2.39E-02 | 8.03E-01 | 7.10E-01 |

| | | | | | | | | | | |
|---|---|---|---|---|---|---|---|---|---|---|
| | | O49822 | apx1 | Ascorbate peroxidase | 4.57 | 5.70 | 5.27 | 2.90E-02 | 1.39E-04 | 1.03E-02 |
| | | A8J7X9 | CCPR1 | Cytochrome c peroxidase | 4.88 | 4.58 | 3.49 | 2.06E-03 | 5.60E-03 | 1.35E-02 |
| | | A8JG56 | CHLREDRAFT_165193 | L-ascorbate peroxidase | 1.26 | 0.62 | 0.14 | 1.43E-02 | 3.48E-03 | 1.58E-05 |
| | | A8J285 | APX2 | L-ascorbate peroxidase | 2.44 | 4.11 | 4.41 | 4.52E-02 | 1.69E-02 | 8.30E-02 |
| | | O81648 | Lci2 | Low CO2 inducible gene | 0.76 | 0.52 | 0.46 | 4.51E-01 | 7.40E-03 | 1.16E-02 |
| | | A8J537 | CAT1 | Catalase | 4.78 | 6.46 | 6.15 | 8.52E-03 | 6.37E-03 | 3.62E-03 |
| | | A8HN12 | CAT2 | Catalase/peroxidase | 2.23 | 3.01 | 3.26 | 8.57E-02 | 4.36E-02 | 2.14E-02 |
| | | A8IXD6 | CLPR4 | ATP-dependent Clp protease proteolytic subunit | 6.14 | 5.76 | 3.45 | 1.33E-02 | 8.21E-02 | 5.57E-03 |
| 4 | Protein folding | A8J524 | CCT2 | T-complex protein 1 beta subunit | 1.77 | 1.55 | 0.99 | 1.68E-03 | 2.18E-01 | 9.67E-01 |
| | | A8J7J2 | CCT5 | T-complex protein epsilon subunit | 1.23 | 2.58 | 1.74 | 4.37E-01 | 9.33E-03 | 2.36E-01 |
| | | A8HQ74 | CCT7 | T-complex protein eta subunit | 0.81 | 1.96 | 1.70 | 2.70E-01 | 3.16E-02 | 8.51E-02 |
| | | A8IF08 | CCT8 | T-complex protein theta subunit | 1.44 | 1.61 | 1.16 | 2.87E-01 | 1.81E-02 | 7.68E-01 |
| | | Q66YD3 | CDJ1 | Chloroplast DnaJ-like protein | 0.57 | 0.57 | 1.14 | 1.24E-03 | 2.13E-02 | 4.62E-01 |
| | | A8J594 | DLC3 | Flagellar outer dynein arm 16 kDa light chain LC3 | 3.34 | 3.22 | 1.65 | 4.94E-02 | 1.52E-01 | 8.43E-02 |
| | | A8IQC5 | DNJ1 | DnaJ-like protein | 0.31 | 1.13 | 1.39 | 1.23E-02 | 5.25E-01 | 3.77E-01 |
| | | A8HMC0 | CRT2 | Calreticulin 2 calcium-binding protein | 1.17 | 0.48 | 0.22 | 6.52E-01 | 4.65E-03 | 2.77E-05 |
| | | A8IV02 | CYN1a\|CYN1b | Peptidyl-prolyl cis-transisomerase cyclophilin type | 0.97 | 0.87 | 0.76 | 7.69E-01 | 3.07E-01 | 5.54E-03 |
| | | A8JD64 | CYN19-2 | Peptidyl-prolyl cis-trans isomerase | 0.88 | 0.51 | 0.29 | 6.04E-01 | 1.44E-02 | 1.23E-02 |
| | | A8HUU9 | CYN19-3 | Peptidyl-prolyl cis-trans isomerase | 0.62 | 0.69 | 0.34 | 8.03E-02 | 1.72E-01 | 1.52E-03 |
| | | A8J282 | CYN20-1 | Peptidyl-prolyl cis-trans isomerase | 2.93 | 5.91 | 2.59 | 2.41E-02 | 7.34E-04 | 7.15E-02 |
| | | A8JDL5 | CYN20-3 | Peptidyl-prolyl cis-trans isomerase | 1.18 | 0.60 | 0.51 | 7.68E-01 | 1.82E-02 | 2.59E-02 |
| | | A8ID98 | CYN20-5 | Peptidyl-prolyl cis-trans isomerase cyclophilin-type | 0.67 | 0.66 | 0.98 | 3.63E-03 | 1.61E-01 | 9.23E-01 |
| | | A8JFN4 | CYN23a\|CYN23b | Cyclophilin-like protein | 2.26 | 1.79 | 1.30 | 6.35E-02 | 1.13E-01 | 4.40E-02 |
| | | A8IE53 | CYN26 | Peptidyl-prolyl cis-trans isomerase cyclophilin-type | 0.65 | 0.35 | 0.38 | 7.37E-03 | 1.92E-04 | 5.52E-04 |
| | | A8JHN8 | CYN28 | Peptidyl-prolyl cis-trans isomerase cyclophilin-type | 0.95 | 0.46 | 0.30 | 6.72E-01 | 6.46E-03 | 2.90E-03 |
| | | A8INE5 | CYN37 | Peptidyl-prolyl cis-trans isomerase cyclophilin-type | 1.02 | 0.46 | 0.72 | 9.55E-01 | 2.30E-03 | 2.67E-01 |
| | | A8I0M0 | CYN65 | Peptidyl-prolyl cis-trans isomerase cyclophilin-type | 1.12 | 0.59 | 0.98 | 8.96E-02 | 3.69E-02 | 8.87E-01 |
| | | A8I6Y0 | ERJ1 | ER DnaJ-like protein 1 | 0.65 | 0.54 | 0.70 | 2.13E-02 | 5.09E-03 | 2.81E-05 |
| | | A8J3C1 | FKB16-7a\|FKB16-7b | Peptidyl-prolyl cis-trans isomerase | 0.61 | 0.29 | 0.35 | 3.54E-05 | 2.49E-04 | 2.07E-03 |
| | | A8J3L6 | FKB16-2a\|FKB16-2c\|FKB16-2b | Peptidyl-prolyl cis-trans isomerase | 1.22 | 0.73 | 0.45 | 1.29E-01 | 8.34E-02 | 1.91E-03 |
| | | A8I9C7 | FKB42 | Peptidyl-prolyl cis-trans isomerase FKBP-type | 1.32 | 1.77 | 1.47 | 5.84E-03 | 1.80E-03 | 4.82E-01 |
| | | A8J1U1 | HSP90A | Heat shock protein 90A | 3.10 | 4.59 | 3.90 | 7.22E-02 | 1.19E-02 | 5.60E-02 |
| | | A8I7T1 | HSP90B | Heat shock protein 90B | 2.24 | 2.85 | 3.28 | 9.60E-02 | 5.77E-03 | 6.40E-03 |
| | | Q66T67 | HSP90C | Heat shock protein 90C | 0.75 | 0.42 | 0.36 | 3.83E-01 | 3.92E-03 | 1.99E-03 |
| | | Q944P3 | HSP33 | Heat shock protein 33 | 0.54 | 0.32 | 0.62 | 7.90E-04 | 9.03E-04 | 1.65E-02 |
| | | A8JES1 | MGE1 | GrpE protein homolog | 3.31 | 2.63 | 1.05 | 8.42E-03 | 4.65E-03 | 8.55E-01 |
| | | A8HPB8 | MSRA4 | Peptidyl-prolyl cis-trans isomerase, FKBP-type | 0.76 | 0.38 | 0.19 | 2.49E-01 | 3.46E-07 | 2.35E-06 |
| | | A8JDH3 | NUDC | Nuclear movement family protein | 0.87 | 0.60 | 0.40 | 4.12E-01 | 1.43E-01 | 5.88E-03 |

| | | | | | | | | | | |
|---|---|---|---|---|---|---|---|---|---|---|
| | | A8JBH7 | PDI3 | Protein disulfide isomerase | 1.43 | 0.42 | 0.38 | 1.20E-01 | 3.44E-02 | 1.49E-04 |
| | | A8IHI1 | PDI4 | Protein disulfide isomerase | 0.58 | 2.13 | 1.91 | 1.57E-02 | 2.01E-01 | 7.67E-02 |
| | | A8JD56 | TIG1 | Chloroplast trigger factor | 0.54 | 0.26 | 0.30 | 1.57E-03 | 2.78E-07 | 4.16E-03 |
| | | A8IZR5 | TRXh | Thioredoxin | 9.35 | 9.45 | 11.36 | 2.54E-02 | 3.55E-02 | 4.68E-02 |
| | | A8HP58 | TRXm | Thioredoxin | 1.39 | 1.18 | 0.17 | 4.53E-01 | 6.23E-01 | 7.60E-05 |
| | | Q84XS0 | TRXo | Thioredoxin o | 1.29 | 0.97 | 0.20 | 7.17E-02 | 9.14E-01 | 6.66E-05 |
| | | Q84XR9 | TRXx | Thioredoxin x | 9.23 | 8.98 | 9.09 | 1.58E-01 | 1.62E-02 | 4.23E-02 |
| | | A8J0Q8 | CITRX | Thioredoxin-related protein CITRX | 4.51 | 4.51 | 3.24 | 4.33E-03 | 4.25E-03 | 1.16E-01 |
| | | A8IQA9 | CHLREDRAFT_11164 | Thioredoxin-like protein | 1.06 | 0.75 | 0.92 | 8.41E-01 | 2.32E-03 | 7.00E-01 |
| | | A8I0I4 | CHLRE_10g456250v5 | Thioredoxin-like protein | 0.56 | 0.60 | 0.67 | 9.46E-04 | 5.40E-03 | 9.08E-04 |
| | | A8I211 | CHLREDRAFT_161043 | Predicted protein | 1.84 | 2.17 | 1.99 | 7.90E-03 | 1.03E-02 | 9.29E-03 |
| | | A8IHN7 | CHLREDRAFT_188533 | Predicted protein | 0.60 | 1.07 | 1.14 | 2.39E-02 | 8.03E-01 | 7.10E-01 |
| | | A8HUK0 | FKB12 | Peptidyl-prolyl cis-trans isomerase | 0.81 | 0.29 | 0.23 | 3.94E-01 | 5.31E-06 | 3.19E-05 |
| | | A8J746 | FKB15-1 | Peptidyl-prolyl cis-trans isomerase | 0.66 | 0.31 | 0.23 | 2.28E-02 | 1.63E-04 | 2.25E-04 |
| | | A8J740 | FKB15-4 | Peptidyl-prolyl cis-trans isomerase | 0.64 | 0.13 | 0.13 | 1.12E-01 | 9.30E-06 | 3.53E-06 |
| | | A8I6B6 | FKB16-1 | Peptidyl-prolyl cis-trans isomerase | 0.77 | 0.33 | 0.29 | 4.16E-01 | 1.67E-05 | 4.69E-04 |
| | | A8J3L3 | FKB16-5 | Peptidyl-prolyl cis-trans isomerase | 0.75 | 0.42 | 0.17 | 1.23E-01 | 2.38E-05 | 4.12E-04 |
| | | A8JEI4 | FKB16-8 | Peptidyl-prolyl cis-trans isomerase | 0.76 | 0.37 | 0.29 | 3.34E-02 | 3.59E-05 | 7.49E-04 |
| | | A8J1E5 | FKB17-1 | Peptidyl-prolyl cis-trans isomerase | 0.69 | 0.37 | 0.40 | 1.90E-01 | 5.38E-03 | 2.19E-02 |
| | | A8JEK6 | FKB17-2 | Peptidyl-prolyl cis-trans isomerase | 0.51 | 0.73 | 0.68 | 3.17E-02 | 1.65E-01 | 2.50E-03 |
| | | A8I1U5 | FKB18 | Peptidyl-prolyl cis-trans isomerase | 0.91 | 0.37 | 0.37 | 6.19E-01 | 3.01E-09 | 1.07E-04 |
| | | A8IVN2 | FKB19 | Peptidyl-prolyl cis-trans isomerase | 4.87 | 2.13 | 1.35 | 5.09E-04 | 1.67E-01 | 3.73E-01 |
| | | A8JAS8 | FKB53 | Peptidyl-prolyl cis-trans isomerase | 1.01 | 0.56 | 0.69 | 9.70E-01 | 4.54E-02 | 1.83E-01 |
| | | A8J0I6 | FKB62 | Peptidyl-prolyl cis-trans isomerase FKBP-type | 0.72 | 0.84 | 0.84 | 2.96E-02 | 1.46E-01 | 6.95E-02 |
| | | A8HPN3 | FKB99 | Peptidyl-prolyl cis-trans isomerase FKBP-type | 1.07 | 1.09 | 1.43 | 2.48E-01 | 5.21E-01 | 4.57E-02 |
| | | A8I9E1 | TPR1 | Predicted chloroplast-targeted protein | 0.68 | 0.54 | 0.44 | 2.45E-01 | 7.79E-02 | 4.88E-02 |
| 5 | Intracellular protein trafficking | A8J729 | AP1B1 | Putative uncharacterized protein AP1B1 | 1.97 | 3.71 | 3.03 | 1.77E-01 | 5.78E-03 | 1.06E-03 |
| | | A8IMC5 | AP4E1 | Epsilon-adaptin | 0.76 | 1.70 | 1.68 | 2.32E-01 | 1.85E-02 | 2.22E-03 |
| | | A8IZV8 | AP1S1 | Sigma1-Adaptin | 1.53 | 1.70 | 1.51 | 1.73E-05 | 4.03E-02 | 6.54E-02 |
| | | A8HXA2 | AP1M1 | Mu1-Adaptin | 0.93 | 1.97 | 1.44 | 7.75E-01 | 3.14E-03 | 6.76E-02 |
| | | A8ILF4 | AP2A1 | Alpha-adaptin | 1.25 | 3.37 | 2.95 | 2.67E-01 | 8.19E-03 | 1.72E-03 |
| | | A8J7K1 | CLC1 | Clathrin light chain | 3.43 | 2.41 | 1.44 | 3.25E-02 | 4.35E-02 | 2.28E-01 |
| | | A8I4S9 | CHC1 | Clathrin heavy chain | 1.03 | 8.04 | 6.58 | 9.30E-01 | 2.29E-02 | 7.17E-03 |
| | | A8HRR9 | COPA1 | Alpha-COP | 1.42 | 5.23 | 4.70 | 3.56E-01 | 9.10E-03 | 3.75E-03 |
| | | A8JEP9 | COPB1 | Coatomer subunit beta | 0.66 | 4.04 | 2.90 | 2.26E-01 | 4.14E-02 | 5.71E-02 |
| | | A8JGS8 | COPB2 | Beta-cop | 2.31 | 4.05 | 2.41 | 1.14E-02 | 1.44E-03 | 1.87E-02 |
| | | A8JCA4 | CHLREDRAFT_195581 | RabGAP/TBC protein | 3.94 | 7.18 | 6.10 | 3.33E-02 | 3.28E-06 | 1.06E-04 |
| | | A8JC30 | SAR1 | Sar-type small GTPase | 1.11 | 1.03 | 0.80 | 2.60E-01 | 5.01E-01 | 1.10E-02 |

| | | | | | | | | | | |
|---|---|---|---|---|---|---|---|---|---|---|
| | | A8HZC8 | SEC23B | COP-II coat subunit | 0.93 | 2.06 | 1.23 | 1.67E-01 | 1.55E-01 | 1.55E-02 |
| | | A8I985 | SEC24A | COP-II coat subunit | 0.68 | 1.59 | 1.88 | 4.24E-02 | 6.24E-02 | 9.95E-02 |
| | | A8I1Y7 | SNAPA1 | Alpha-SNAP | 3.10 | 5.13 | 5.41 | 1.28E-02 | 8.95E-03 | 1.46E-03 |
| | | Q8S4W5 | SYP6 | Qc-SNARE protein Tlg1/Syntaxin 6-family | 2.79 | 3.31 | 2.84 | 7.32E-03 | 4.36E-04 | 1.96E-03 |
| | | A8IR64 | SYP5 | Qc-SNARE protein Syn8/Syntaxin8-family | 3.57 | 5.29 | 4.91 | 1.89E-02 | 4.18E-02 | 1.21E-02 |
| | | A8JBN4 | VCL1 | VPS16-like protein | 0.66 | 0.72 | 1.37 | 1.12E-01 | 4.01E-01 | 4.83E-02 |
| | | A8J4X1 | VPS26 | Subunit of retromer complex | 4.28 | 3.71 | 2.75 | 6.01E-03 | 1.76E-02 | 3.48E-03 |
| | | A8HQF0 | VPS35 | Subunit of retromer complex | 1.62 | 2.51 | 2.35 | 4.46E-03 | 3.29E-02 | 3.29E-02 |
| | | A8JCM8 | VPS45 | SM/Sec1-family protein | 2.30 | 2.25 | 2.10 | 4.03E-02 | 7.28E-02 | 2.18E-01 |
| | | A8I211 | CHLREDRAFT_161043 | Predicted protein | 1.84 | 2.17 | 1.99 | 7.90E-03 | 1.03E-02 | 9.29E-03 |
| | | A8I3S5 | CHLRE_02g108050v5 | Predicted protein | 2.46 | 3.82 | 4.44 | 2.51E-02 | 3.24E-02 | 1.81E-02 |
| | | A8IYJ6 | CGL38 | Predicted protein | 1.44 | 0.34 | 0.33 | 7.21E-02 | 1.84E-03 | 2.10E-03 |
| | | A8IAJ1 | VPS4 | AAA-ATPase of VPS4/SKD1 family | 0.90 | 0.94 | 0.49 | 3.58E-01 | 5.48E-01 | 3.00E-02 |
| | | A8HSQ2 | FAP66 | Flagellar associated protein | 0.66 | 2.44 | 1.86 | 2.39E-03 | 4.85E-02 | 2.44E-02 |
| | | A8J7M4 | CHLREDRAFT_150402 | Predicted protein (Fragment) | 0.74 | 1.40 | 1.39 | 1.45E-02 | 7.61E-03 | 2.04E-02 |
| | | A8IYH0 | CHLRE_12g549950v5 | Tetraspanning membrane protein SFT2-like protein | 0.14 | 0.48 | 0.53 | 1.86E-04 | 3.32E-02 | 7.95E-03 |
| | | A8I023 | TIC110 | 110 kDa translocon of chloroplast envelope inner membrane (Fragment) | 0.20 | 0.41 | 0.29 | 5.34E-07 | 6.72E-04 | 1.63E-03 |
| | | A8IJ46 | RNB2 | 3'-5' exoribonuclease II (Fragment) | 0.77 | 0.72 | 0.65 | 2.53E-01 | 8.61E-03 | 1.60E-01 |
| | | Q6Y682 | Rap38 | 38 kDa ribosome-associated protein | 1.05 | 0.70 | 0.38 | 7.94E-03 | 1.96E-02 | 9.83E-03 |
| | | A8IA39 | EFG1 | Chloroplast elongation factor G | 0.36 | 0.45 | 0.15 | 1.35E-03 | 6.65E-03 | 9.62E-05 |
| | | Q5S7Y5 | TIM | Chloroplast triosephosphate isomerase | 2.65 | 4.02 | 3.91 | 1.42E-01 | 8.98E-03 | 3.29E-02 |
| | | A8J680 | SECA2 | Chloroplast-associated SecA protein (Fragment) | 0.70 | 0.76 | 0.71 | 3.95E-02 | 1.76E-02 | 7.94E-02 |
| | | A8J6V5 | CHLREDRAFT_205591 | Diaminohydroxyphosphoribosylaminopyrimidine deaminase | 0.71 | 0.62 | 0.74 | 1.39E-01 | 2.62E-02 | 1.08E-02 |
| | | A8IZX1 | TEF24 | LrgB-like protein (Fragment) | 0.46 | 1.14 | 1.62 | 3.72E-02 | 3.86E-01 | 1.09E-01 |
| | | A8HWL8 | CHLREDRAFT_142189 | Predicted protein | 1.05 | 1.69 | 2.28 | 8.62E-01 | 4.25E-02 | 7.42E-02 |
| | | A8IB12 | CHLREDRAFT_126186 | Predicted protein (Fragment) | 0.91 | 0.83 | 0.54 | 3.45E-01 | 3.93E-02 | 1.89E-01 |
| | | A8J209 | CGL59 | Predicted protein | 0.36 | 0.41 | 0.49 | 3.62E-03 | 1.03E-02 | 1.29E-02 |
| | | A8J682 | SECA1 | Chloroplast-associated SecA protein | 0.38 | 0.69 | 0.78 | 6.39E-03 | 6.16E-02 | 9.33E-02 |
| 6 | TCA and ATP production | Q96550 | atpA | ATP synthase subunit alpha | 3.11 | 3.36 | 3.25 | 2.40E-02 | 1.01E-03 | 3.40E-02 |
| | | Q37304 | atpH | ATP synthase subunit c chloroplastic | 0.30 | 0.22 | 0.31 | 1.10E-04 | 7.18E-04 | 1.23E-05 |
| | | A8ID12 | ATP1B | Mitochondrial F1F0 ATP synthase alpha subunit (Fragment) | 0.77 | 0.75 | 0.41 | 2.81E-01 | 1.47E-01 | 2.94E-04 |
| | | A8HX15 | CHLREDRAFT_187139 | Sodium/potassium-transporting ATPase alpha subunit | 2.93 | 12.74 | 7.50 | 1.99E-04 | 2.51E-02 | 7.30E-03 |
| | | A8I164 | ATPvA1 | Vacuolar ATP synthase subunit A | 2.58 | 5.07 | 3.97 | 1.88E-02 | 7.24E-03 | 1.80E-02 |
| | | A8IA45 | ATPvB | Vacuolar ATP synthase subunit B | 4.48 | 3.30 | 2.45 | 1.46E-02 | 4.10E-02 | 2.93E-01 |
| | | A8IW47 | ATPvE | Vacuolar ATP synthase subunit E | 2.81 | 2.43 | 2.00 | 1.08E-02 | 6.37E-03 | 4.98E-02 |
| | | A8IDY0 | ATPvD1 | Vacuolar H+ ATPase V0 sector subunit D | 3.37 | 3.84 | 2.71 | 7.69E-03 | 7.00E-02 | 1.37E-01 |
| | | A8IST3 | ATPvA3 | Vacuolar proton ATPase subunit A | 1.27 | 3.97 | 3.96 | 4.52E-01 | 1.88E-03 | 3.35E-02 |
| | | A8J1K0 | ATPvA2 | Vacuolar proton translocating ATPase subunit A | 1.39 | 4.26 | 3.48 | 2.66E-01 | 1.82E-03 | 1.65E-03 |

| | | | | | | | | | | |
|---|---|---|---|---|---|---|---|---|---|---|
| | | A8J588 | ATPvL2 | Vacuolar proton-ATPase subunit c" proteolipid | 0.67 | 1.30 | 1.74 | 4.94E-02 | 2.88E-01 | 8.54E-02 |
| | | A8HZ87 | ATPvF | V-type proton ATPase subunit F | 1.81 | 1.88 | 1.20 | 8.71E-03 | 1.05E-04 | 5.45E-01 |
| | | Q8RVB8 | ATP6 | ATP synthase F1F0 subunit 6 | 1.51 | 0.89 | 1.05 | 2.24E-02 | 2.16E-01 | 6.20E-01 |
| | | o63075 | ATPC | ATP synthase gamma chain | 7.84 | | 14.71 | 6.64E-02 | 1.13E-01 | 3.25E-02 |
| | | o63075 | atpI | ATP synthase subunit a chloroplastic | 0.49 | 0.37 | 0.79 | 6.64E-02 | 1.13E-01 | 3.25E-02 |
| | | Q8HTL5 | atpF | ATP synthase subunit b chloroplastic | 0.61 | 0.80 | 0.99 | 3.06E-06 | 4.11E-01 | 7.23E-01 |
| | | A8ICT4 | ATP15 | F1F0 ATP synthase epsilon subunit | 2.66 | 0.93 | 0.44 | 2.44E-01 | 8.70E-01 | 4.27E-04 |
| | | A8J3U9 | ATP5 | Mitochondrial ATP synthase subunit 5 OSCP subunit | 3.92 | 1.84 | 2.66 | 5.64E-03 | 7.44E-03 | 1.54E-02 |
| | | A8J9X1 | ATP4 | Mitochondrial F1F0 ATP synthase delta subunit | 0.88 | 0.26 | 0.25 | 5.45E-01 | 1.24E-04 | 1.38E-05 |
| | | A8IVG0 | OGD1 | 2-oxoglutarate dehydrogenase E1 subunit | 1.32 | 3.89 | 3.32 | 5.12E-01 | 4.02E-02 | 2.90E-02 |
| | | A8HMQ1 | ACH1 | Aconitate hydratase | 1.47 | 2.98 | 2.00 | 2.43E-01 | 1.37E-02 | 6.05E-02 |
| | | A8JHC9 | CIS1 | Citrate synthase | 12.65 | 14.93 | 15.41 | 3.20E-03 | 8.86E-03 | 2.76E-03 |
| | | A8J2S0 | CIS2 | Citrate synthase | 0.30 | 0.27 | 0.19 | 3.32E-05 | 6.08E-04 | 9.34E-05 |
| | | A8HX04 | SDH2 | Iron-sulfur subunit of mitochondrial succinate dehydrogenase | 11.93 | 12.17 | 9.00 | 2.54E-02 | 4.14E-03 | 4.07E-03 |
| | | A8J9S7 | IDH3 | Isocitrate dehydrogenase [NADP] | 4.22 | 5.58 | 2.85 | 8.92E-02 | 4.00E-03 | 3.96E-02 |
| | | A8J0R7 | IDH2 | Isocitrate dehydrogenase NAD-dependent | 1.74 | 1.00 | 0.70 | 3.60E-02 | 9.75E-01 | 2.15E-01 |
| | | A8ICG9 | MDH2 | Malate dehydrogenase | 0.15 | 0.75 | 0.86 | 1.65E-06 | 2.22E-01 | 2.80E-01 |
| | | A8J0W9 | MDH3 | Malate dehydrogenase | 0.98 | 1.87 | 2.17 | 8.80E-01 | 1.34E-02 | 4.93E-03 |
| | | A8JHU0 | MDH4 | Malate dehydrogenase | 9.33 | 11.97 | 9.63 | 4.38E-02 | 8.14E-03 | 9.98E-03 |
| | | Q6X898 | MAS1 | Malate synthase | 0.56 | 0.18 | 0.20 | 1.14E-03 | 2.56E-04 | 7.34E-04 |
| | | Q9FNS5 | NADP-mdh | NADP-Malate dehydrogenase | 0.26 | 0.25 | 0.32 | 4.42E-06 | 9.68E-05 | 3.02E-04 |
| | | A8HP06 | SDH1 | Succinate dehydrogenase subunit A | 1.52 | 1.74 | 1.12 | 2.13E-04 | 1.63E-03 | 7.47E-01 |
| | | A8HPU1 | SDH4 | Succinate dehydrogenase subunit D | 0.78 | 0.57 | 0.42 | 1.10E-01 | 1.99E-03 | 9.67E-06 |
| 7 | Amino acid and sulfur metabolism | A8J6Q7 | SHKA1 | 3-deoxy-D-arabino-heptulosonate 7-phosphate synthetase | 0.26 | 0.44 | 0.45 | 1.51E-04 | 1.26E-05 | 1.73E-02 |
| | | A8JH48 | SHKG1 | 3-phosphoshikimate 1-carboxyvinyltransferase | 0.13 | 0.25 | 0.39 | 3.77E-06 | 4.41E-04 | 1.04E-03 |
| | | A8J2Z6 | SHKH1 | Chorismate synthase | 0.65 | 0.45 | 0.48 | 1.75E-01 | 1.38E-02 | 7.97E-04 |
| | | A8J434 | OASTL1 | Cysteine synthase | 0.60 | 0.56 | 0.58 | 8.62E-03 | 5.61E-03 | 3.39E-02 |
| | | A8IEE5 | OASTL3 | Cysteine synthase | 2.69 | 3.01 | 3.11 | 1.34E-01 | 9.87E-02 | 5.00E-02 |
| | | A8ISA9 | OASTL4 | Cysteine synthase | 19.91 | 19.97 | 20.34 | 3.46E-04 | 4.81E-03 | 1.22E-03 |
| | | A8JG03 | LEU1L | Isopropylmalate dehydratase large subunit | 1.62 | 3.80 | 7.11 | 1.53E-01 | 1.04E-01 | 8.03E-05 |
| | | A8IFZ9 | MAA7 | Tryptophan synthase beta subunit | 0.44 | 0.56 | 0.36 | 2.95E-05 | 4.42E-02 | 1.22E-04 |
| | | A8I9R1 | THD1 | Threonine deaminase | 0.28 | 0.42 | 0.46 | 5.15E-04 | 3.69E-03 | 7.39E-03 |
| | | A8HPI1 | AGK1 | Acetylglutamate kinase-like protein | 7.02 | 5.19 | 5.30 | 8.98E-03 | 6.80E-02 | 2.32E-03 |
| | | A8IRD4 | PROB2 | Glutamate 5-kinase | 0.45 | 0.55 | 0.39 | 2.53E-02 | 1.72E-02 | 1.26E-02 |
| | | A8IRD5 | PROB1 | Glutamate 5-kinase | 0.70 | 0.76 | 0.87 | 1.63E-02 | 1.76E-04 | 6.18E-01 |
| | | A8J2W0 | GSD1 | Glutamic-gamma-semialdehyde dehydrogenase | 0.33 | 1.25 | 1.21 | 9.03E-05 | 9.32E-02 | 1.59E-02 |
| | | A8HTR6 | CHLRE_13g579800v5 | Predicted amino acid kinase (Fragment) | 0.37 | 0.50 | 0.33 | 2.03E-05 | 4.03E-02 | 1.69E-02 |
| | | A8IBN0 | PRT1 | Anthranilate phosphoribosyltransferase | 0.44 | 0.31 | 0.39 | 9.34E-03 | 6.67E-03 | 1.99E-02 |

| | | | | | | | | | | |
|---|---|---|---|---|---|---|---|---|---|---|
| | | A8IMY5 | ANS1 | Anthranilate synthase alpha subunit (Fragment) | 0.74 | 0.48 | 0.63 | 1.11E-01 | 5.91E-03 | 5.49E-02 |
| | | A8HP28 | ASB1 | Anthranilate synthase beta subunit | 1.17 | 0.72 | 1.21 | 1.81E-01 | 1.32E-02 | 2.55E-01 |
| | | A8J599 | TSA1 | Tryptophan synthetase alpha subunit | 0.57 | 0.55 | 0.47 | 1.90E-02 | 7.27E-02 | 2.38E-02 |
| | | A8HT60 | IGS1 | Indole-3-glycerol-phosphate synthase | 0.76 | 0.63 | 0.53 | 2.68E-03 | 8.92E-05 | 2.06E-02 |
| | | A8J3Q6 | APK1 | Adenylyl-sulfate kinase | 2.75 | 2.17 | 2.09 | 4.66E-02 | 1.61E-01 | 3.67E-02 |
| | | A8IXF1 | ATS1 | ATP-sulfurylase | 1.04 | 4.89 | 4.87 | 9.13E-01 | 1.16E-01 | 4.55E-03 |
| | | A8I3V3 | ATS2 | ATP-sulfurylase | 8.52 | 17.05 | 18.20 | 2.18E-03 | 1.36E-02 | 1.07E-02 |
| | | A8HYU5 | METM | S-adenosylmethionine synthase | 0.68 | 0.41 | 0.13 | 9.51E-02 | 1.27E-02 | 3.02E-06 |
| | | A8IXE0 | SAH1 | Adenosyl homocysteinase | 7.50 | 5.86 | 1.39 | 2.25E-03 | 1.26E-04 | 5.19E-01 |
| | | A8HRS5 | CHLREDRAFT_127560 | $N^{10}$-formyltetrahydrofolate synthetase | 1.22 | 0.60 | 0.29 | 4.19E-01 | 1.14E-03 | 1.05E-03 |
| | | A8IQS8 | CHLREDRAFT_206121 | Methylenetetrahydrofolate dehydrogenase/ [illegible] | 0.79 | 0.63 | 0.56 | 5.94E-03 | 6.28E-02 | 4.36E-02 |
| | | A8IAY6 | CHLREDRAFT_111330 | Methylenetetrahydrofolate reductase | 0.95 | 0.54 | 0.22 | 5.54E-01 | 4.96E-02 | 1.52E-04 |
| | | A8J4Z8 | CAH3 | Carbonic anhydrase 3 | 0.76 | 0.26 | 0.23 | 2.84E-01 | 1.09E-04 | 4.77E-04 |
| | | A8IT01 | CAH1 | Carbonic anhydrase | 0.27 | 0.23 | 0.28 | 1.80E-03 | 1.08E-04 | 5.24E-05 |
| | | A8IAK9 | PYR2 | Aspartate carbamoyltransferase | 0.56 | 0.30 | 0.36 | 1.39E-01 | 2.58E-03 | 1.37E-04 |
| | | A8IMN5 | CMPS1 | Carbamoyl phosphate synthase small subunit | 1.73 | 1.82 | 1.52 | 3.17E-01 | 5.80E-06 | 1.15E-01 |
| | | A8JBN5 | PYR4 | Dihydropryrimidine dehydrogenase | 1.86 | 2.62 | 3.10 | 2.65E-02 | 2.81E-02 | 1.80E-02 |
| | | A8JJR9 | FAK2 | Flagellar adenylate kinase (Fragment) | 1.51 | 10.07 | 7.44 | 4.72E-01 | 2.89E-02 | 9.32E-02 |
| | | A8HN92 | PYR5 | Uridine 5'-monophosphate synthase | 1.23 | 2.15 | 1.50 | 3.78E-02 | 2.64E-02 | 2.08E-01 |
| 8 | Response to cytokinin | A8J3F8 | CHLRE_16g672750v5 | Predicted protein | 0.58 | 0.25 | 0.23 | 4.94E-04 | 1.92E-06 | 7.76E-06 |
| | | A8JF10 | LHCA3 | Light-harvesting chlorophyll-a/b protein of photosystem I type III | 0.72 | 0.61 | 0.59 | 3.46E-01 | 4.66E-02 | 1.45E-02 |
| | | Q70DX8 | S1 | Plastid ribosomal protein S1 | 0.49 | 0.19 | 0.13 | 6.83E-03 | 4.02E-05 | 1.11E-04 |
| | | A8IRG9 | CPLD28 | Predicted protein | 0.91 | 0.74 | 0.91 | 1.03E-01 | 3.28E-02 | 3.33E-01 |
| | | A8HWZ6 | PRPL13 | Plastid ribosomal protein L13 | 0.32 | 0.08 | 0.08 | 3.65E-04 | 1.59E-08 | 4.22E-07 |
| | | A8IU62 | PIN3 | Peptidyl-prolyl cis-trans isomerase parvulin-type | 0.63 | 0.58 | 0.57 | 2.36E-03 | 2.44E-02 | 4.45E-02 |
| | | A8HXG5 | PSBR | 10 kDa photosystem II polypeptide | 0.78 | 0.42 | 0.27 | 2.42E-02 | 9.25E-03 | 1.62E-02 |
| | | A8HVJ9 | CHLREDRAFT_112806 | Photosystem II stability/assembly factor HCF136 | 0.24 | 0.10 | 0.13 | 1.47E-04 | 4.77E-06 | 2.84E-05 |
| | | A8J503 | PRPL6 | Plastid ribosomal protein L6 | 0.94 | 0.16 | 0.15 | 8.44E-01 | 3.37E-05 | 6.67E-05 |
| | | Q8HTL1 | rpl5 | 50S ribosomal protein L5 chloroplastic | 2.79 | 0.68 | 0.37 | 1.09E-02 | 6.66E-02 | 1.61E-03 |
| | | A8JAL6 | PRPL15 | Plastid ribosomal protein L15 | 0.24 | 0.04 | 0.05 | 3.39E-04 | 2.86E-07 | 2.65E-07 |
| | | A8J785 | ATPG | ATP synthase subunit b' chloroplastic | 0.90 | 1.00 | 1.21 | 2.95E-02 | 9.98E-01 | 2.05E-03 |
| | | A8JC21 | UROD1 | Uroporphyrinogen decarboxylase | 0.41 | 0.21 | 0.24 | 3.97E-03 | 3.99E-04 | 7.65E-03 |
| | | A8IYP4 | PRK1 | Phosphoribulokinase | 1.04 | 0.12 | 0.11 | 8.34E-01 | 2.66E-06 | 1.77E-09 |
| | | A8J8U1 | RPN10 | 26S proteasome regulatory subunit | 0.84 | 0.65 | 0.79 | 2.76E-01 | 1.31E-03 | 4.13E-03 |
| | | A8I8X2 | DEG1A | DegP-type protease | 1.40 | 1.14 | 0.37 | 3.26E-01 | 8.02E-01 | 3.93E-02 |
| | | A8IKQ0 | FBP1 | Fructose-1,6-bisphosphatase | 0.66 | 0.28 | 0.07 | 1.41E-05 | 1.39E-04 | 5.99E-08 |
| | | A8INR7 | PRPL27 | Plastid ribosomal protein L27 | 0.26 | 0.14 | 0.15 | 8.50E-04 | 8.24E-05 | 9.15E-05 |

**Table S3.** List of differentially expressed proteins in *hpm91* delineated from Dataset S6.

| Ranking | GOPB | Uniprot accession | Gene ID | Protein name | Ratio (Mut/Mu0) | | | *p*-value | | |
|---|---|---|---|---|---|---|---|---|---|---|
| | | | | | 24 h | 72 h | 120 h | 24 h | 72 h | 120 h |
| 1 | Translation | A8J576 | RPS27-A | 40S ribosomal protein S27 | 0.45 | 0.17 | 0.12 | 2.93E-02 | 7.18E-03 | 5.28E-03 |
| | | A8J0V6 | RPS27-B | 40S ribosomal protein S27 | 2.19 | 2.93 | 2.10 | 6.83E-02 | 4.64E-02 | 8.00E-02 |
| | | A8HVQ1 | RPS8 | 40S ribosomal protein S8 | 0.34 | 0.16 | 0.12 | 1.55E-03 | 2.76E-04 | 3.92E-04 |
| | | A8IB25 | CHLREDRAFT_126059 | 40S ribosomal protein SA | 0.88 | 0.68 | 0.54 | 2.84E-01 | 3.35E-02 | 4.21E-02 |
| | | A8IUV7 | RPL13 | 60S ribosomal protein L13 | 1.55 | 0.52 | 0.36 | 1.81E-01 | 9.02E-03 | 2.48E-03 |
| | | A8IKZ2 | RPL18 | 60S ribosomal protein L18 | 1.18 | 0.57 | 0.32 | 5.55E-01 | 3.45E-02 | 7.44E-03 |
| | | A8IQC1 | RPL27 | 60S ribosomal protein L27 | 6.40 | 3.47 | 3.09 | 2.04E-03 | 5.01E-02 | 5.34E-02 |
| | | Q8GUQ9 | RPL38 | 60S ribosomal protein L38 | 3.92 | 1.77 | 1.91 | 1.10E-02 | 6.47E-04 | 1.43E-01 |
| | | A8JCA8 | EFG5 | Elongation factor EF-Tu-like protein | 0.96 | 2.46 | 1.88 | 5.97E-01 | 5.96E-03 | 4.29E-02 |
| | | A8JB67 | NHP2 | Nucleolar protein small subunit of H/ACA snoRNPs | 0.76 | 0.57 | 0.54 | 8.38E-02 | 3.16E-02 | 6.80E-02 |
| | | A8J9X5 | CHLREDRAFT_18599 | Predicted protein | 0.85 | 0.58 | 0.58 | 3.29E-01 | 4.36E-02 | 1.43E-01 |
| | | A8JHJ5 | CHLRE_15g641200v5 | Predicted protein | 0.14 | 0.14 | 0.12 | 1.87E-02 | 1.95E-02 | 1.67E-02 |
| | | A8I9M5 | CHLREDRAFT_141578 | Predicted protein | 0.16 | 0.10 | 0.09 | 7.09E-04 | 1.28E-04 | 1.26E-04 |
| | | A8I4T2 | RPL10a | Ribosomal protein | 10.16 | 9.34 | 7.90 | 7.41E-03 | 9.96E-03 | 1.64E-02 |
| | | A8IZK3 | RPL10 | Ribosomal protein L10 | 0.69 | 0.41 | 0.21 | 2.71E-01 | 6.23E-02 | 2.27E-02 |
| | | A8HQ81 | RPL11 | Ribosomal protein L11 | 4.01 | 1.94 | 1.52 | 4.15E-04 | 2.21E-03 | 4.46E-03 |
| | | A8IA18 | RPL19 | Ribosomal protein L19 | 0.43 | 0.15 | 0.11 | 2.37E-02 | 6.66E-03 | 5.78E-03 |
| | | A8J951 | RPL21 | Ribosomal protein L21 | 3.25 | 1.56 | 1.31 | 2.87E-02 | 2.27E-01 | 5.72E-01 |
| | | A8J239 | RPL23a | Ribosomal protein L23a | 1.00 | 0.38 | 0.34 | 9.93E-01 | 1.74E-02 | 3.25E-02 |
| | | A8I2T0 | RPL27a | Ribosomal protein L27a | 0.56 | 0.29 | 0.17 | 1.13E-01 | 2.20E-02 | 1.34E-02 |
| | | A8ID84 | RPL3 | Ribosomal protein L3 | 0.35 | 0.28 | 0.13 | 2.44E-05 | 1.00E-05 | 3.36E-06 |
| | | A8ILG8 | RPL31 | Ribosomal protein L31 | 0.80 | 0.25 | 0.23 | 6.12E-01 | 5.20E-03 | 3.38E-03 |
| | | A8J2G4 | RPL32 | Ribosomal protein L32 | 0.85 | 0.45 | 0.25 | 6.17E-01 | 5.13E-02 | 6.51E-03 |
| | | A8J8P4 | RPL34 | Ribosomal protein L34 | 0.42 | 0.35 | 0.28 | 1.62E-02 | 3.43E-03 | 3.90E-03 |
| | | A8HNX3 | RPL35 | Ribosomal protein L35 | 10.24 | 4.02 | 3.79 | 7.92E-03 | 9.88E-02 | 3.01E-01 |
| | | A8J0I0 | RPL4 | Ribosomal protein L4 | 0.55 | 0.33 | 0.20 | 8.05E-02 | 2.13E-02 | 1.34E-02 |
| | | A8HP55 | RPL5 | Ribosomal protein L5 | 2.31 | 1.65 | 1.36 | 2.93E-02 | 8.88E-02 | 3.81E-01 |
| | | A8J567 | RPL7a | Ribosomal protein L7a | 4.76 | 3.33 | 2.29 | 4.52E-02 | 7.26E-02 | 2.15E-01 |
| | | A8JHC3 | RPS11 | Ribosomal protein S11 | 0.23 | 0.18 | 0.09 | 1.11E-02 | 8.20E-03 | 5.55E-03 |
| | | A8JE07 | RPS15a | Ribosomal protein S15a | 2.94 | 1.35 | 0.96 | 1.46E-02 | 5.48E-01 | 6.01E-01 |
| | | A8JGK1 | RPS17 | Ribosomal protein S17 | 8.26 | 4.47 | 3.14 | 2.23E-02 | 4.37E-02 | 1.08E-01 |
| | | A8HVP2 | RPS18 | Ribosomal protein S18 | 7.16 | 5.10 | 4.23 | 1.90E-02 | 2.74E-02 | 1.05E-01 |
| | | A8I403 | RPS19 | Ribosomal protein S19 | 9.38 | 4.81 | 3.55 | 2.78E-03 | 4.07E-03 | 5.75E-02 |

| | | | | | | | | |
|---|---|---|---|---|---|---|---|---|
| A8HME4 | RPS2 | Ribosomal protein S2 | 1.08 | 0.59 | 0.43 | 7.55E-01 | 2.69E-02 | 1.13E-02 |
| A8J8M9 | RPS20 | Ribosomal protein S20 | 0.94 | 0.35 | 0.32 | 7.13E-01 | 2.02E-03 | 8.37E-03 |
| A8IS22 | RPS26 | Ribosomal protein S26 | 0.68 | 0.35 | 0.25 | 3.12E-01 | 4.12E-02 | 2.10E-02 |
| A8HVK4 | RPS27a | Ribosomal protein S27a | 0.23 | 0.16 | 0.12 | 2.02E-04 | 2.78E-04 | 1.06E-04 |
| A8I4P5 | RPS3 | Ribosomal protein S3 | 0.92 | 0.42 | 0.36 | 8.37E-01 | 6.11E-02 | 4.83E-02 |
| A8JF66 | RPS30 | Ribosomal protein S30 | 0.70 | 0.37 | 0.27 | 1.68E-01 | 1.78E-02 | 8.68E-03 |
| A8IMP6 | RPS4 | Ribosomal protein S4 | 0.90 | 0.41 | 0.27 | 8.48E-01 | 4.79E-02 | 2.11E-02 |
| A8J2I5 | RPS5 | Ribosomal protein S5 | 5.85 | 5.29 | 4.81 | 5.04E-02 | 1.87E-02 | 1.45E-01 |
| A8JGF8 | RPS9 | Ribosomal protein S9 component of cytosolic 80S ribosome and 40S small subunit | 5.46 | 2.10 | 1.72 | 3.72E-02 | 9.80E-02 | 2.94E-01 |
| A8J8J8 | CHLREDRAFT_120661 | GTPase Der (Fragment) | 0.47 | 0.48 | 0.32 | 3.12E-02 | 3.46E-02 | 1.62E-02 |
| A8I2G3 | CHLREDRAFT_112251 | Predicted protein (Fragment) | 0.53 | 0.53 | 0.48 | 2.58E-03 | 2.01E-02 | 3.95E-03 |
| A8JHU2 | RPL36 | 60S ribosomal protein L36 | 7.38 | 2.61 | 2.37 | 2.91E-02 | 1.14E-01 | 1.88E-01 |
| A8I982 | RPL15 | Ribosomal protein L15 | 0.84 | 0.53 | 0.29 | 7.78E-01 | 1.28E-01 | 3.72E-02 |
| A8JI94 | RPL22 | Ribosomal protein L22 | 20.81 | 11.42 | 10.69 | 5.90E-03 | 1.56E-02 | 1.23E-02 |
| A8HMG7 | RPL26 | Ribosomal protein L26 | 1.45 | 0.58 | 0.42 | 1.27E-01 | 4.75E-02 | 1.87E-02 |
| A8I0Y2 | RPL35a | Ribosomal protein L35a | 1.14 | 0.42 | 0.32 | 6.68E-01 | 5.45E-02 | 2.44E-02 |
| A8IVE2 | RPL7 | Ribosomal protein L7 | 5.18 | 3.42 | 2.59 | 1.37E-03 | 4.42E-03 | 3.59E-02 |
| A8IVK1 | RPL8 | Ribosomal protein L8 | 0.32 | 0.20 | 0.20 | 1.66E-02 | 9.03E-03 | 1.11E-02 |
| Q7YKX3 | rps11 | 30S ribosomal protein S11 chloroplastic | 0.45 | 0.21 | 0.25 | 4.65E-02 | 1.23E-02 | 1.45E-02 |
| P59776 | rps19 | 30S ribosomal protein S19 chloroplastic | 0.27 | 0.07 | 0.07 | 3.50E-03 | 2.63E-04 | 2.61E-04 |
| O47027 | rps2-1 | 30S ribosomal protein S2 chloroplastic | 0.26 | 0.18 | 0.18 | 1.01E-02 | 8.05E-03 | 7.46E-03 |
| Q08365 | rps3 | 30S ribosomal protein S3 chloroplastic | 0.23 | 0.08 | 0.08 | 2.07E-02 | 1.01E-02 | 9.66E-03 |
| P48270 | rps4 | 30S ribosomal protein S4 chloroplastic | 0.66 | 0.26 | 0.27 | 1.09E-02 | 1.74E-03 | 3.97E-04 |
| P48267 | rps7 | 30S ribosomal protein S7 chloroplastic | 2.65 | 1.47 | 1.71 | 2.50E-03 | 1.12E-01 | 2.17E-02 |
| O20029 | rps9 | 30S ribosomal protein S9 chloroplastic | 0.23 | 0.09 | 0.10 | 1.92E-03 | 9.68E-04 | 9.97E-04 |
| Q8HTL2 | rpl2 | 50S ribosomal protein L2 chloroplastic | 0.22 | 0.06 | 0.05 | 1.86E-03 | 8.92E-04 | 8.51E-04 |
| P26565 | rpl20 | 50S ribosomal protein L20 chloroplastic | 1.28 | 0.47 | 0.26 | 3.53E-01 | 2.83E-02 | 1.18E-02 |
| Q8HTL3 | rpl23 | 50S ribosomal protein L23 chloroplastic | 0.64 | 0.16 | 0.13 | 1.51E-01 | 1.56E-02 | 1.59E-02 |
| A8JEP1 | PRPL35 | 50S ribosomal protein L35 | 0.91 | 0.35 | 0.28 | 8.87E-01 | 3.70E-02 | 7.66E-02 |
| Q8HTL1 | rpl5 | 50S ribosomal protein L5 chloroplastic | 1.33 | 0.64 | 0.52 | 2.64E-01 | 6.14E-03 | 5.84E-04 |
| A8I8Z4 | PRPL1 | Plastid ribosomal protein L1 | 0.41 | 0.08 | 0.07 | 5.36E-03 | 1.22E-03 | 1.04E-03 |
| A8ICE4 | PRPL11 | Plastid ribosomal protein L11 | 0.86 | 0.40 | 0.36 | 7.23E-01 | 6.94E-03 | 4.67E-03 |
| A8HWZ6 | PRPL13 | Plastid ribosomal protein L13 | 0.43 | 0.12 | 0.15 | 3.68E-02 | 1.08E-02 | 1.08E-02 |
| A8JAL6 | PRPL15 | Plastid ribosomal protein L15 | 0.33 | 0.05 | 0.05 | 2.04E-02 | 5.06E-03 | 5.00E-03 |
| A8I3M4 | PRPL17 | Plastid ribosomal protein L17 | 1.41 | 0.57 | 0.52 | 2.03E-01 | 5.56E-02 | 3.12E-02 |
| A8HNJ8 | PRPL18 | Plastid ribosomal protein L18 | 0.82 | 0.17 | 0.12 | 4.36E-01 | 1.99E-02 | 1.55E-02 |
| A8IW44 | PRPL19 | Plastid ribosomal protein L19 | 0.33 | 0.11 | 0.13 | 2.24E-02 | 6.37E-03 | 6.79E-03 |
| A8J9D9 | PRPL24 | Plastid ribosomal protein L24 | 1.06 | 0.21 | 0.28 | 8.00E-01 | 2.37E-04 | 7.63E-03 |

| | | | | | | | | | | |
|---|---|---|---|---|---|---|---|---|---|---|
| | | A8INR7 | PRPL27 | Plastid ribosomal protein L27 | 0.19 | 0.05 | 0.04 | 2.00E-03 | 8.47E-04 | 7.99E-04 |
| | | A8HWS8 | PRPL28 | Plastid ribosomal protein L28 | 0.23 | 0.06 | 0.04 | 5.36E-03 | 2.14E-03 | 1.96E-03 |
| | | A8JE35 | PRPL3 | Plastid ribosomal protein L3 | 0.28 | 0.07 | 0.05 | 1.78E-03 | 5.59E-04 | 4.50E-04 |
| | | A8IUC3 | PRPL32 | Plastid ribosomal protein L32 | 2.07 | 0.40 | 0.38 | 1.06E-01 | 1.19E-02 | 1.11E-02 |
| | | A8I1D3 | PRPL33 | Plastid ribosomal protein L33 | 1.24 | 0.55 | 0.39 | 3.48E-01 | 2.82E-02 | 2.34E-02 |
| | | A8J503 | PRPL6 | Plastid ribosomal protein L6 | 0.79 | 0.20 | 0.15 | 1.58E-01 | 3.33E-03 | 2.03E-03 |
| | | A8IYS1 | PRPL9 | Plastid ribosomal protein L9 | 0.80 | 0.23 | 0.18 | 5.23E-01 | 4.80E-02 | 4.01E-02 |
| | | A8JGS2 | PRPS17 | Plastid ribosomal protein S17 | 0.24 | 0.08 | 0.08 | 1.41E-03 | 9.29E-05 | 9.46E-05 |
| | | A8JDN4 | PRPS20 | Plastid ribosomal protein S20 | 2.80 | 1.28 | 1.39 | 9.34E-04 | 2.28E-01 | 2.27E-01 |
| | | A8J8M5 | PRPS5 | Plastid ribosomal protein S5 | 0.16 | 0.04 | 0.06 | 1.85E-04 | 8.32E-05 | 8.88E-05 |
| | | A8IMN3 | PSRP-6 | Plastid-specific ribosomal protein 6 | 0.11 | 0.04 | 0.04 | 1.77E-04 | 6.56E-05 | 7.10E-05 |
| | | A8IA39 | EFG1 | Chloroplast elongation factor G | 0.42 | 0.48 | 0.52 | 1.49E-02 | 2.51E-03 | 4.08E-02 |
| | | A8IPJ0 | MITC11 | Mitochondrial carrier protein | 1.74 | 1.25 | 1.28 | 3.06E-02 | 4.37E-01 | 5.06E-01 |
| | | A8IZL2 | MRPL21 | Mitochondrial ribosomal protein L21 | 1.00 | 0.80 | 1.02 | 9.29E-01 | 1.25E-02 | 7.37E-01 |
| | | A8HXM1 | MRPL29 | Mitochondrial ribosomal protein L29 | 11.63 | 4.41 | 3.79 | 3.95E-03 | 2.66E-02 | 1.72E-01 |
| | | A8JFK9 | MRPL7/L12 | Mitochondrial ribosomal protein L7/L12 | 2.95 | 3.97 | 3.80 | 5.64E-02 | 1.93E-02 | 3.47E-02 |
| | | A8HXY9 | UCP2 | Uncoupling protein | 3.24 | 3.56 | 2.82 | 1.54E-02 | 3.15E-03 | 7.65E-02 |
| | | A8JCF4 | MRPS17 | Mitochondrial ribosomal protein S17 | 0.56 | 0.27 | 0.59 | 1.29E-02 | 3.47E-03 | 9.44E-03 |
| | | A8J2A5 | MSCP1 | Mitochondrial substrate carrier protein | 0.91 | 0.34 | 0.41 | 6.79E-01 | 4.78E-03 | 1.05E-02 |
| 2 | Protein folding | A8J524 | CCT2 | T-complex protein 1 beta subunit | 1.72 | 2.10 | 1.52 | 1.85E-02 | 7.82E-03 | 3.03E-02 |
| | | A8II42 | CCT3 | T-complex protein 1 gamma subunit | 1.05 | 2.02 | 1.32 | 9.96E-01 | 1.88E-02 | 3.01E-01 |
| | | A8JE04 | CCT4 | T-complex protein 1 subunit delta | 1.49 | 3.08 | 1.98 | 2.59E-01 | 8.85E-03 | 2.13E-02 |
| | | A8J7J2 | CCT5 | T-complex protein epsilon subunit | 1.34 | 3.71 | 3.17 | 4.79E-01 | 2.42E-02 | 1.48E-01 |
| | | A8J014 | CCT6 | T-complex protein zeta subunit | 3.00 | 3.29 | 2.50 | 3.38E-02 | 1.77E-03 | 2.35E-02 |
| | | A8HQ74 | CCT7 | T-complex protein eta subunit | 0.89 | 2.87 | 2.40 | 4.60E-01 | 9.45E-03 | 7.80E-02 |
| | | A8IUG8 | CDSP32 | Plastidic thioredoxin-like protein | 1.02 | 0.26 | 0.42 | 9.20E-01 | 9.08E-03 | 2.25E-02 |
| | | A8HMC0 | CRT2 | Calreticulin 2 calcium-binding protein | 1.46 | 0.54 | 0.20 | 2.61E-01 | 6.07E-02 | 1.26E-02 |
| | | A8IV02 | CYN1a\|CYN1b | Peptidyl-prolyl cis-transisomerase cyclophilin type | 0.86 | 0.82 | 0.65 | 4.09E-01 | 1.33E-01 | 1.60E-02 |
| | | A8JD64 | CYN19-2 | Peptidyl-prolyl cis-trans isomerase | 0.70 | 0.60 | 0.45 | 2.07E-01 | 1.16E-01 | 4.91E-02 |
| | | A8J282 | CYN20-1 | Peptidyl-prolyl cis-trans isomerase | 4.78 | 7.32 | 2.43 | 1.27E-03 | 4.11E-03 | 2.37E-01 |
| | | A8JDL5 | CYN20-3 | Peptidyl-prolyl cis-trans isomerase | 0.79 | 0.44 | 0.31 | 5.31E-01 | 7.53E-02 | 3.44E-02 |
| | | A8ID98 | CYN20-5 | Peptidyl-prolyl cis-trans isomerase cyclophilin-type | 0.70 | 0.50 | 0.58 | 1.64E-01 | 6.48E-03 | 2.10E-02 |
| | | A8IE53 | CYN26 | Peptidyl-prolyl cis-trans isomerase cyclophilin-type | 1.07 | 0.29 | 0.44 | 8.49E-01 | 1.76E-02 | 3.66E-02 |
| | | A8JHN8 | CYN28 | Peptidyl-prolyl cis-trans isomerase cyclophilin-type | 1.03 | 0.32 | 0.57 | 6.74E-01 | 2.96E-02 | 1.16E-01 |
| | | A8HN48 | CYN40 | Peptidyl-prolyl cis-trans isomerase cyclophilin-type | 1.88 | 2.14 | 1.68 | 2.78E-02 | 1.07E-01 | 1.93E-01 |
| | | A8IQC5 | DNJ1 | DnaJ-like protein | 0.47 | 2.01 | 1.34 | 7.12E-02 | 2.98E-02 | 2.79E-01 |
| | | A8I6Y0 | ERJ1 | ER DnaJ-like protein 1 | 0.80 | 0.69 | 0.76 | 3.08E-01 | 3.86E-02 | 3.77E-01 |
| | | A8J3L6 | FKB16-2a\|FKB16-[illegible] | Peptidyl-prolyl cis-trans isomerase | 0.85 | 0.43 | 0.20 | 4.01E-01 | 6.75E-03 | 1.03E-02 |

| | | | | | | | | | | |
|---|---|---|---|---|---|---|---|---|---|---|
| | | A8J3C1 | FKB16-7a\|FKB16-7b | Peptidyl-prolyl cis-trans isomerase | 0.66 | 0.25 | 0.25 | 1.93E-01 | 1.58E-03 | 1.32E-03 |
| | | Q944P3 | HSP33 | Heat shock protein 33 | 0.54 | 0.37 | 0.51 | 2.24E-02 | 7.48E-03 | 1.95E-02 |
| | | A8J1U1 | HSP90A | Heat shock protein 90A | 3.89 | 5.74 | 3.69 | 5.61E-02 | 2.12E-02 | 4.93E-02 |
| | | A8I7T1 | HSP90B | Heat shock protein 90B | 5.17 | 5.76 | 5.33 | 5.98E-03 | 1.24E-03 | 2.06E-03 |
| | | A8JES1 | MGE1 | GrpE protein homolog | 5.54 | 2.70 | 3.45 | 8.02E-04 | 2.90E-03 | 1.06E-01 |
| | | A8HPB8 | MSRA4 | Peptidyl-prolyl cis-trans isomerase | 0.69 | 0.45 | 0.38 | 1.26E-02 | 4.24E-03 | 3.33E-03 |
| | | A8IHI1 | PDI4 | Protein disulfide isomerase | 0.93 | 2.49 | 2.33 | 4.36E-01 | 3.08E-03 | 1.02E-02 |
| | | A8JD56 | TIG1 | Chloroplast trigger factor | 0.61 | 0.60 | 0.71 | 2.94E-02 | 5.08E-02 | 8.36E-02 |
| | | A8IZR5 | TRXh | Thioredoxin | 9.04 | 19.31 | 18.47 | 5.35E-02 | 5.12E-02 | 3.58E-02 |
| | | A8HP58 | TRXm | Thioredoxin | 1.05 | 0.60 | 0.35 | 8.14E-01 | 8.48E-02 | 3.57E-02 |
| | | Q84XR9 | TRXx | Thioredoxin x | 4.04 | 5.26 | 5.91 | 3.51E-02 | 3.96E-02 | 7.57E-02 |
| | | A8IQA9 | CHLREDRAFT_11164 | Thioredoxin-like protein | 0.97 | 0.67 | 0.55 | 9.94E-01 | 1.61E-01 | 7.22E-03 |
| | | A8I0I4 | CHLRE_10g456250v5 | Thioredoxin-like protein | 0.58 | 0.36 | 0.32 | 3.72E-02 | 1.27E-02 | 1.38E-02 |
| | | A8J0Q8 | CITRX | Thioredoxin-related protein CITRX | 3.06 | 3.95 | 3.60 | 9.08E-02 | 2.45E-02 | 3.16E-02 |
| | | A8I211 | CHLREDRAFT_161043 | Predicted protein | 1.24 | 1.44 | 1.77 | 1.51E-01 | 4.69E-02 | 1.75E-03 |
| | | A8HUK0 | FKB12 | Peptidyl-prolyl cis-trans isomerase | 0.64 | 0.20 | 0.07 | 3.65E-02 | 3.34E-04 | 6.21E-05 |
| | | A8J746 | FKB15-1 | Peptidyl-prolyl cis-trans isomerase | 0.69 | 0.26 | 0.10 | 4.05E-02 | 5.50E-04 | 8.91E-05 |
| | | A8I6B6 | FKB16-1 | Peptidyl-prolyl cis-trans isomerase | 0.98 | 0.42 | 0.16 | 9.32E-01 | 8.20E-03 | 2.20E-03 |
| | | A8J3L6 | FKB16-2a\|FKB16- [illegible] | Peptidyl-prolyl cis-trans isomerase | 0.85 | 0.43 | 0.20 | 4.01E-01 | 6.75E-03 | 1.03E-02 |
| | | A8JC71 | FKB16-4 | Peptidyl-prolyl cis-trans isomerase | 0.61 | 0.22 | 0.24 | 1.93E-01 | 4.16E-02 | 6.74E-02 |
| | | A8J3L3 | FKB16-5 | Peptidyl-prolyl cis-trans isomerase | 0.85 | 0.57 | 0.34 | 5.35E-01 | 6.94E-02 | 1.74E-02 |
| | | A8J3C1 | FKB16-7a\|FKB16-7b | Peptidyl-prolyl cis-trans isomerase | 0.66 | 0.25 | 0.25 | 1.93E-01 | 1.58E-03 | 1.32E-03 |
| | | A8JEI4 | FKB16-8 | Peptidyl-prolyl cis-trans isomerase | 0.97 | 0.32 | 0.34 | 6.73E-01 | 1.74E-03 | 1.18E-02 |
| | | A8I0A8 | FKB16-9 | Peptidyl-prolyl cis-trans isomerase | 1.19 | 0.67 | 0.45 | 4.22E-01 | 1.15E-01 | 4.98E-02 |
| | | A8JEK6 | FKB17-2 | Peptidyl-prolyl cis-trans isomerase | 0.72 | 0.74 | 0.84 | 1.78E-02 | 1.15E-01 | 4.11E-01 |
| | | A8I1U5 | FKB18 | Peptidyl-prolyl cis-trans isomerase | 0.94 | 0.30 | 0.32 | 5.81E-01 | 2.82E-03 | 2.00E-03 |
| | | A8IVN2 | FKB19 | Peptidyl-prolyl cis-trans isomerase | 4.07 | 1.99 | 1.55 | 1.65E-02 | 1.56E-01 | 3.62E-01 |
| 3 | Carbon metabolism | A8J4D3 | AMYA2 | Alpha-amylase | 2.27 | 3.32 | 1.95 | 4.37E-02 | 3.93E-02 | 5.14E-02 |
| | | A8IZ00 | CHLREDRAFT_173725 | Alpha-amylase-like protein | 5.64 | 2.54 | 2.24 | 4.09E-02 | 1.56E-01 | 3.05E-01 |
| | | A8IZE7 | ATF1 | Glucosamine--fructose-6-phosphate aminotransferase | 0.69 | 2.58 | 2.32 | 3.12E-01 | 1.00E-02 | 4.49E-02 |
| | | A8IEM7 | GHL1 | Glycosyl hydrolase | 1.44 | 3.06 | 2.39 | 9.72E-02 | 1.15E-01 | 3.72E-02 |
| | | A8HPD2 | GPM2 | Phosphoglucomutase | 1.25 | 0.71 | 0.80 | 1.42E-01 | 2.96E-01 | 3.16E-02 |
| | | A8ICG9 | MDH2 | Malate dehydrogenase | 0.37 | 0.94 | 0.34 | 3.21E-02 | 8.11E-01 | 3.41E-02 |
| | | A8J0W9 | MDH3 | Malate dehydrogenase | 1.88 | 3.42 | 3.31 | 2.34E-01 | 4.50E-03 | 1.01E-02 |
| | | A8JHU0 | MDH4 | Malate dehydrogenase | 4.62 | 5.22 | 4.72 | 1.10E-02 | 2.10E-02 | 2.64E-02 |
| | | Q9FNS5 | NADP-mdh | NADP-Malate dehydrogenase | 0.32 | 0.16 | 0.45 | 2.27E-04 | 9.60E-05 | 5.58E-03 |
| | | A8HQW1 | CHLREDRAFT_101528 | 6-phosphogluconolactonase-like protein | 8.14 | 7.27 | 8.51 | 1.14E-04 | 2.73E-03 | 2.98E-04 |
| | | A8HMX2 | PFL1 | Pyruvate-formate lyase | 6.73 | 4.05 | 5.09 | 5.09E-04 | 2.39E-02 | 1.47E-02 |

| | | | | | | | | | | |
|---|---|---|---|---|---|---|---|---|---|---|
| | | A8IYK1 | PHOB | Phosphorylase | 4.07 | 5.96 | 5.03 | 9.69E-02 | 3.26E-02 | 4.11E-02 |
| | | A8IYP4 | PRK1 | Phosphoribulokinase | 0.81 | 0.08 | 0.06 | 3.61E-01 | 2.68E-03 | 2.43E-03 |
| | | A8HQP0 | TAL1 | Transaldolase | 1.32 | 0.61 | 0.75 | 4.53E-01 | 2.95E-02 | 4.71E-02 |
| | | A8IUU3 | TAL2 | Transaldolase | 0.82 | 0.12 | 0.08 | 6.21E-01 | 1.93E-04 | 1.91E-04 |
| | | A8J914 | UGD2 | UDP-glucose dehydrogenase | 0.93 | 1.77 | 1.04 | 8.81E-01 | 2.44E-02 | 9.99E-01 |
| | | A8ITZ0 | CHLRE_07g347100v5 | Predicted protein | 1.11 | 2.11 | 1.79 | 9.35E-01 | 2.73E-02 | 1.66E-01 |
| | | A8J352 | CHLREDRAFT_119219 | Predicted protein | 3.83 | 3.70 | 4.64 | 1.16E-02 | 4.62E-03 | 5.03E-03 |
| | | A8ICN2 | CHLREDRAFT_96789 | Predicted protein | 3.88 | 6.13 | 6.10 | 3.71E-01 | 1.99E-02 | 2.77E-02 |
| | | A8J3Q5 | CHLREDRAFT_80327 | Predicted protein | 0.59 | 0.51 | 0.34 | 3.03E-02 | 2.04E-02 | 8.86E-03 |
| | | A8IXP6 | CHLRE_03g200650v5 | Predicted protein | 1.63 | 1.00 | 1.12 | 2.78E-02 | 9.63E-01 | 6.59E-01 |
| | | A8HMQ1 | ACH1 | Aconitate hydratase | 1.79 | 3.77 | 3.24 | 3.56E-01 | 1.36E-02 | 8.69E-03 |
| | | A8JHC9 | CIS1 | Citrate synthase | 12.56 | 16.14 | 15.51 | 6.88E-03 | 1.31E-03 | 1.02E-02 |
| | | A8J2S0 | CIS2 | Citrate synthase | 0.54 | 0.42 | 0.27 | 5.60E-02 | 2.04E-02 | 5.23E-03 |
| | | A8HX04 | SDH2 | Iron-sulfur subunit of mitochondrial succinate dehydrogenase | 23.15 | 37.75 | 34.39 | 1.02E-01 | 9.14E-03 | 5.02E-02 |
| | | A8J9S7 | IDH3 | Isocitrate dehydrogenase [NADP] | 5.16 | 7.19 | 5.65 | 1.50E-02 | 6.04E-04 | 4.02E-03 |
| | | A8J6V1 | IDH1 | Isocitrate dehydrogenase NAD-dependent | 1.19 | 1.61 | 1.74 | 4.73E-01 | 1.31E-01 | 3.97E-02 |
| | | A8ICG9 | MDH2 | Malate dehydrogenase | 0.37 | 0.94 | 0.34 | 3.21E-02 | 8.11E-01 | 3.41E-02 |
| | | A8J0W9 | MDH3 | Malate dehydrogenase | 1.88 | 3.42 | 3.31 | 2.34E-01 | 4.50E-03 | 1.01E-02 |
| | | A8JHU0 | MDH4 | Malate dehydrogenase | 4.62 | 5.22 | 4.72 | 1.10E-02 | 2.10E-02 | 2.64E-02 |
| | | Q6X898 | MAS1 | Malate synthase | 0.58 | 0.14 | 0.10 | 8.07E-02 | 1.28E-02 | 1.15E-02 |
| | | Q9FNS5 | NADP-mdh | NADP-malate dehydrogenase | 0.32 | 0.16 | 0.45 | 2.27E-04 | 9.60E-05 | 5.58E-03 |
| | | A8HP06 | SDH1 | Succinate dehydrogenase subunit A | 1.70 | 2.45 | 1.81 | 1.83E-01 | 2.71E-02 | 1.02E-01 |
| | | A8HPU1 | SDH4 | Succinate dehydrogenase subunit D | 1.02 | 0.68 | 0.41 | 8.02E-01 | 1.16E-01 | 3.72E-02 |
| | | A8J244 | ICL1 | Isocitrate lyase | 0.78 | 0.24 | 0.11 | 6.10E-01 | 6.01E-02 | 3.41E-02 |
| | | A8IXQ5 | CHLRE_03g200250v5 | Predicted protein | 0.64 | 0.14 | 0.19 | 3.82E-02 | 1.36E-03 | 1.45E-03 |
| | | A8J2R7 | CHLREDRAFT_104431 | Predicted protein (Fragment) | 2.03 | 5.22 | 7.63 | 1.99E-01 | 2.51E-02 | 8.56E-03 |
| | | A8IKQ0 | FBP1 | Fructose-1,6-bisphosphatase | 1.11 | 0.52 | 0.35 | 6.64E-01 | 1.86E-02 | 6.54E-03 |
| | | A8IE23 | PGI1 | Glucose-6-phosphate isomerase | 4.77 | 6.51 | 5.53 | 6.61E-03 | 8.84E-03 | 1.64E-02 |
| | | A8J0N7 | PCK1b\|PCK1a | Phosphoenolpyruvate carboxykinase splice variant | 1.28 | 0.46 | 0.33 | 4.80E-01 | 7.80E-02 | 2.54E-02 |
| | | A8JCE1 | CHLREDRAFT_121619 | Predicted protein (Fragment) | 0.14 | 0.13 | 0.19 | 1.88E-02 | 1.70E-02 | 2.24E-02 |
| | | A8IRK4 | SEBP1 | Sedoheptulose-1,7-bisphosphatase | 0.42 | 0.13 | 0.12 | 3.94E-02 | 1.07E-02 | 1.37E-02 |
| | | A8HX70 | PFK1 | Phosphofructokinase family protein | 3.05 | 3.05 | 2.55 | 5.83E-03 | 3.03E-04 | 1.02E-01 |
| | | A8IYM0 | PFK2 | Phosphofructokinase family protein | 2.01 | 3.14 | 3.29 | 2.08E-03 | 7.28E-04 | 1.35E-03 |
| 4 | Photosynthesis | P22666 | psbH | Photosystem II reaction center protein H | 30.03 | 25.79 | 19.68 | 4.75E-02 | 1.59E-02 | 1.09E-03 |
| | | A8J0E4 | PsbO | Oxygen-evolving enhancer protein 1 of photosystem II | 0.99 | 0.82 | 0.78 | 8.87E-01 | 6.22E-05 | 2.81E-05 |
| | | A8IXU9 | CGL30/PsbP | Photosystem II thylakoid lumenal 29.8 kDa protein PsbP | 16.11 | 14.48 | 16.24 | 5.36E-02 | 1.46E-02 | 3.91E-02 |
| | | A8J3S9 | PsbP2 | PsbP-like protein | 1.01 | 0.46 | 0.54 | 8.96E-01 | 8.70E-02 | 2.53E-02 |
| | | A8JEV1 | PsbQ | Oxygen evolving enhancer protein 3 | 21.52 | 27.63 | 25.07 | 7.66E-02 | 1.15E-03 | 2.14E-02 |

| | | | | | | | | | | |
|---|---|---|---|---|---|---|---|---|---|---|
| | | A8JFQ7 | PsbW | Photosystem II reaction center W protein | 0.21 | 0.07 | 0.09 | 3.68E-02 | 1.82E-02 | 1.98E-02 |
| | | A8III5 | Psb28 | Photosystem II reaction center psb28 protein | 0.71 | 0.18 | 0.13 | 1.65E-01 | 1.48E-02 | 1.24E-02 |
| | | P12154 | psaA | Photosystem I P700 chlorophyll a apoprotein A1 | 0.89 | 4.42 | 4.04 | 4.87E-01 | 2.83E-02 | 3.94E-02 |
| | | P09144 | psaB | Photosystem I P700 chlorophyll a apoprotein A2 | 1.94 | 11.43 | 8.18 | 7.64E-01 | 1.34E-02 | 1.04E-01 |
| | | Q5NKW4 | PsaD | Photosystem I reaction center subunit II 20 kDa | 13.21 | 20.61 | 19.15 | 3.30E-02 | 2.01E-03 | 3.28E-03 |
| | | A8J4S1 | PsaF | Photosystem I reaction center subunit III | 7.69 | 1.36 | 1.71 | 4.46E-02 | 6.01E-01 | 5.47E-01 |
| | | A8JHN9 | PsaG | Photosystem I reaction center subunit V | 0.42 | 0.06 | 0.05 | 5.62E-04 | 4.85E-06 | 8.48E-06 |
| | | A8J6K8 | PsaK | Photosystem I reaction center subunit psaK | 0.77 | 0.29 | 0.27 | 1.24E-01 | 3.78E-03 | 5.12E-03 |
| | | A8IL32 | PsaL | Photosystem I reaction center subunit XI | 0.87 | 0.34 | 0.49 | 7.35E-01 | 1.38E-02 | 2.82E-02 |
| | | A8I835 | PsaN | Photosystem I reaction center subunit N | 0.82 | 0.05 | 0.05 | 3.29E-01 | 2.39E-05 | 3.02E-05 |
| | | O20030 | ycf4 | Photosystem I assembly protein Ycf4 | 0.28 | 0.25 | 0.20 | 1.14E-04 | 6.28E-04 | 1.77E-04 |
| | | P23577 | petA | Apocytochrome f | 2.90 | 0.33 | 0.22 | 2.11E-02 | 1.06E-01 | 7.28E-02 |
| | | Q00471 | petB | Cytochrome b6 | 0.57 | 0.24 | 0.31 | 1.30E-01 | 2.59E-02 | 2.86E-02 |
| | | A8HQJ5 | TEF30 | Predicted protein | 1.17 | 0.42 | 0.13 | 9.81E-01 | 1.05E-01 | 3.12E-02 |
| | | A8I531 | CHLD | Magnesium chelatase subunit D | 0.18 | 0.14 | 0.19 | 7.89E-04 | 4.69E-04 | 6.84E-04 |
| | | A8IMZ5 | CHLI1 | Magnesium chelatase subunit I | 0.13 | 0.08 | 0.15 | 4.50E-04 | 2.82E-04 | 4.74E-04 |
| | | A8IKQ6 | CHLI2 | Magnesium chelatase subunit I | 0.26 | 0.11 | 0.25 | 2.07E-03 | 1.02E-03 | 3.65E-03 |
| | | A8HPJ2 | POR | Light-dependent protochlorophyllide reductase | 0.52 | 0.06 | 0.14 | 4.33E-02 | 5.12E-03 | 7.21E-03 |
| | | Q93WL4 | LHCBM3 | Light-harvesting chlorophyll-a/b binding protein LhcII-1.3 | 2.33 | 0.73 | 0.38 | 5.49E-02 | 2.12E-01 | 3.91E-02 |
| | | Q9ZSJ4 | LHCBM5 | Chlorophyll a-b binding protein of LHCII | 12.89 | 4.40 | 2.80 | 1.08E-02 | 7.89E-02 | 3.48E-02 |
| | | A8J287 | LHCBM6 | Chloropyll a-b binding protein of LHCII type I chloroplast | 0.97 | 0.35 | 0.13 | 8.54E-01 | 2.03E-02 | 5.18E-03 |
| | | A8J270 | LHCBM8 | Chlorophyll a-b binding protein of LHCII | 0.78 | 0.29 | 0.25 | 6.71E-01 | 5.77E-02 | 4.29E-02 |
| | | Q8S3T9 | LHCBM9 | Chlorophyll a-b binding protein of LHCII | 36.48 | 27.70 | 26.74 | 5.89E-03 | 1.47E-03 | 5.01E-03 |
| | | A8J249 | LHCA1 | Light-harvesting protein of photosystem I | 2.32 | 0.73 | 0.29 | 2.43E-01 | 2.66E-01 | 2.20E-03 |
| | | A8IKC8 | LHCA2 | Light-harvesting protein of photosystem I | 1.52 | 0.21 | 0.80 | 2.70E-01 | 7.76E-03 | 2.33E-01 |
| | | Q75VY8 | LHCA5 | Light-harvesting chlorophyll-a/b protein of photosystem I | 0.78 | 0.29 | 0.40 | 3.76E-01 | 5.76E-05 | 4.28E-04 |
| | | Q75VY6 | LHCA6 | Light-harvesting chlorophyll-a/b protein of photosystem I | 0.90 | 0.25 | 0.35 | 5.06E-01 | 6.28E-03 | 2.85E-02 |
| | | A8ISG0 | LHCA7 | Light-harvesting protein of photosystem I | 0.87 | 0.23 | 0.21 | 4.58E-01 | 4.93E-03 | 3.44E-03 |
| | | A8ITV3 | LHCA9 | Light-harvesting protein of photosystem I | 0.75 | 0.13 | 0.13 | 2.34E-01 | 8.53E-05 | 1.51E-04 |
| | | A8J0A7 | ELI3 | Early light-inducible protein | 1.34 | 2.21 | 2.22 | 2.45E-01 | 3.93E-02 | 1.32E-01 |
| 5 | Redox homeostasis | A8HNQ7 | NTRC1 | Thioredoxin reductase | 0.49 | 0.34 | 0.36 | 1.71E-03 | 1.50E-04 | 2.40E-03 |
| | | A8IUG8 | CDSP32 | Plastidic thioredoxin-like protein | 1.02 | 0.26 | 0.42 | 9.20E-01 | 9.08E-03 | 2.25E-02 |
| | | Q9FE86 | PRX1 | 2-cys peroxiredoxin chloroplastic | 16.83 | 17.62 | 16.67 | 1.14E-03 | 9.71E-05 | 7.07E-04 |
| | | A8J0Q8 | CITRX | Thioredoxin-related protein CITRX | 3.06 | 3.95 | 3.60 | 9.08E-02 | 2.45E-02 | 3.16E-02 |
| | | A8JDC5 | DLC5 | Flagellar outer arm dynein 14 kDa light chain LC5 | 0.44 | 0.40 | 0.44 | 4.73E-02 | 8.59E-03 | 2.69E-02 |
| | | A8HPL8 | DLD2 | Dihydrolipoamide dehydrogenase | 0.72 | 0.31 | 0.20 | 2.82E-01 | 2.62E-02 | 2.06E-02 |
| | | A8J6A7 | MET16/APR1 | Adenylylphosphosulfate reductase | 8.56 | 14.80 | 15.71 | 1.84E-03 | 1.70E-04 | 8.61E-04 |
| | | A8J1T4 | GCSL | Dihydrolipoyl dehydrogenase | 2.95 | 5.47 | 5.00 | 1.98E-01 | 5.53E-02 | 4.37E-02 |

| | | | | | | | | | | |
|---|---|---|---|---|---|---|---|---|---|---|
| | | A8JHA9 | GRX1 | Glutaredoxin CPYC type | 2.76 | 3.31 | 3.57 | 1.98E-02 | 2.71E-04 | 1.06E-04 |
| | | A8JH05 | GRX3 | Glutaredoxin CGFS type | 0.73 | 0.45 | 0.48 | 4.94E-02 | 1.18E-02 | 1.09E-02 |
| | | A8HN52 | GRX6 | Glutaredoxin CGFS type | 0.46 | 0.22 | 0.24 | 3.50E-02 | 1.27E-03 | 3.54E-03 |
| | | A8J0E5 | GSHR2 | Glutathione reductase | 1.24 | 2.16 | 2.08 | 2.91E-01 | 6.69E-03 | 2.01E-02 |
| | | A8J448 | NTR1 | NADPH-dependent thioredoxin reductase 1 | 0.98 | 1.64 | 1.22 | 9.47E-01 | 4.73E-02 | 6.37E-01 |
| | | A8HQT1 | PDI2 | Protein disulfide isomerase | 10.61 | 10.09 | 8.47 | 4.38E-03 | 1.98E-02 | 3.96E-02 |
| | | A8IHI1 | PDI4 | Protein disulfide isomerase | 0.93 | 2.49 | 2.33 | 4.36E-01 | 3.08E-03 | 1.02E-02 |
| | | A8I4C2 | PDI5 | Protein disulfide isomerase | 0.65 | 0.58 | 0.34 | 1.11E-02 | 1.76E-01 | 1.64E-02 |
| | | A8HZQ4 | PRX3 | Peroxiredoxin type II | 0.83 | 0.57 | 0.36 | 1.69E-01 | 9.36E-03 | 1.34E-03 |
| | | A8JFC3 | SCO1 | Cytochrome c oxidase assembly factor | 3.13 | 4.09 | 4.95 | 4.26E-02 | 1.62E-02 | 2.05E-02 |
| | | A8IZR5 | TRXh | Thioredoxin | 9.04 | 19.31 | 18.47 | 5.35E-02 | 5.12E-02 | 3.58E-02 |
| | | A8HP58 | TRXm | Thioredoxin | 1.05 | 0.60 | 0.35 | 8.14E-01 | 8.48E-02 | 3.57E-02 |
| | | Q84XR9 | TRXx | Thioredoxin x | 4.04 | 5.26 | 5.91 | 3.51E-02 | 3.96E-02 | 7.57E-02 |
| | | A8IXH4 | CHLREDRAFT_205510 | EF-Hand domain-containing thioredoxin | 6.82 | 6.24 | 5.33 | 3.63E-02 | 3.42E-02 | 1.77E-02 |
| | | A8I0I4 | CHLRE_10g456250v5 | Thioredoxin-like protein | 0.58 | 0.36 | 0.32 | 3.72E-02 | 1.27E-02 | 1.38E-02 |
| | | A8IQA9 | CHLREDRAFT_11164 | Thioredoxin-like protein | 0.97 | 0.67 | 0.55 | 9.94E-01 | 1.61E-01 | 7.22E-03 |
| | | A8JG35 | CHLREDRAFT_160132 | Predicted protein | 0.95 | 0.52 | 0.48 | 5.06E-01 | 5.21E-02 | 3.39E-02 |
| | | A8JDA2 | CHLRE_17g715500v5 | Predicted protein | 2.43 | 1.31 | 1.08 | 2.89E-03 | 3.49E-01 | 9.41E-01 |
| | | A8J0Q4 | CHLREDRAFT_191208 | Predicted protein | 2.38 | 4.20 | 3.65 | 4.62E-01 | 4.41E-02 | 1.22E-01 |
| | | A8JA70 | CHLRE_16g687294v5 | Ferredoxin thioredoxin reductase variable chain | 1.98 | 0.68 | 0.33 | 2.49E-02 | 1.18E-01 | 1.94E-02 |
| | | O49822 | apx1 | Ascorbate peroxidase | 7.53 | 11.12 | 13.11 | 1.21E-02 | 3.31E-02 | 7.36E-03 |
| | | A8J7X9 | CCPR1 | Cytochrome c peroxidase | 12.06 | 12.80 | 10.92 | 5.09E-05 | 5.23E-03 | 7.00E-03 |
| | | A8J285 | APX2 | L-ascorbate peroxidase | 3.09 | 3.46 | 4.49 | 3.05E-02 | 4.10E-02 | 4.71E-02 |
| | | O81648 | Lci2 | Low CO2 inducible gene | 0.79 | 0.07 | 0.16 | 3.32E-01 | 1.87E-04 | 6.86E-04 |
| | | A8J537 | CAT1 | Catalase | 3.13 | 4.92 | 4.37 | 5.83E-02 | 1.17E-03 | 4.88E-05 |
| | | A8IXD6 | CLPR4 | ATP-dependent Clp protease proteolytic subunit | 6.84 | 5.82 | 5.04 | 2.06E-02 | 4.72E-02 | 1.29E-01 |
| 6 | Intracellular protein trafficking | A8I1Y7 | SNAPA1 | Alpha-SNAP | 0.94 | 3.31 | 2.91 | 9.51E-01 | 3.73E-02 | 8.47E-02 |
| | | A8I4S9 | CHC1 | Clathrin heavy chain | 2.60 | 16.32 | 14.00 | 6.11E-01 | 1.47E-02 | 1.78E-02 |
| | | A8HQF0 | VPS35 | Subunit of retromer complex | 2.77 | 3.49 | 3.31 | 1.04E-02 | 3.36E-03 | 9.94E-04 |
| | | A8IM71 | COPG1 | Coatomer subunit gamma | 1.22 | 4.00 | 2.82 | 8.46E-01 | 4.12E-02 | 6.82E-02 |
| | | A8J4X1 | VPS26 | Subunit of retromer complex | 6.42 | 4.86 | 4.91 | 6.47E-04 | 1.25E-02 | 1.16E-03 |
| | | A8INI9 | CHLREDRAFT_170116 | Predicted protein | 1.36 | 0.86 | 1.11 | 3.53E-02 | 2.79E-01 | 7.56E-01 |
| | | A8JEP9 | COPB1 | Coatomer subunit beta | 1.01 | 3.93 | 3.32 | 9.99E-01 | 3.69E-02 | 5.48E-02 |
| | | A8I211 | CHLREDRAFT_161043 | Predicted protein | 1.24 | 1.44 | 1.77 | 1.51E-01 | 4.69E-02 | 1.75E-03 |
| | | Q8S4W5 | SYP6 | Qc-SNARE protein Tlg1/Syntaxin 6-family | 2.82 | 2.89 | 2.98 | 4.07E-02 | 1.65E-02 | 4.07E-03 |
| | | A8J729 | AP1B1 | Putative uncharacterized protein AP1B1 | 2.70 | 5.15 | 4.06 | 5.51E-02 | 2.56E-04 | 1.34E-03 |
| | | A8IR64 | SYP5 | Qc-SNARE protein Syn8/Syntaxin8-family | 4.09 | 5.61 | 5.62 | 6.63E-02 | 8.15E-02 | 4.75E-02 |
| | | A8JBG7 | SYP3 | Qa-SNARE protein Sed5/Syntaxin5-family | 1.58 | 1.16 | 0.71 | 9.51E-02 | 6.30E-01 | 3.55E-02 |

| | | | | | | | | | | |
|---|---|---|---|---|---|---|---|---|---|---|
| | | A8JC30 | SAR1 | Sar-type small GTPase | 3.05 | 4.19 | 3.02 | 7.23E-02 | 2.70E-02 | 5.48E-02 |
| | | A8JGS8 | COPB2 | Beta'-cop | 1.56 | 3.25 | 1.91 | 7.90E-02 | 4.00E-03 | 1.49E-02 |
| | | A8IMC5 | AP4E1 | Epsilon-adaptin | 0.83 | 2.44 | 2.23 | 2.81E-01 | 2.20E-02 | 1.18E-02 |
| | | A8J7K1 | CLC1 | Clathrin light chain | 2.90 | 2.62 | 1.96 | 2.37E-02 | 7.70E-02 | 1.31E-01 |
| | | A8JCA4 | CHLREDRAFT_195581 | RabGAP/TBC protein | 5.94 | 12.65 | 9.92 | 3.04E-02 | 3.90E-03 | 1.97E-03 |
| | | A8IYJ6 | CGL38 | Predicted protein | 1.47 | 0.20 | 0.21 | 5.06E-01 | 1.81E-02 | 1.01E-02 |
| | | A8HXA2 | AP1M1 | Mu1-Adaptin | 1.89 | 2.95 | 2.66 | 2.80E-01 | 5.20E-03 | 2.16E-03 |
| | | A8IL38 | CHLREDRAFT_128005 | RabGAP/TBC protein | 1.37 | 1.70 | 1.85 | 3.65E-01 | 4.21E-02 | 1.15E-01 |
| | | A8ILF4 | AP2A1 | Alpha-adaptin | 2.46 | 7.45 | 5.88 | 3.94E-02 | 3.50E-03 | 4.37E-02 |
| | | A8HRR9 | COPA1 | Alpha-COP | 1.61 | 6.81 | 5.72 | 5.33E-01 | 5.06E-03 | 1.57E-02 |
| | | A8IAJ1 | VPS4 | AAA-ATPase of VPS4/SKD1 family | 1.15 | 1.00 | 0.53 | 5.02E-01 | 7.83E-01 | 3.65E-02 |
| | | A8HSQ2 | FAP66 | Flagellar associated protein | 0.66 | 2.61 | 2.85 | 1.30E-01 | 3.44E-02 | 5.30E-02 |
| | | A8IYH0 | CHLRE_12g549950v5 | Tetraspanning membrane protein SFT2-like protein | 0.69 | 0.21 | 0.12 | 7.70E-02 | 5.06E-03 | 1.07E-03 |
| | | A8I023 | TIC110 | 110 kDa translocon of chloroplast envelope inner membrane (Fragment) | 0.42 | 0.39 | 0.44 | 9.45E-03 | 1.19E-02 | 2.15E-02 |
| | | Q6Y682 | Rap38 | 38 kDa ribosome-associated protein | 1.16 | 0.37 | 0.10 | 1.38E-01 | 7.37E-05 | 6.20E-05 |
| | | A8IA39 | EFG1 | Chloroplast elongation factor G | 0.42 | 0.48 | 0.52 | 1.49E-02 | 2.51E-03 | 4.08E-02 |
| | | A8J680 | SECA2 | Chloroplast-associated SecA protein (Fragment) | 0.54 | 0.75 | 0.65 | 9.55E-03 | 6.62E-02 | 2.94E-02 |
| | | A8JBT7 | EGY1 | Membrane associated metalloprotease | 0.79 | 0.39 | 0.30 | 2.89E-01 | 9.43E-02 | 2.70E-02 |
| | | A8HWL8 | CHLREDRAFT_142189 | Predicted protein | 1.56 | 2.54 | 2.27 | 1.46E-01 | 2.35E-02 | 1.56E-02 |
| | | A8J209 | CGL59 | Predicted protein | 0.57 | 0.53 | 0.33 | 9.96E-02 | 6.71E-02 | 3.24E-03 |
| | | A8J682 | SECA1 | Protein translocase subunit SecA | 0.45 | 0.69 | 0.83 | 1.09E-02 | 6.89E-02 | 3.79E-01 |
| | | Q5S7Y5 | TIM | Triosephosphate isomerase | 2.68 | 4.40 | 3.56 | 2.30E-01 | 4.89E-02 | 8.18E-02 |
| 7 | N- and S-metabolism | A8J6Q7 | SHKA1 | 3-deoxy-D-arabino-heptulosonate 7-phosphate synthetase | 0.31 | 0.62 | 0.57 | 3.50E-02 | 1.41E-01 | 1.98E-01 |
| | | A8JH48 | SHKG1 | 3-phosphoshikimate 1-carboxyvinyltransferase | 0.36 | 0.43 | 0.47 | 1.15E-03 | 2.78E-03 | 2.62E-03 |
| | | A8J434 | OASTL1 | Cysteine synthase | 0.58 | 0.45 | 0.54 | 1.93E-02 | 7.21E-03 | 2.25E-02 |
| | | A8IEE5 | OASTL3 | Cysteine synthase | 5.41 | 4.25 | 4.99 | 7.89E-02 | 6.16E-02 | 2.19E-02 |
| | | A8ISA9 | OASTL4 | Cysteine synthase | 17.24 | 18.20 | 17.81 | 3.38E-02 | 2.18E-02 | 3.53E-02 |
| | | A8JG03 | LEU1L | Isopropylmalate dehydratase large subunit | 1.97 | 6.80 | 8.04 | 1.58E-01 | 2.47E-03 | 2.14E-03 |
| | | A8I9R1 | THD1 | Threonine deaminase | 0.42 | 0.40 | 0.50 | 5.07E-03 | 6.49E-03 | 3.53E-02 |
| | | A8IFZ9 | MAA7 | Tryptophan synthase beta subunit | 0.60 | 0.81 | 0.58 | 1.45E-02 | 4.77E-02 | 8.29E-02 |
| | | A8J129 | AST1 | Aspartate aminotransferase | 0.59 | 0.62 | 0.33 | 1.95E-01 | 2.09E-01 | 4.36E-02 |
| | | A8I263 | AST3 | Aspartate aminotransferase | 2.01 | 2.14 | 2.32 | 1.04E-01 | 2.76E-04 | 3.30E-02 |
| | | A8IAK9 | PYR2 | Aspartate carbamoyltransferase | 0.43 | 0.26 | 0.33 | 1.36E-01 | 4.94E-02 | 8.04E-02 |
| | | Q7XXT3 | gdh | Glutamate dehydrogenase | 8.04 | 14.61 | 17.62 | 5.11E-02 | 3.34E-02 | 1.01E-02 |
| | | A8JGD1 | GDH2 | Glutamate dehydrogenase | 14.42 | 27.91 | 31.36 | 4.37E-04 | 2.15E-04 | 6.02E-04 |
| | | A8J3Q6 | APK1 | Adenylyl-sulfate kinase | 3.70 | 4.53 | 3.82 | 7.24E-03 | 1.60E-02 | 2.31E-04 |
| | | A8IXF1 | ATS1 | ATP-sulfurylase | 2.00 | 6.83 | 7.68 | 3.94E-01 | 1.95E-02 | 2.85E-02 |
| | | A8I3V3 | ATS2 | ATP-sulfurylase | 14.22 | 33.32 | 36.36 | 2.98E-02 | 7.02E-03 | 6.98E-03 |

| | | | | | | | | | | |
|---|---|---|---|---|---|---|---|---|---|---|
| | | A8JFW4 | DXS1 | 1-deoxy-D-xylulose 5-phosphate synthase (Fragment) | 0.31 | 0.20 | 0.19 | 2.71E-03 | 1.03E-03 | 1.53E-03 |
| | | A8ILN4 | HDS1 | 1-hydroxy-2-methyl-2-(E)-butenyl 4-diphosphate synthase | 0.54 | 1.12 | 1.22 | 3.99E-03 | 5.88E-01 | 3.36E-01 |
| | | A8IJL3 | CHLRE_12g503550v5 | 2-C-methyl-D-erythritol 24-cyclodiphosphate synthase | 3.88 | 6.20 | 5.70 | 9.18E-03 | 1.95E-03 | 1.45E-02 |
| | | A8J0V1 | CMK1 | 4-diphosphocytidyl-2-C-methyl-D-erythritol kinase | 0.87 | 0.36 | 0.24 | 4.94E-01 | 2.18E-02 | 1.52E-02 |
| | | A8IMN5 | CMPS1 | Carbamoyl phosphate synthase small subunit | 1.46 | 1.43 | 1.33 | 1.61E-01 | 2.38E-02 | 6.28E-02 |
| | | A8JBN5 | PYR4 | Dihydropryrimidine dehydrogenase | 1.23 | 3.39 | 3.95 | 8.19E-01 | 5.44E-02 | 2.14E-02 |
| | | A8HQF3 | FAK1 | ODA5-associated flagellar adenylate kinase | 0.48 | 1.18 | 0.72 | 3.12E-03 | 7.21E-01 | 3.18E-01 |
| | | A8HN92 | PYR5 | Uridine 5'-monophosphate synthase | 1.35 | 2.97 | 2.19 | 5.01E-01 | 1.33E-02 | 5.98E-03 |
| 8 | ATP hydrolysis coupled proton transport | Q96550 | atpA | ATP synthase subunit alpha | 3.23 | 3.60 | 2.66 | 3.79E-02 | 3.56E-02 | 1.10E-01 |
| | | A8HX15 | CHLREDRAFT_187139 | Sodium/potassium-transporting ATPase alpha subunit | 5.75 | 7.33 | 7.63 | 4.32E-02 | 3.30E-04 | 1.70E-03 |
| | | A8I164 | ATPvA1 | Vacuolar ATP synthase subunit A | 2.48 | 5.57 | 4.19 | 4.17E-01 | 1.72E-02 | 4.95E-02 |
| | | A8IA45 | ATPvB | Vacuolar ATP synthase subunit B | 3.16 | 2.97 | 1.59 | 2.29E-02 | 4.28E-02 | 3.25E-01 |
| | | A8IW47 | ATPvE | Vacuolar ATP synthase subunit E | 2.52 | 2.85 | 1.99 | 8.14E-03 | 7.69E-03 | 3.96E-02 |
| | | A8HQ97 | ATPvH | Vacuolar ATP synthase subunit H | 3.17 | 8.94 | 7.24 | 1.56E-01 | 3.52E-02 | 2.97E-02 |
| | | A8IDY0 | ATPvD1 | Vacuolar H+ ATPase V0 sector subunit D | 2.38 | 4.44 | 2.60 | 1.03E-01 | 3.68E-02 | 7.71E-03 |
| | | A8HYU2 | ATPvC | Vacuolar H+ ATPase V1 sector subunit C | 2.54 | 4.82 | 3.38 | 1.87E-01 | 4.76E-02 | 7.18E-03 |
| | | A8IST3 | ATPvA3 | Vacuolar proton ATPase subunit A | 1.18 | 3.69 | 3.38 | 5.66E-01 | 5.36E-02 | 3.82E-02 |
| | | A8J1K0 | ATPvA2 | Vacuolar proton translocating ATPase subunit A | 4.24 | 7.42 | 5.63 | 7.28E-03 | 3.36E-04 | 2.32E-03 |
| 9 | Response to cytokinin | A8J3F8 | CHLRE_16g672750v5 | Predicted protein | 0.70 | 0.24 | 0.19 | 7.35E-02 | 4.33E-03 | 4.27E-03 |
| | | Q70DX8 | S1 | Plastid ribosomal protein S1 | 0.33 | 0.08 | 0.07 | 2.00E-02 | 3.55E-03 | 3.42E-03 |
| | | A8JD33 | CHLREDRAFT_153076 | Predicted protein | 1.74 | 2.44 | 1.98 | 1.76E-01 | 1.58E-02 | 1.21E-01 |
| | | A8IWA6 | GSN1 | Glutamate synthase NADH-dependent | 1.60 | 5.62 | 4.40 | 7.26E-01 | 2.23E-02 | 4.60E-03 |
| | | A8HWZ6 | PRPL13 | Plastid ribosomal protein L13 | 0.43 | 0.12 | 0.15 | 3.68E-02 | 1.08E-02 | 1.08E-02 |
| | | A8IU62 | PIN3 | Peptidyl-prolyl cis-trans isomerase parvulin-type | 0.74 | 0.50 | 0.45 | 9.70E-02 | 1.24E-02 | 4.89E-02 |
| | | A8HVJ9 | CHLREDRAFT_112806 | Photosystem II stability/assembly factor HCF136 | 0.24 | 0.09 | 0.10 | 1.33E-03 | 5.74E-04 | 5.72E-04 |
| | | A8J503 | PRPL6 | Plastid ribosomal protein L6 | 0.79 | 0.20 | 0.15 | 1.58E-01 | 3.33E-03 | 2.03E-03 |
| | | Q8HTL1 | rpl5 | 50S ribosomal protein L5 chloroplastic | 1.33 | 0.64 | 0.52 | 2.64E-01 | 6.14E-03 | 5.84E-04 |
| | | A8JAL6 | PRPL15 | Plastid ribosomal protein L15 | 0.33 | 0.05 | 0.05 | 2.04E-02 | 5.06E-03 | 5.00E-03 |
| | | A8J785 | ATPG | ATP synthase subunit b' chloroplastic | 0.61 | 0.26 | 0.33 | 1.51E-01 | 4.60E-03 | 3.82E-02 |
| | | A8IYP4 | PRK1 | Phosphoribulokinase | 0.81 | 0.08 | 0.06 | 3.61E-01 | 2.68E-03 | 2.43E-03 |
| | | A8J8U1 | RPN10 | 26S proteasome regulatory subunit | 0.84 | 0.68 | 0.57 | 8.64E-02 | 7.18E-03 | 1.47E-02 |
| | | A8I8X2 | DEG1A | DegP-type protease | 2.45 | 1.38 | 1.01 | 3.97E-03 | 3.79E-01 | 9.50E-01 |
| | | A8IKQ0 | FBP1 | Fructose-1,6-bisphosphatase | 1.11 | 0.52 | 0.35 | 6.64E-01 | 1.86E-02 | 6.54E-03 |
| | | A8INR7 | PRPL27 | Plastid ribosomal protein L27 | 0.19 | 0.05 | 0.04 | 2.00E-03 | 8.47E-04 | 7.99E-04 |

**Table S4.** List of primers used in this work.

| Primers | Nucleotide sequences (5’ to 3’) |
|---|---|
| *LHCA2*-F | GAAGAGCGAGGAGATGAAGC |
| *LHCA2*-R | CCGACGAGGTGTAGATGTTG |
| *LHCA5*-F | AGACCAAGGAGATCAAGAACGG |
| *LHCA5*-R | TGCAAGTGCCGATGTTCTTC |
| *LHCA6*-F | TGATGCTGGTTGCCAAGAAC |
| *LHCA6*-R | TTGAACCACTTCAGGGACTCG |
| *LHCA7*-F | TCATCCTCACCTCCATTGGT |
| *LHCA7*-R | TTCTTGAAGTCGTACCAGCG |
| *LHCA9*-F | TTCATCAACTCCTTCCCCTTCG |
| *LHCA9*-R | AGGTGATGTTCTTGCCGAAG |
| *CBLP*-F | ATGACCACCAACCCCATCATC |
| *CBLP*-R | GGTCCCACAGCATGGCAATG |

**Legends for Movie S1 and Datasets**

**Movie S1 (**separate file**).** Demonstration of a laboratory set-up making a $H_2$-fuel-cell-powered toycar (MS812-E-1; Ningbo MS, China) drive by input of algal-$H_2$ collected from *hpm91* and ambient air.

**Dataset S1** (separate file)**.** List of confidentially identified proteins with quantitative information.

**Dataset S2** (separate file)**.** List of proteins corresponding to group I of Fig. 3C.

**Dataset S3** (separate file)**.** List of proteins corresponding to group II of Fig. 3C.

**Dataset S4** (separate file)**.** List of proteins corresponding to group III of Fig. 3C.

**Dataset S5** (separate file)**.** GOBP enrichment results derived from Dataset S3.

**Dataset S6** (separate file)**.** GOBP enrichment results derived from Dataset S4.
